# Fidelity in co-diversified symbiosis

**DOI:** 10.64898/2025.12.23.696161

**Authors:** Inès Pons, Marleny García-Lozano, Christiane Emmerich, Aftab Mahmood Ayas, Christine Henzler, Hagay Enav, Ruth E. Ley, Hassan Salem

## Abstract

Obligate co-dependence can arise in symbiosis, yielding heritable partnerships. These interactions are considered to be highly specific, but partner fidelity is difficult to quantify owing to the experimental constraints of symbiont exchange between host species. Here, we overcome this challenge by leveraging the unique transmission dynamics of *Stammera capleta*, the obligate digestive symbiont of tortoise beetles (Chrysomelidae: Cassidinae). Despite its extracellular localization, *S. capleta* possesses a drastically reduced genome (∼0.25 Mb) and is vertically transmitted through egg-associated spheres. Manipulating these spheres allowed us to experimentally exchange *S. capleta* between beetle species to determine their impact on host development. We show that non-native *S. capleta* can successfully colonize the symbiotic organs of a novel host, but that the interaction outcome correlates with genetic relatedness to the native symbiont. Genetically distant species trigger a more pronounced transcriptional response and can only partially rescue host development. While more closely related symbionts proliferate similarly to the native one and induce a comparable host response, they fail to propagate to the next generation, underscoring how transmission fidelity, host-symbiont compatibility, and local adaptation can further specificity within a Paleocene-aged partnership.

## Introduction

The metabolic intimacy of symbiosis demands the work of specialists, each evolving distinct yet complementary roles to sustain the partnership ^1,2^. Obligate co-dependence can arise within these interactions ^3,4^, necessitating adaptations that ensure the recognition, regulation, and persistence of mutualisms ^5–7^. The resulting partnerships often feature heritable endosymbionts that are intertwined in the evolutionary history of their hosts ^8–10^. As symbiont and host co-diversify, they exhibit parallel lineage splitting over evolutionary timescales ^8,9,11^.

Among animals, obligately beneficial symbionts are typically acquired early in development ^2,4,12–14^. For insects that host endosymbionts within specialized cells (i.e. bacteriocytes) and organs, microbial colonization can take place during embryogenesis or oogenesis ^13–15^. For example, aphids acquire their nutritional endosymbiont, *Buchnera aphidicola*, during embryo development through calibrated cycles of exocytosis and endocytosis ^13^. Endosymbiont cells released from maternal symbiotic organs colonize cells fated to become bacteriocytes in the embryo ^13^. Ants similarly integrate their endosymbiont, *Blochmannia floridanus*, during embryogenesis through a shared regulatory network that is coopted by the microbe to facilitate infection ^14^. Such models, and others ^6,15–18^, continue to offer striking insights into the cellular mechanisms guiding symbiont colonization and proliferation ^12^. But our inability to cultivate most obligate symbionts in pure culture ^18^, compounded by the brief developmental window that their hosts spend aposymbiotically ^6,13,14^, constrains the experimental exchange of symbionts and the ability to test fidelity within co-diversified partnerships.

In contrast to the experimental challenges posed by heritable endosymbionts, partnerships with beneficial microbes acquired from the environment have provided crucial insights into the mechanisms that govern specificity in animal-microbe symbioses ^19–25^. For example, the Hawaiian bobtail squid (*Euprymna scolopes*) can differentiate its bioluminescent symbiont *Vibrio fischeri* from thousands of marine microbial species through an intricately honed process that relies on mechanical, chemical, immunological, and developmental exclusion ^26–29^. Similarly, the bean bug (*Riptortus pedestris*) acquires its crypt-associated gut symbiont *Caballeronia insecticola* every generation from the soil ^19,30–32^. Despite a horizontal acquisition mode ^30^, several factors contribute to a highly selective recognition process, including a microbial-sorting structure within the gut ^20^, the competitive exclusion of non-symbiotic strains ^33^, and, finally, the morphological differentiation and closure of the symbiotic organ following passage by *C. insecticola* ^31^. While horizontally transmitted symbionts lack evolutionary histories that mirror their hosts ^19,22^, the partnerships between the squid-*V.fischeri* and bean bug-*C.insecticola* display striking adaptations for partner recognition at the genetic and molecular levels ^20,21,31,34–36^. Whether these features extend to heritable endosymbionts, however, remains largely unexplored across study systems.

Here, we investigated the specificity of a co-diversified symbiosis by examining the partnership between tortoise leaf beetles (Chrysomelidae: Cassidinae) and their digestive bacterial symbiont, *Candidatus* Stammera capleta (Figures 1, A-H) ^37–39^. The tortoise beetle-*S. capleta* partnership offers a tractable experimental framework to test fidelity in an obligate symbiosis given the microbe’s unique transmission route and colonization dynamics ^37,40,41^.

**Figure 1.**
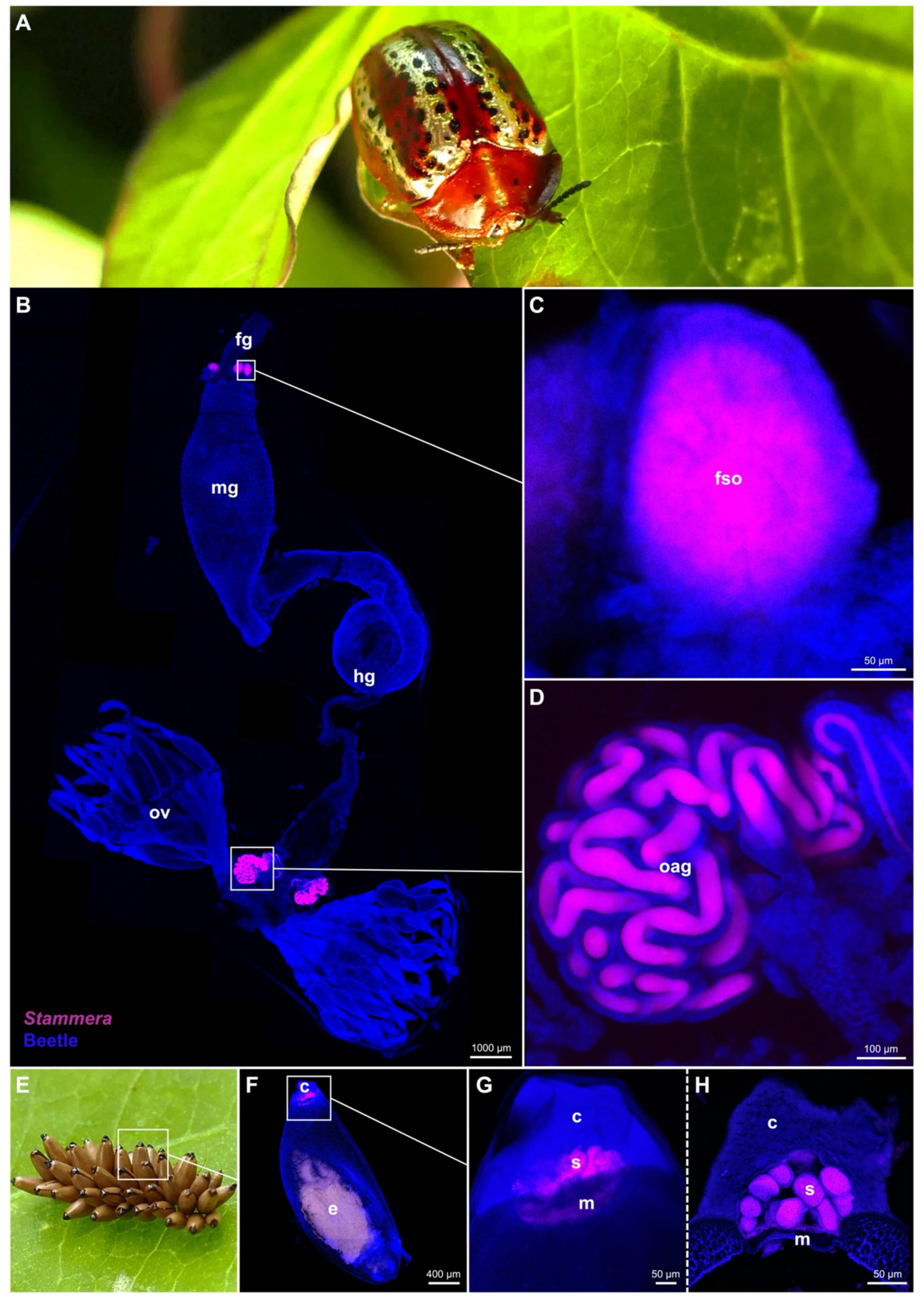
Localization and transmission of *Stammera capleta* in tortoise beetles. (**A**) Adult tortoise beetle (Coleoptera: Cassidinae: *Chelymorpha alternans*). (**B**) Whole-mount fluorescence *in situ* hybridization (FISH) demonstrating *S. capleta* localization within each type of symbiotic organ. (**C**) Foregut symbiotic organs. (**D**) Ovary-associated glands. (**E**) Egg mass bearing caplets deposited at the anterior pole of each egg. (**F-G**) Whole-mount FISH of an egg and its caplet. (**H**) FISH cross-section of a caplet showing enclosed spheres, where *S. capleta* is embedded. Probes used: *S. capleta* (magenta: 16S rRNA) and the beetle host (blue: 18S rRNA). Abbreviations: fg, foregut; fso, foregut symbiotic organs; mg, midgut; hg, hindgut; ov, ovaries; oag, ovary-associated glands; e, embryo; c, caplet; s, *S. capleta*-bearing spheres; m, caplet membrane. Scale bars are included for reference.

Tortoise beetles maintain *S. capleta* in foregut symbiotic organs to facilitate folivory and in ovary-associated glands to ensure transmission (Figures 1, A-D) ^37–39,41–43^. Despite possessing a drastically reduced genome (∼0.2-0.3 Mb), *S. capleta* encodes and exports several plant cell wall-degrading enzymes, including polygalacturonase, a pectinase that underpins convergent nutritional symbioses across leaf beetles ^37–39,44,45^. Tortoise beetles vertically transmit *S. capleta* by depositing an individual caplet at the anterior pole of each egg ^37,40^ (Figure 1E). This transmission mode is reflected in the strict co-cladogenesis between symbiont and host ^38,39,42^. The caplet is populated with symbiont-bearing spheres where the microbe is embedded extracellularly (Figures 1, F-H) ^37,40^. Embryos acquire *S. capleta* late in development by piercing the caplet membrane and consuming the spheres ^40^, which initiates infection ^41^.

Removing the egg caplet disrupts symbiont transmission, yielding *S. capleta*-free (aposymbiotic) larvae that exhibit a diminished digestive phenotype and drastically reduced survivorship ^37,40,41^. However, reapplying the spheres to aposymbiotic eggs experimentally reconstitutes the symbiosis ^40,41^, highlighting a potential experimental mechanism to exchange the microbe across different species of tortoise beetles by leveraging these structures.

In this study, we capitalized on the conserved symbiont transmission route shared by tortoise beetles to demonstrate that: (i) heritable endosymbionts can be exchanged between host species, (ii) non-native symbionts can colonize and differentially restore survivorship in a novel host, with outcomes ranging from full fitness recovery to reduced survival depending on genetic divergence among symbionts, (iii) but that, ultimately, a high level of fidelity governs this partnership during host development and the symbiont’s propagation to the next generation. Our findings highlight the key, complementary roles that partner recognition, transmission fidelity, and local adaptation all play to ensure specificity and stabilize an ancient, co-diversified symbiosis.

## Results and Discussion

### Reciprocal symbiont exchange in tortoise beetles

Symbiont acquisition in tortoise beetles is governed by a strict developmental window ^40,41^. Although the foregut symbiotic organs form three days before larval emergence from the egg, we previously identified the final 24 hours of embryogenesis as the critical period for *S. capleta* colonization ^40,41^. Reapplying symbiont-bearing spheres during this period can restore *S. capleta* infection in aposymbiotic embryos ^40,41^. Given this, we explored whether the same approach could facilitate the exchange of *S. capleta* between different beetle species.

Using the tortoise beetle *Chelymorpha alternans* (Figure 1A) as a model, we validated this method through four experimental treatments: (a) untreated control, (b) eggs with caplets removed (aposymbiotic), (c) eggs with caplets removed but re-supplied with their original symbiont-bearing spheres at the anterior pole (re-infected), and (d) eggs with caplets removed but instead supplied with spheres collected from *Chelymorpha gressoria*, a closely related species (cross-infected) (Figure 2A).

**Figure 2.**
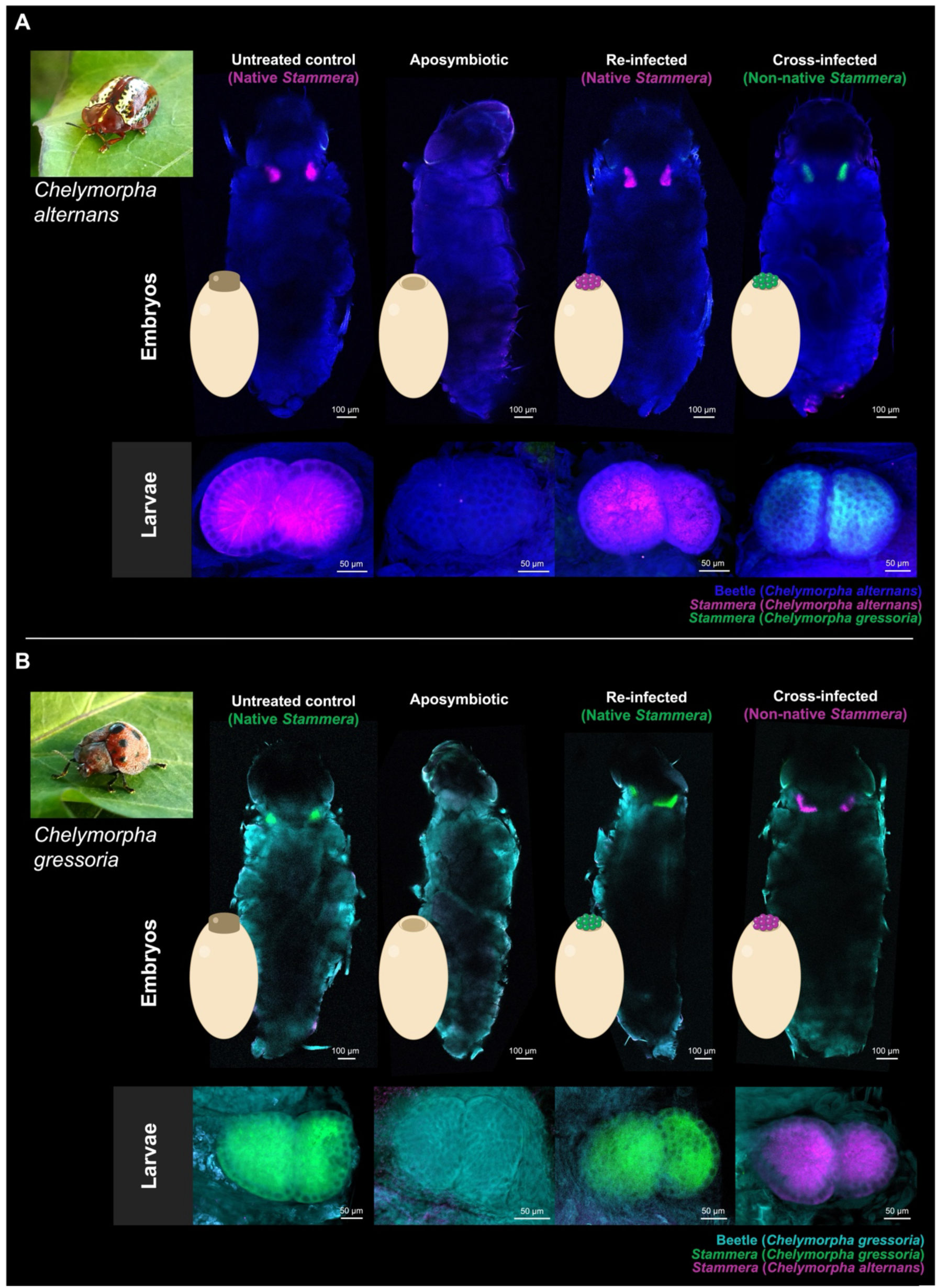
Experimental exchange of *Stammera capleta* in tortoise beetles. Fluorescence *in situ* hybridization (FISH) whole-mount images of entire embryos and foregut symbiotic organs in larvae of (**A**) *Chelymorpha alternans* and (**B**) *Chelymorpha gressoria* across four experimental treatments: untreated control, aposymbiotic (caplets removed), re-infected with the native symbiont, and cross-infected with the non-native symbiont. Probes used: *S. capleta* from *C. alternans* (magenta: 16S rRNA), *S. capleta* from *C. gressoria* (green: 16S rRNA), *C. alternans* host (blue: 18S rRNA), and *C. gressoria* host (cyan: 18S rRNA). Symbiont-specific probes were applied simultaneously during hybridization. Scale bars are included for reference (Figure S1 for replicates).

Consistent with previous studies ^37,40,41^, caplet removal disrupted *S. capleta* transmission, yielding aposymbiotic embryos in contrast to the untreated and re-infected controls (Figure 2A; Figure S1i). Notably, cross-infected embryos (i.e., supplied with spheres from *C. gressoria*) were successfully colonized by non-native *S. capleta* (Figure 2A; Figure S1i). The symbiont occupied the foregut symbiotic organs of its new host, in line with the colonization dynamics observed for the native *S. capleta* of *C. alternans* (Figure 2A; Figure S1i).

To determine whether symbiont exchange is reciprocal across tortoise beetle species, we repeated these assays using *C. gressoria* as a host (Figure 2B). The same four experimental treatments were applied: (a) untreated control, (b) aposymbiotic, (c) re-infected, and (d) cross-infected (i.e., aposymbiotic *C. gressoria* supplied with spheres collected from *C. alternans*). The colonization dynamics mirrored the findings observed in *C. alternans* (Figure 2B; Figure S1ii). Aposymbiotic *C. gressoria* lacked their native *S. capleta*, in contrast to the untreated and re-infected controls. In contrast, cross-infected embryos were successfully colonized by the symbiont of *C. alternans* (Figure 2B; Figure S1ii), indicating that the cross-infection protocol was reciprocal.

To confirm whether these colonization dynamics persist beyond the embryo stage, we evaluated the localization of each symbiont in 5-day-old larvae of both *C. alternans* and *C. gressoria* (Figures 2, A and B). The same patterns were observed across both beetle species: aposymbiotic insects remained uncolonized, while untreated and re-infected controls maintained the native *S. capleta* of each species (Figures 2, A and B; Figures S1, iii and iv). Finally, non-native symbionts continued to occupy the foregut symbiotic organs of their new hosts (Figures 2, A and B; Figures S1, iii and iv). These findings demonstrate that obligate, heritable symbionts can be experimentally exchanged between tortoise beetle species, and that reciprocal cross-infection persists beyond larval eclosion. With this established, we next asked: how do non-native symbionts influence the development of their novel hosts?

### Non-native symbionts spur a gradient of mutualistic outcomes

Numerous insect traits are endowed through symbiosis ^46^, with obligate dependence shaping interactions across diverse clades such as bugs ^47–49^, ants ^14^, termites ^50,51^, bees ^52,53^, and beetles ^5,6,54^. Insects specializing on nutritionally imbalanced diets continue to offer important insights into these obligate interactions and the extent to which host development is stunted in their absence ^55,56^. For example, aposymbiotic aphids deprived of *Buchnera* – and the essential amino acids it supplements – suffer significantly reduced survivorship and complete loss of fecundity ^57,58^. Similarly, bedbugs, unable to balance the nutritional deficiencies of their bloodmeal, are developmentally constrained in the absence of their B vitamin-supplementing endosymbiont ^59^. Such findings are consistent with consequences of aposymbiosis in tortoise beetles, where the experimental loss of *S. capleta* impairs the insects’ ability to process a leafy diet, resulting in low survivorship and failure to reach adulthood ^37,40^. Given these pronounced effects, we asked whether non-native *S. capleta* could rescue aposymbiotic mortality. And if so, how do these beetles develop compared to larvae colonized by the native symbiont?

We pursued the above assays using six species of tortoise beetles: the aforementioned *C. alternans* and *C. gressoria* as before, and in addition, *Chelymorpha bullata*, *Acromis sparsa*, *Aspidimorpha quinquefasciata*, and *Cassida rubiginosa* (Figure 3A; Table S1). Each of these beetles harbors a single, genetically distinct species of *S. capleta* within its foregut symbiotic organs ^37–39^ (Figures 3, B-D; Figures S2-3), which is transmitted extracellularly via egg-associated spheres ^60^.

**Figure 3.**
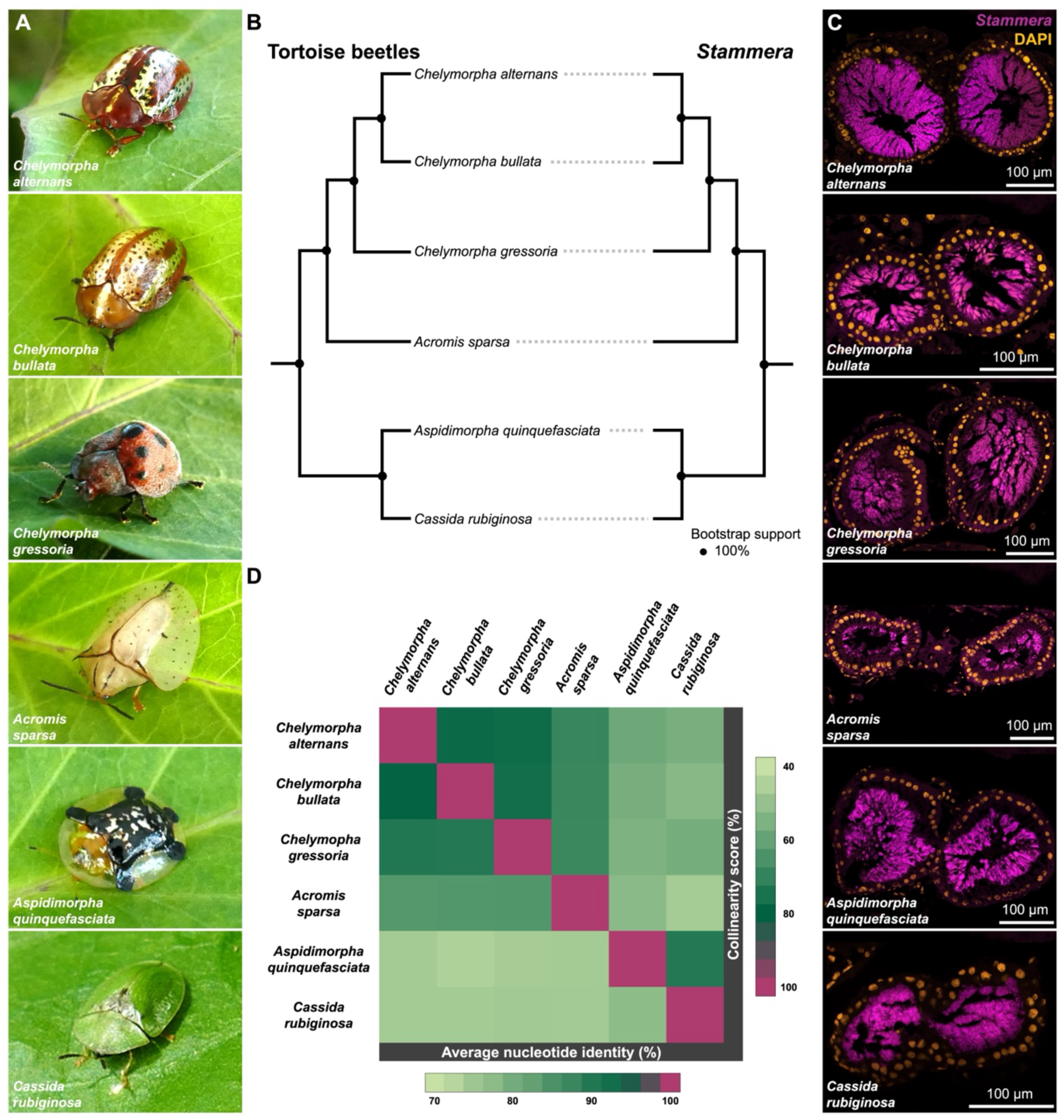
Tortoise beetles co-diversifying with Stammera capleta. (**A**) Tortoise beetles, top to bottom: *Chelymorpha alternans*, *Chelymorpha bullata*, *Chelymorpha gressoria*, *Acromis sparsa*, *Aspidimorpha quinquefasciata*, and *Cassida rubiginosa*. (**B**) Co-cladogenesis between tortoise beetles and their *S. capleta* symbionts based on Maximum Likelihood (ML) phylogenies (Test of Co-cladogenesis, *p* = 0.0099). The host tree leveraged a concatenated alignment of 15 mitochondrial genes, while the *S. capleta* phylogeny is based on a concatenated alignment of 61 single-copy core genes. Node indicates bootstrap support. (**C**) Fluorescence *in situ* hybridization (FISH) cross-sections of foregut symbiotic organs from the beetle species listed above targeting their *S. capleta* symbionts (magenta: 16S rRNA) and DAPI-stained DNA (orange). Scale bars are included for reference. (**D**) Heatmap illustrating genetic distance between different *S. capleta* symbionts. Pairwise comparisons of average nucleotide identity (ANI) (light green: 70%; dark green: 95%) and collinearity scores (light green: 40%; dark green: 85%) are shown.

Using *C. alternans* as a host, we assessed the colonization efficiency of different *S. capleta* species in a novel host and quantified their effects on larval survivorship. We structured our bioassays around eight experimental treatments: (a) untreated control, (b) aposymbiotic eggs, (c) eggs whose caplets were removed but re-infected with the native symbiont, or (d-h) eggs whose caplets were removed but cross-infected with *S. capleta*-bearing spheres collected from each of *C. bullata*, *C. gressoria*, *A. sparsa*, *A. quinquefasciata*, and *C. rubiginosa*.

Five-day-old larvae were dissected to assess their symbiotic status. Experimental manipulation significantly influenced *S. capleta* colonization (Figure 4A; Fisher’s Exact Test, *p* = 0.0005; Table S2A). As demonstrated earlier (Figure 2), removal of the egg caplet disrupted symbiont transmission relative to the untreated and re-infected controls (Figure 4A; pairwise comparisons of proportions, *p* < 0.001; Table S2A). In contrast, cross-infected larvae were consistently symbiotic, irrespective of the donor host (Figure 4A; pairwise comparisons of proportions, *C. bullata* symbiont: *p* = 1, *C. gressoria* symbiont: *p* = 1, *A. sparsa* symbiont: *p* = 1, *A. quinquefasciata* symbiont: *p* = 0.24, *C. rubiginosa* symbiont: *p* = 0.67; Table S2A).

**Figure 4.**
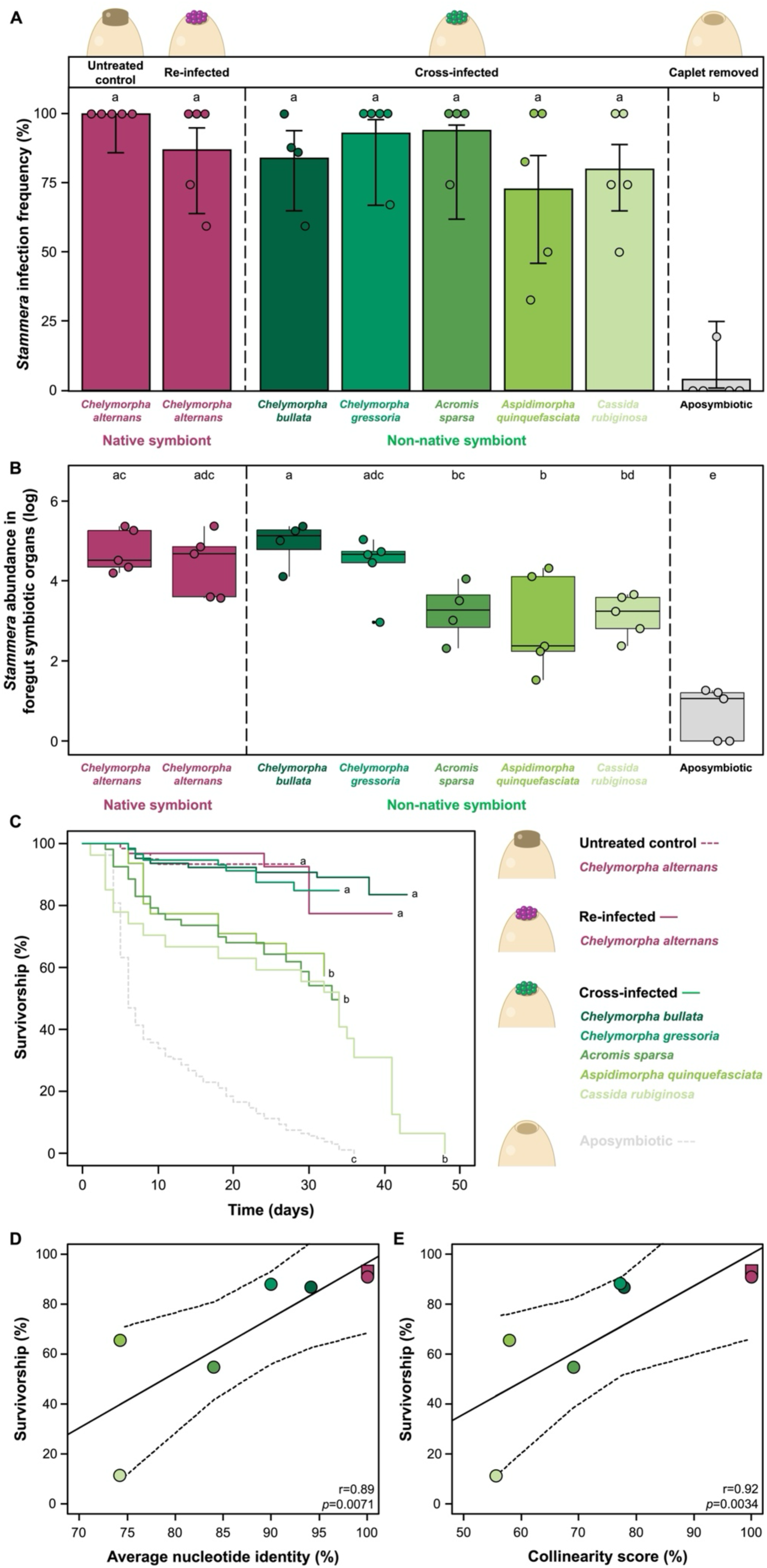
Non-native symbionts differ in their mutualistic potential. (**A**) Symbiont infection frequency in the foregut symbiotic organs of 5-day-old *Chelymorpha alternans* larvae. Bar colors represent experimental treatments: untreated control (5 rep (*n*=24)), aposymbiotic (5 rep (*n*=19)), re-infected with the native bacterium (5 rep (*n*=20)), or cross-infected with the non-native bacterium from *C. bullata* (4 rep (*n*=25)), *C. gressoria* (5 rep (*n*=13)), *A. sparsa* (4 rep (*n*=15)), *A. quinquefasciata* (5 rep (*n*=20)), and *C. rubiginosa* (5 rep (*n*=20)). Whiskers indicate 95% binomial confidence intervals, and dots represent the average of each replicate. Significant differences between treatments are denoted by letters (Fisher’s Exact Test, *p* = 0.0005). (**B**) Symbiont abundance in the foregut symbiotic organs of 5-day-old *C. alternans* larvae, quantified via *16S rRNA* gene copy numbers. Box colors indicate experimental treatments. Lines represent medians, boxes indicate the interquartile range (25–75%), whiskers denote the range, and dots show the average of each replicate. Letters indicate significant differences between treatments (GLM, *p* < 0.001). (**C**) Larval survivorship to adult eclosion across treatments. Line colors correspond to experimental treatments: dotted lines represent untreated control (7 rep (*n*=61)) and aposymbiotic (8 rep (*n*=109)) treatments, while solid lines indicate caplet-free eggs re-infected with the native symbiont (magenta; 7 rep (*n*=32)) or cross-infected with non-native symbionts from 5 different beetle species (green gradient; *C. bullata*, 7 rep (*n*=64); *C. gressoria*, 10 rep (*n*=57); *A. sparsa*, 5 rep (*n*=53); *A. quinquefasciata*, 7 rep (*n*=31); and *C. rubiginosa*, 5 rep (*n*=27)). Beetle survival was monitored until metamorphosis; truncated lines denote surviving individuals censored from the assay. Letters denote significant differences between treatments (Mixed effects Cox regression model, *p* < 0.001). (**D-E**) Correlations between larval survivorship and pairwise genome-wide comparisons of *S. capleta* average nucleotide identity and collinearity scores. Squares represent the untreated control treatment, while circles indicate caplet-free eggs re-infected with the native symbiont (magenta) or cross-infected with non-native symbionts (green gradient). Spearman correlation coefficients (*r*) and their significance (*p*) are shown in each panel.

Given the consistent cross-infection of *S. capleta* in *C. atlernans* stemming from six donor beetle species, we further quantified symbiont loads to determine whether infection success was reflected in the bacterium’s abundance. Quantitative analyses revealed treatment-dependent variation in symbiont titer (Figure 4B; GLM (quasi-Poisson), *χ^2^* = 119.32, d.f. = 7, *p* < 0.001; Table S2B). *S. capleta* abundances were significantly comparable between larvae harboring their native symbiont and those cross-infected with *C. bullata*, *C. gressoria*, and *A. sparsa* symbionts (Figure 4B; pairwise comparison, reinfection: *p* =1, *C. bullata* symbiont: *p* = 1, *C. gressoria* symbiont: *p* = 1, *A. sparsa* symbiont: *p* = 0.084, Table S2B). Conversely, larvae cross-infected with *A. quinquefasciata* and *C. rubiginosa* symbionts contained lower symbiont titers than the untreated larvae (Figure 4B; pairwise comparison, *A. quinquefasciata*: *p* < 0.001, *C. rubiginosa*: *p* < 0.001, Table S2B). QPCR amplification in aposymbiotic individuals was indistinguishable from background (NTC) levels, indicating that the reported copy numbers are likely overestimations and that aposymbiotic beetles are devoid of symbionts, as corroborated by the diagnostic PCR results (Figure 4A) and consistent with prior studies ^37,40,41^.

We next quantified how these non-native symbionts shaped the development of *C. alternans* larvae and found that their effects on host survival varied, yielding a gradient of mutualistic outcomes in larvae (Figure 4C; Cox’s model, *χ^2^* = 229.044, d.f. = 7, *p* < 0.001; Table S2C). While all cross-infected treatments consistently outpaced aposymbiotic insects (Log-rank tests, *p* < 0.001, Table S2C), only two non-native *S. capleta* species completely rescued larval survivorship to levels mirroring to the untreated and re-infected controls (Figure 4C). These symbionts stem from *C. bullata* (Log-rank tests, *p* = 0.7 and 0.99; Table S2C) and *C. gressoria* (Log-rank tests, *p* = 0.38 and 0.76; Table S2C), two beetle species that are congeneric (*Chelymorpha*) to the recipient host, *C. alternans* (Figure 3B). In contrast, the three *S. capleta* species associated with *A. sparsa*, *A. quinquefasciata*, and *C. rubiginosa* could only partially rescue the survivorship of their novel host (Log-rank tests, *A. sparsa*, *p* = 0.00029 and 0.0077; *A. quinquefasciata*, *p*= 0.002 and 0.037, and *C. rubiginosa*, *p*= 00013 and 0.00052, respectively; Table S2C), underscoring the diverse developmental consequences of colonization by a non-native symbiont (Figure 4C). Collectively, this indicates that the observed mortality is only partly explained by a reduction in symbiont abundance, as seen in the cross-infection with the *A. quinquefasciata* symbiont (Figure 4B). However, this pattern does not hold for all high-mortality cross-infections, such as those involving the *A. sparsa* symbiont, where symbiont abundance is comparable to those of the native *S. capleta* (Figure 4B). This variation suggests that host survival following our cross-infections may be influenced by factors beyond symbiont abundance, including reduced symbiont activity and/or the degree of compatibility between host and microbe.

We next examined whether the mutualistic potential of *S. capleta* in a novel host correlates with its evolutionary relatedness to the native symbiont. *S. capleta* that are closely related to the original symbiont of *C. alternans* were more likely to rescue larval survivorship than more distantly related species (Figures 4, D and E; Table S2D). Using genome-wide pairwise comparisons of average nucleotide identity (ANI) and collinearity as measures of genetic distance, we observed a strong positive correlation between symbiont relatedness and host survivorship (Figures 4, D and E; Spearman’s rank correlations: ANI, *r* = 0.89, *p* = 0.0071; collinearity score, *r* = 0.92, *p* = 0.0034; Table S2D).

How do different symbiont genotypes influence the survivorship of a novel host? Despite strong metabolic conservation across the *S. capleta* pangenome in categories related to informational processing (e.g. transcription, translation, and replication), different species encode and supplement two iterations of host-beneficial factors to tortoise beetles ^37–39^. All genomes of *S. capleta* species sequenced to date encode polygalacturonase (PG), a pectinase that breaks down homogalacturonan, the most abundant pectic substrate ^37–39^. However, different species also produce rhamnogalacturonan lyase (RL), a secondary pectinase that degrades the heteropolymer rhamnogalacturonan I, or α-glucuronidase (AG), a xylanase involved in hemicellulose digestion.^37–39^.

The six *S. capleta* species used in our study encompass these two configurations of host-beneficial factors: PG and RL, or PG and AG (Table S1). The symbiont of *C. alternans,* along with those of *C. gressoria*, *C. bullata*, and *A. sparsa,* all encode PG and AG ^37–39^. In contrast, the symbionts of *C. rubiginosa* and *A. quinquefasciata* provide their beetle hosts with PG and RL ^37–39^. Since the latter two species only partially restored survivorship in a novel beetle host that typically depends on PG and AG from its native symbiont (Figure 4C; Table S2C), we hypothesized that differences in host-beneficial factors might explain the observed variation in mutualistic outcomes. However, this is unlikely to be the sole explanation, since the cross-infected symbiont from *A. sparsa* encodes the same set of plant cell wall-degrading enzymes (PG and AG) as the native *S. capleta* in *C. alternans* yet fails to restore larval survivorship relative to the untreated and re-infected controls (Figure 4C; Table S2C). It is conceivable that differences in the expression or regulation of these host-beneficial factors could contribute to the observed variation. Alternatively, the mutualistic potential of *S. capleta* in a novel host may be constrained by other features of its genotype rather than the digestive enzymes it supplements. To explore this further, we investigated the role that partner recognition plays in shaping the symbiotic outcome between tortoise beetles and *S. capleta*. Specifically, we asked: are genetically distant species of *S. capleta* less likely to be recognized as symbionts by their host?

### Recognition of an obligate symbiont

Animals that acquire their symbionts horizontally must winnow thousands of encountered microbes down to their selected partner ^29,30^. These have shed light on the intricate molecular mechanisms behind symbiont acquisition and partner recognition ^22,26,32^. In addition to enriching the environment with beneficial microbes ^30,61,62^, the host deploys a honed immune system to facilitate symbiont colonization by removing environmental, and potentially pathogenic microbes ^28,29,31,35,63,64^. In bean bugs, hundreds of antimicrobial peptides are expressed in the gut, including crypt-specific cysteine-rich peptides ^35,65,66^. These peptides damage the membranes of diverse bacteria but not the crypt-dwelling symbiont, *C. insecticola*, which is resistant ^35,65,66^. Cnidarians, while indiscriminate in their uptake of microalgae from the environment, eject non-symbiotic dinoflagellates through vomocytosis (i.e., engulfment and expulsion by phagocytes) ^24^. Similarly, the Hawaiian bobtail squid discriminates for bioluminescent *V. fischeri* through hemocyte binding of its outer membrane ^28,63^. Non-symbionts induce an elevated immune response defined by the enrichment of antimicrobial peptides in the symbiotic light organ ^29^, complementing mechanical and biochemical selective measures ^21,34^.

Given the varied symbiotic outcomes of *S. capleta* colonisation along a genetic gradient (Figures 4, C and D), we investigated whether non-native symbionts trigger a distinct host response. To address this question, we sequenced the transcriptome of the foregut symbiotic organs of *C. alternans* larvae (5-days-old) under three experimental conditions: (a) re-infection with the native symbiont, (b) cross-infection with the symbiont of *C. gressoria* and (c) cross-infection with the symbiont of *A. quinquefasciata* (Figure 5A). The two non-native symbionts were chosen because of their contrasting effects on host development — while the *C. gressoria* symbiont fully restores larval development in its novel host, the *A. quinquefasciata* symbiont provides only partial rescue (Figure 4B).

**Figure 5.**
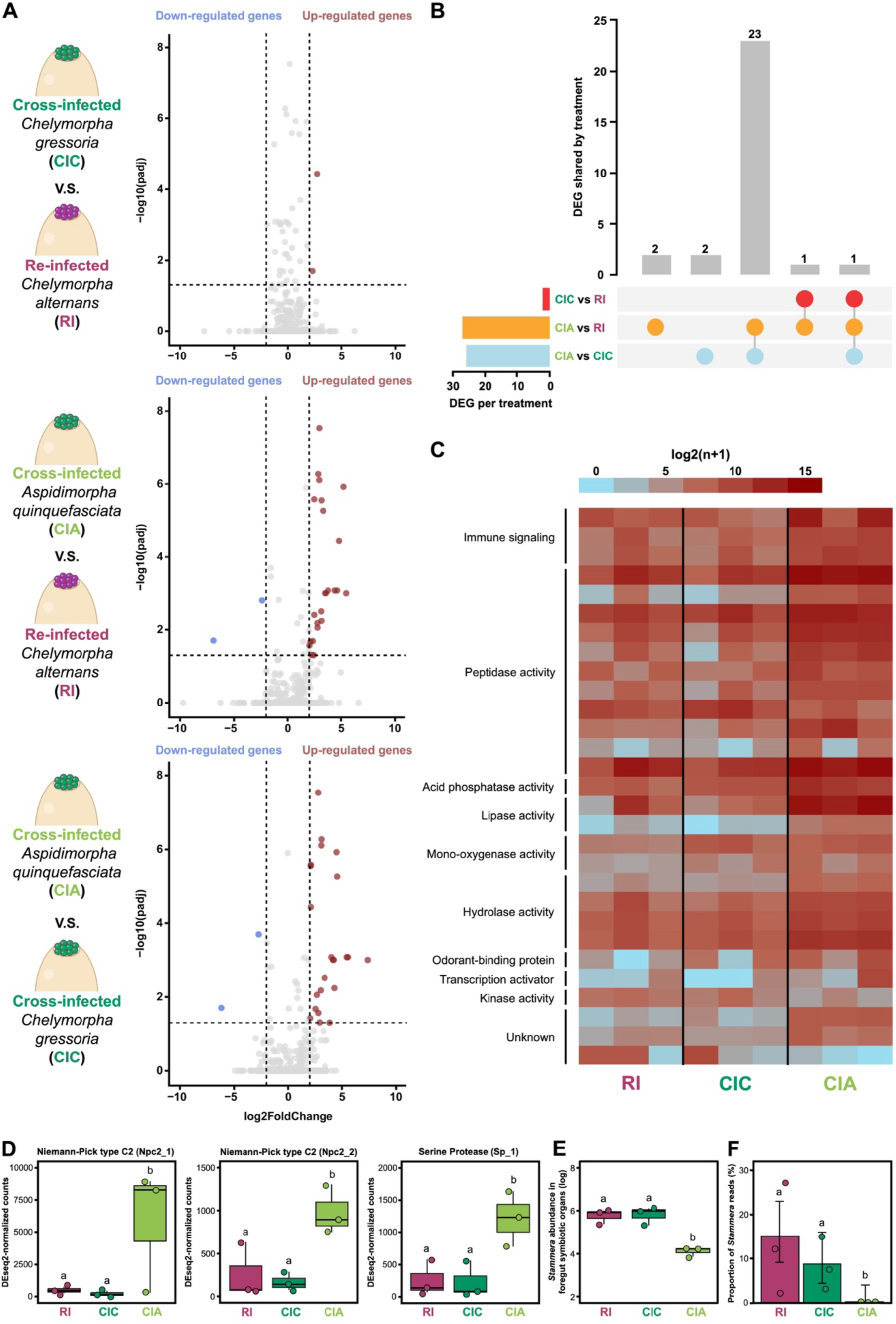
Host response to colonization by a non-native symbiont. (**A**) Differential gene expression in the foregut symbiotic organs of *Chelymorpha alternans* following colonization by native and non-native symbionts. Comparisons include infections by: *Chelymorpha gressoria* symbiont vs. native symbiont (top), *Aspidimorpha quinquefasciata* symbiont vs. native symbiont (middle), and *C. gressoria* symbiont vs. *A. quinquefasciata* symbiont (bottom). Genes with significant expression changes (adjusted *p* < 0.05 and fold change > 2) are highlighted: down-regulated genes in blue and up-regulated genes in red. Three replicates were analyzed per treatment. (**B**) UpSet plot showing differentially expressed genes (DEGs) across treatments: non-native *C. gressoria* symbiont vs. native symbiont (red), non-native *A. quinquefasciata* symbiont vs. native symbiont (orange), and *C. gressoria* symbiont vs. *A. quinquefasciata* symbiont (blue). Horizontal bars represent the total number of DEGs per treatment, while vertical bars indicate shared and unique DEGs across treatments. (**C**) Heatmap of log2(x+1) normalized counts, illustrating host DEG expression patterns in response to colonization by *S. capleta* symbionts. (**D**) Expression levels of host immune genes after colonization of foregut symbiotic organs by *S. capleta* symbionts. Gene counts were normalized using DESeq2’s median of ratios. Bar colors represent experimental treatments: re-infected with the native bacterium (magenta) and cross-infected with the non-native bacterium from *C. gressoria* (dark green) or *A. quinquefasciata* (light green). Medians are shown as lines, boxes represent the 25th to 75th percentiles, whiskers denote the range, and dots indicate the replicates. Letters show significant differences between treatments (Npc2_1 (NB-GLM, *p* < 0.001), Npc2_2 (NB-GLM, *p* < 0.001), Sp_1 (LM, *p* = 0.0012)). (**E**) Symbiont abundance in the foregut symbiotic organs of 5-day-old *C. alternans* larvae, quantified via *16S rRNA* gene copy numbers. Box colors indicate experimental treatments. Lines represent medians, boxes indicate the interquartile range (25–75%), whiskers denote the range, and dots show the replicates. Letters indicate significant differences between treatments (LM, *p* < 0.001). (**F**) Proportion of *S. capleta* reads mapped to the symbiont genome relative to total reads mapped (symbiont + beetle genomes). Whiskers denote 95% binomial confidence intervals, and dots represent the replicates. Letters indicate significant differences between treatments (Fisher’s Exact Test, *p* < 0.001).

Paired-end, dual RNA sequencing (∼30 million reads) was performed on an Illumina NextSeq 2000 and spanning three biological replicates for each treatment (Table S3A). Strikingly, only two host genes were differentially expressed between larvae bearing the *C. gressoria* symbiont and insects colonised by their native *S. capleta* (Figure 5A; Tables S3, B and C, adjusted *p* < 0.05). In contrast, 27 genes were differentially expressed in larvae cross-infected with the *A. quinquefasciata* symbiont compared to the re-infected control (Figure 5A; Tables S3, B and D, adjusted *p* < 0.05). The *A. quinquefasciata* symbiont, which only partially rescues larval development, triggers a more pronounced host response compared to a non-native symbiont that fully restores it (Figure 5A).

When the two cross-infected treatments were compared (*A. quinquefasciata* vs. *C. gressoria*), we found 26 genes to be differentially expressed (Figure 5A; Tables S3, B and E; adjusted *p* < 0.05). Notably, 23 of these genes were also differentially expressed when larvae colonized by their native *S. capleta* were compared to the cross-infected treatment featuring the *A. quinquefasciata* symbiont (Figure 5B; Table S3F, PERMANOVA, *p* = 0.025; Figure S4 and Table S4A). This substantial overlap indicates that the *C. gressoria* symbiont appears to recapitulate the metabolism of its novel host as would the native *S. capleta*.

Genes encoding Niemann-Pick type C2 (NPC2) proteins were highly expressed in cross-infected larvae colonized by the *A. quinquefasciata* symbiont compared to re-infected control and cross-infected larvae harboring the *C. gressoria* symbiont (Figures 5, C and D; Tables S3, B-E; adjusted *p* < 0.05; *Npc2_1*, NB-GLM, *χ*² = 38.7, d.f. = 2, *p* < 0.001; *Npc2_2*, NB-GLM, *χ*² = 35.3, d.f. = 2, *p* < 0.001; Table S4B). NPC2 proteins are known to be involved in sterol homeostasis, steroid biosynthesis, and in innate immunity by modulating the IMD pathway in insects ^67–70^. As demonstrated in *Drosophila melanogaster* and the Asian honeybee (*Apis cerana*), NPC2-encoding genes become highly expressed following pathogenic challenges ^68,70^. These proteins actively combat microbial infections by directly binding to bacterial and fungal cell walls ^68,70^, thereby stimulating the expression of antimicrobial peptides ^67–69^. It is conceivable that *S. capleta* from *A. quinquefasciata* induces *Npc2* expression because of structural differences in its cell wall compared to the symbionts of *C. gressoria* and *C. alternans*. Supporting this, while the *S. capleta* species from *C. gressoria* and *C. alternans* retained a subset of ligases involved in peptidoglycan biosynthesis (including *murD*, *murE*, D-alanine:D-alanine ligase) and outer membrane assembly proteins (including *BamA* and *PhoE*), the encoding genes are absent from the *A. quinquefasciata* symbiont genome ^38,39^.

Colonization by the *A. quinquefasciata* symbiont also led to elevated expression of protease-encoding genes, including numerous cysteine cathepsins and a serine protease (Figures 5, C and D; Figure S5; Tables S3, B-E; adjusted p < 0.05; NB-GLM/GLM-quasipoisson, *p* < 0.001; Table S4B). These enzymes are known to have pleiotropic roles in digestion, development, and immune function. In response to bacterial pathogens, serine proteases are shown to activate the Spätzle-mediated Toll-like receptor and the prophenoloxidase pathway in *D. melanogaster* and the mealworm beetle *Tenebrio molitor* ^67–69^. Cathepsins, on the other hand, are lysosomal proteases and are classified into over 10 subfamilies based on their primary sequence and enzymatic activity ^71^. Many members of the cathepsin superfamily are biologically active and are deployed by insects to combat pathogenic infections and regulate microbial populations along epithelial cells in the gut ^64,72,73^.

Our findings suggest that host immunity may contribute to symbiont recognition. Yet they also raise the possibility that a fine-tuned metabolic interplay between host and symbiont, mediated by transcriptional crosstalk, could be essential for proper symbiont growth. Our results further reveal upregulation of genes involved in metabolic functions, such as lipase activity (Figure 5C, Figure S5, Tables S3, B-E; adjusted *p* < 0.05; Table S4B). Such changes could represent a compensatory host response to reduced nutrient access arising from colonization by a metabolically suboptimal symbiont. Consistent with this interpretation, the native (*C. alternans*) and *A. quinquefasciata* symbionts differ in their potential metabolic profiles, especially in lipid metabolism ^37^. *C. alternans* and *C. gressoria* symbionts encode four and three genes, respectively, related to lipid transport and metabolism, whereas the *A. quinquefasciata* symbiont lacks genes associated with these functions.

We next quantified symbiont abundance using RNA-seq on cDNA and detected significant differences among treatments (Figure 5E; LM, F _(2,9)_ = 42.12, *p* < 0.001; Table S4C), consistent with the previously observed proliferation levels of these non-native symbionts (Figure 4B). *S. capleta* titers did not differ significantly between larvae colonized by their native symbiont or cross-infected with the *C. gressoria* symbiont (Figure 5E; pairwise comparison, *p* = 1; Table S4C). In contrast, both groups contained higher symbiont abundance than larvae cross-infected with *S. capleta* from *A. quinquefasciata* (Figure 5E; pairwise comparison, *p* < 0.001 in both cases; Table S4C). We also mapped *S. capleta* reads to measure the relative presence of symbiotic RNA. The proportion of symbiont transcripts differed significantly across the three experimental groups (Figure 5F; Fisher’s Exact Test, *p* < 0.001; Table S4D). Larvae colonized by their native symbiont or cross-infected with the *C. gressoria* symbiont exhibited similar proportions of *S. capleta* transcripts (Figure 5F; pairwise comparison of proportions, *p* = 0.23; Table S4D), and both groups showed significantly higher proportions than larvae cross-infected with *S. capleta* from *A. quinquefasciata* (Figure 5F; pairwise comparison of proportions, *p* = 0.00072 and *p* = 0.023; Table S4D). The pronounced reduction in *A. quinquefasciata* symbiont abundance suggests that an immune response may have inhibited its proliferation within the foregut symbiotic organ of a novel host. However, it remains unclear whether this decrease reflects bacterial killing or, alternatively, reduced growth in an unfavorable host environment.

Collectively, our findings indicate that (i) non-native *S. capleta* elicit distinct host responses, reflecting the varied symbiotic outcomes of their colonization in tortoise beetles, where (ii) genetically distant species induce a more pronounced host transcriptional response than those closely related to the native microbe. But given the strict pattern of co-cladogenesis between tortoise beetles and *S. capleta* ^38,39^, even among closely related taxa such as *C. alternans* and *C. gressoria*, we next asked: is specificity reinforced at a finer resolution later in development?

### Fidelity throughout host development and symbiont transmission

We quantified *S. capleta* colonization efficiency in adult beetles to determine whether non-native symbionts can persist throughout host development and propagate to future generations. We assessed the symbiotic status of *C. alternans* hosts that were either re-infected with their native symbiont or cross-infected with *S. capleta* from *C. gressoria* at two time points: 5 and 15 days after adult eclosion. In both cases, and despite the extensive transformation and internal reorganization that takes place during metamorphosis, the symbionts persisted throughout pupation and continued to occupy the foregut symbiotic organs of adult beetles (Figure 6A; GLM (binomial), *χ^2^*= 0, d.f. = 1, *p* = 1; Table S4E).

**Figure 6.**
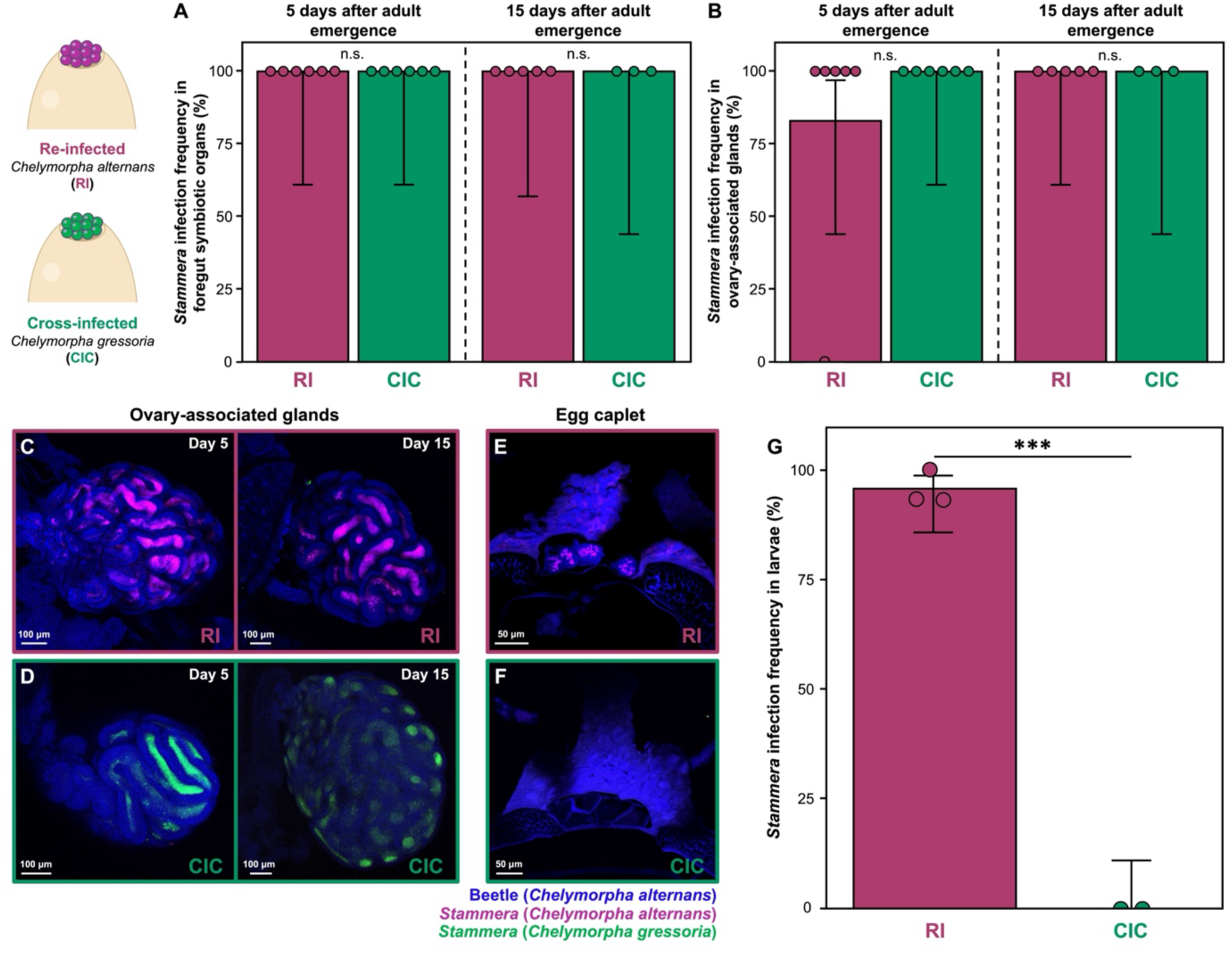
Transmission fidelity ensures propagation of the native symbiont. Infection frequencies of native and non-native symbionts in the (**A**) foregut symbiotic organs and (**B**) ovary-associated glands of *Chelymorpha alternans* females 5- and 15-days following adult emergence. Bar colors represent experimental treatments: females re-infected with the native symbiont (magenta) or cross-infected with the non-native symbiont (green). Whiskers indicate 95% binomial confidence intervals, and dots represent the replicates. Non-significant differences in the interaction between treatment and development time are denoted as “n.s.” (foregut symbiotic organs: GLM, *p* = 1; ovary-associated glands: GLM, *p* = 1). (**C, D**) Fluorescence *in situ* hybridization (FISH) of whole-mounts of ovary-associated glands from *C. alternans* females re-infected with the native symbiont (magenta) or cross-infected with the non-native symbiont (green), visualized at 5- and 15-days following adult emergence. Scale bars represent 100 μm. (**E, F**) FISH of longitudinal egg sections laid by *C. alternans* females either re-infected with the native symbiont (magenta) or cross-infected with the non-native symbiont (green). Dark blue autofluorescence is observed in egg caplets. Scale bars represent 50 μm. Probes used: *S. capleta* from *C. alternans* (magenta: 16S rRNA), *S. capleta* from *C. gressoria* (green: 16S rRNA), and *C. alternans* host (dark blue: 18S rRNA). All symbiont-targeting probes are specific and used simultaneously during hybridization. (**G**) Infection frequency of native and non-native symbionts in larvae emerging from eggs laid by re-infected (*n* = 48) or cross-infected (*n* = 30) females. Whiskers indicate 95% binomial confidence intervals, and dots represent the average of each replicate. Significant differences between symbionts are marked with asterisks (GLM, *p* < 0.001).

Among adult females, we observed that the ovary-associated glands were successfully colonized by non-native *S. capleta*, similar to the re-infected control (Figures 6, B-D; Figure S6; GLM (binomial), *χ^2^* = 0, d.f. = 1, *p* = 1; Table S4E). The mechanism by which symbionts translocate from the foregut symbiotic organs to the ovary-associated glands is still unknown, but it is evident that the non-native symbiont can coopt this route to reach the transmission organs of its novel host (Figures 6, B-D; Figure S6). However, despite successfully colonizing the ovary-associated glands, the vertical transmission of *S. capleta* differed between native and non-native species. While the native symbiont was embedded into the spherical secretions within individual egg caplets, the non-native symbiont was not (Figures 6, E and F; Figure S7).

Eggs laid by cross-infected females contained caplets, but these structures were populated with spheres that were strikingly devoid of *S. capleta* (Figure 6F). Diagnostic PCR confirmed the absence of symbionts in eight egg masses tested, demonstrating that the non-native *S. capleta* failed to embed within the maternally secreted spheres. As a result, larvae emerging from these eggs were entirely aposymbiotic (Figure 6G), in stark contrast to the re-infected treatment (GLM (binomial), *χ^2^* = 88.98, d.f. = 1, *p* < 0.001; Table S4E). The ovary-associated glands cross-infected with the *C. gressoria* symbiont appear to undergo marked morphological remodeling compared to those infected by the native symbiont, from a complex labyrinthine architecture to a simpler conglomerate form between 5 and 15 days (Figures 6D; Figure S6). How, and whether, such a morphological change affects to the failure to transmit the non-native symbiont is unknown.

Extracellular symbiont transmission is widespread in insects ^22^. These routes typically feature a maternal secretions that encapsulate the symbiont ^7,22,30,47,48,74–77^, which are then consumed by offspring to initiate infection ^22^. For instance, plataspid stinkbugs ensure the vertical transmission of their nutritional symbiont, *Ishikawaella capsulata*, by embedding it within a proteinaceous capsule ^7,48,75^. This capsule is primarily composed of an odorant-binding protein that stabilizes the symbiont’s metabolism during extracellular transfer ^7^, compensating for its highly reduced genome ^78^. Similarly, urostylidid stinkbugs propagate their nutritional symbiont, *Tachikawaea gelatinosa*, through jelly-like secretions that are rich in galactan and essential amino acids ^47^. It is conceivable that tortoise beetles rely on similar substrates to produce their *S. capleta*-bearing spheres (Figures 1, F-H). The embedding properties of these substrates may be specifically tailored to the unique cellular and physiological characteristics of each symbiont, which could explain why non-native *S. capleta* fail to populate the spherical secretions of their novel host (Figures 6, E and F). Similar to tortoise beetles, transmission fidelity plays a crucial role for maintaining specificity between beewolves and their defensive ectosymbiont *Streptomyces philanthi* ^79^. While non-symbiotic bacteria can colonize beewolf transmission glands, only the native symbiont can successfully embed within the maternal secretion, thereby ensuring the faithful propagation of this partnership ^79^.

Since many tortoise beetle species are sympatric and often coexist on the same host plant ^60^, we next examined the infection outcome of multiple *S. capleta* strains and explored whether local adaptation to the host environment offers a selective advantage to the native symbiont.

### Competition-based selection reinforces specificity

Competition is a strong selective force shaping microbial community structure in a host environment ^33,80,81^. The host can promote the dominant growth of locally adapted microbes in open systems like the gut, cuticle, or rhizosphere by creating a specialized metabolic landscape ^33,80,81^. Given recent examples of competition-based selection stabilizing specificity in symbioses ^33,82^, we investigated whether this process offers an additional mechanism for tortoise beetles to promote the competitive advantage of their native *S. capleta* during mixed infections.

Using *C. alternans* as a host, we conducted dual-infection assays by introducing two *S. capleta*-bearing spheres to caplet-free eggs: one containing the native symbiont, and the other collected from the congeneric beetle, *C. gressoria* (Figure 7A). Both symbionts successfully colonized the foregut symbiotic organs of five-day-old larvae, demonstrating that tortoise beetles can support simultaneous infections by multiple *S. capleta* strains (Figures 7, Bi-iv; Figure S8). We validated these findings by quantifying symbiont abundance throughout host development (Figure 7C; LM, F _(4,24)_ = 83.61, *p* < 0.001; Table S4F).

**Figure 7.**
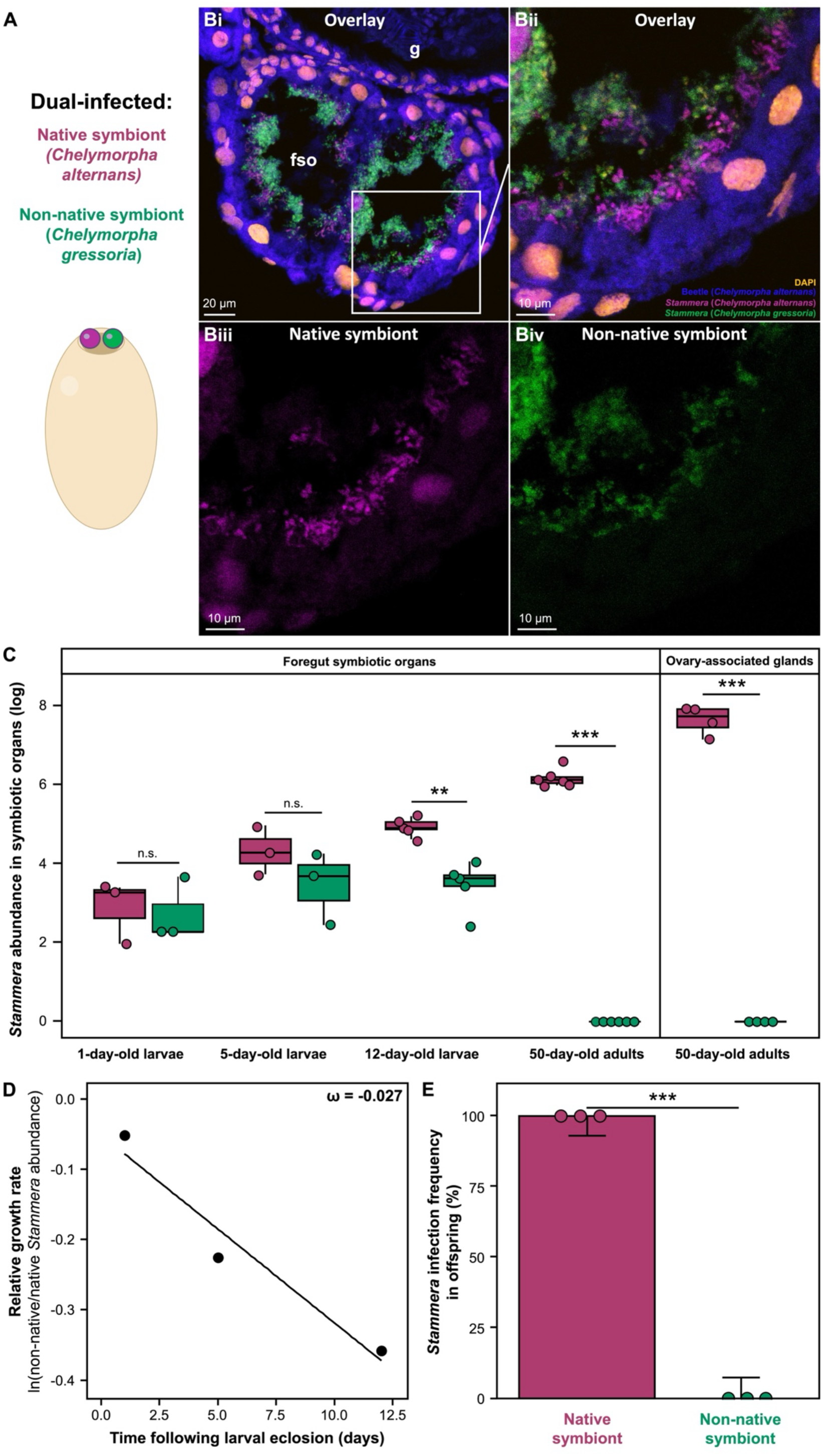
Competition reinforces specificity between tortoise beetles and *Stammera capleta*. (**A**) Illustration of dual-infected *Chelymorpha alternans* embryos. The native symbiont corresponds to *S. capleta* from *C. alternans*, while the non-native symbiont stems from *Chelymorpha gressoria*. (**B**) Fluorescence *in situ* hybridization (FISH) cross-section of foregut symbiotic organs in 5-day-old dual-infected *C. alternans* larvae. (B, ii-iv) correspond to a close-up view of the foregut symbiotic organ. (B, i and ii) correspond to the merged channels images, while (B, iii) corresponds to the native symbiont channel image, and (B, iv) to the non-native channel image. Probes: *S. capleta* from *C. alternans* (magenta: 16S rRNA), *S. capleta* from *C. gressoria* (green: 16S rRNA), *C. alternans* host (dark blue: 18S rRNA), and DAPI-stained DNA (orange). Symbiont-targeting probes are highly specific and used simultaneously during hybridization. Scale bars are included for reference. (**C**) Population dynamics of native and non-native symbionts in the foregut symbiotic organs of dual-infected *C. alternans* beetles during development, quantified via *groEL* gene copy numbers. Box colors indicate either the native symbiont (magenta) or non-native symbiont (green). Lines represent medians, boxes indicate the interquartile range (25–75%), whiskers denote the range, and dots show the replicates. Stars or “n.s.” indicate significant or non-significant differences between symbionts (pairwise comparisons of contrasts; 1-day-old larvae, *p* = 0.99; 5-day-old larvae, *p* = 0.41; 12-day-old larvae, *p* = 0.0024; 50-day-old adults: foregut symbiotic organs, *p* < 0.001; ovary-associated glands, *p* < 0.001). (**D**) Relative growth rate of non-native symbionts compared to native symbionts in the foregut symbiotic organs of dual-infected *C. alternans* larvae. The slope is used to calculate the selection coefficient (ω), shown overlaid on the plot. (**E**) Infection frequencies of native and non-native *S. capleta* in larvae derived from dual-infected *C. alternans* adults. Box colors indicate the experimental treatment: larvae from eggs laid by dual-infected *C. alternans* beetles. Three replicates were collected per treatment (*n* = 48). Whiskers represent 95% binomial confidence intervals, and dots represent the average of each replicate. Significant differences between symbionts are indicated with stars (GLM, *p* < 0.001). Abbreviations: g, gut; fso, foregut symbiotic organs

In newly hatched larvae, native and non-native *S. capleta* initially co-colonized the host at similar titers (Figure 7C; post-eclosion: day 1 (pairwise comparisons of contrasts*, p* = 0.99), day 5 (pairwise comparisons of contrasts*, p* = 0.41); Table S4F). However, symbiont abundance diverged later in larval development (Figure 7C; 12 days post-hatching, pairwise comparisons of contrasts, *p* = 0.0024; Table S4F), despite encoding a near-identical (∼97%) gene set ^39^. Both symbionts share 226 genes but differ in only eight. The *C. alternans* symbiont additionally possesses genes associated with translation and energy metabolism, whereas the *C. gressoria* symbiont encodes genes involved in amino acid and cell wall biosynthesis, reflecting differences in metabolic specialization (Figure S9). The variations in symbiont population growth may be attributed to these specific genetic differences and/or to variations in the regulation of gene expression that led to distinct physiological capacities. To estimate the growth rate of the non-native *S. capleta* relative to the native one during larval development, we calculated a selection coefficient. This yielded a coefficient of −0.027 (Figure 7D), indicating a selective disadvantage for the non-native microbe. A second, independent method leveraging Malthusian parameters corroborated this result, resulting in a similarly negative selection rate of −0.025 for the non-native species.

By adulthood, quantification of symbiont titers revealed the complete loss of the non-native species, while the native *S. capleta* continued to stably colonize both the foregut symbiotic organs and the ovary-associated glands, underscoring the role of microbe-microbe competition in reinforcing specificity between tortoise beetles and *S. capleta* (Figure 7C; pairwise comparisons of contrasts: foregut symbiotic organs, *p* < 0.001; ovary-associated glands, *p* < 0.001; Table S4F). This contrasts the infection dynamics observed when the *C. gressoria* symbiont colonized *C. alternans* alone — where it successfully persisted in both the adult foregut symbiotic organs and ovary-associated glands of its novel host (Figures 6, A-D). Reflecting this, offspring of dual-infected beetles were exclusively colonized by the native symbiont (Figure 7E; GLM (binomial), *χ^2^*= 133.08, d.f. = 1, *p* < 0.001; Table S4G).

Our findings show that once both symbionts colonize the foregut symbiotic organs, the native population rapidly supplants the non-native one, despite their near-functional equivalence, effects on host fitness (Figure 4), and metabolic response (Figure 5). This indicates that the native bacterium is better adapted to its host and reinforces the hypothesis that co-evolutionary history is a key driver of fidelity. We further observed spatial heterogeneity in symbiont distribution across the symbiotic organs. The native bacterium predominantly occupies regions adjacent to the epithelium, whereas the non-native bacterium remains restricted to the lumen (Figures 7, Bi-iv; Figure S8). The observed distribution may partly explain the native symbiont’s competitive advantage, potentially through greater local adaptation and enhanced access to host-derived nutrients. We note that a reciprocal dual-infection experiment in *C. gressoria* would help rule out the alternative possibility that the *C. gressoria* symbiont is intrinsically a weaker competitor due to differences in its metabolic relative to *S. capleta* associated with *C. alternans*.

Overall, our findings align with recent studies on bean bugs ^33^ and mice ^82^, demonstrating that native symbionts consistently outcompete non-native microbes within host environments. This highlights the selective advantage of local adaptation, which can further strengthen specificity within symbiotic partnerships.

## Conclusions and Outlook

We conducted experimental cross-infections in aposymbiotic eggs of the tortoise beetle *C. alternans* using *S. capleta* derived from five additional Cassidinae species. Our results demonstrate that larval survival is restored following these bioassays, but only when the non-native symbionts are closely related to the native one. Consistent with this, the positive correlation between symbiont genetic distance and host survivorship highlights the strong influence of symbiont genotype on host development.

Our findings also demonstrate that while obligate, heritable endosymbionts can successfully establish and fully rescue development in a novel host, long-term persistence and successful vertical transmission are maintained by a robust framework of partner specificity. The introduction of a genetically distant, non-native symbiont triggered a strong transcriptional response in the host, including activation of immune genes. This was accompanied by a decrease in symbiont population, suggesting host discrimination between native and non-native partners. When both were introduced simultaneously, the native symbiont progressively outcompeted the non-native one and was exclusively transmitted to the next generation. Together, these results highlight how the multilayered interplay of host recognition, transmission fidelity, and competitive exclusion acts to ensure that only the native symbiont persists, securing propagation to the next host generation. This selective advantage highlights the evolutionary fine-tuning of the *S. capleta*-tortoise beetle partnership, where co-diversification and local adaptation appear to reinforce a highly specific, heritable symbiosis. Testing whether these mechanisms act in concert to preserve specificity in other vertically transmitted, co-diversified systems would be valuable given the continued development of experimental approaches to clear and reconstitute endosymbionts, as demonstrated in aphids ^83^, weevils ^18^, and plataspid stinkbugs ^84^. Moreover, the application of microinjection into the host hemolymph to reproduce the mounted immune response is a worthy area for future study.

Our results align with and extend findings from horizontally acquired symbioses, such as the *Vibrio fischeri*-Hawaiian bobtail squid and *Caballeronia insecticola*-bean bug systems, where host-driven selection — via immune filtering, niche restriction, and competitive exclusion — ensures the establishment of preferred microbial partners. In the squid, hemocyte-mediated recognition and antimicrobial peptide expression help winnow *Vibrio* from a diverse seawater microbiome ^21,26,28,29,34^, while the bean bug’s gut crypts physically and chemically promote the retention of *Caballeronia* over environmental bacteria ^19,20,30,31,33^. The *S. capleta*-tortoise beetle model demonstrates that even vertically transmitted endosymbionts are subject to similar competitive and immunological pressures, blurring the distinction between horizontal and vertical symbiont selection mechanisms. Future research comparing obligate, facultative, and environmentally acquired symbioses could uncover shared genetic and molecular pathways that mediate partner fidelity, with implications for the establishment and maintenance of complex microbial communities across ecological and evolutionary scales^10,82^.

## Methodology

### Insect rearing

Tortoise beetles were maintained a greenhouse at the Max Planck Institute for Biology in Tübingen, Germany. The insects were reared in mesh containers along with their host plants: *Ipomoea batatas* for *C. alternans*, *C. bullata*, *C. gressoria*, and *A. quinquefasciata* beetles; *Merremia umbellata* for *A. sparsa* beetle; and *Cirsium oleraceum* for *C. rubiginosa* beetle (Table S1). Experimental treatments were maintained in climate chambers at a constant temperature of 26°C, humidity of 60%, and long light regimes (14.30 h/9.30 h light/dark cycles).

### Whole-mount fluorescence *in situ* hybridization

To visualize *S. capleta* in *C. alternans* eggs and adults, whole-mount fluorescence *in situ* hybridization (FISH) was performed, as previously described ^85,86^. Eggs and a single dissection from an ovipositing female, containing the foregut, midgut, hindgut, and ovaries, were fixed in Carnoy’s solution (ethanol: chloroform: acetic acid = 6: 3: 1) at room temperature overnight. After thorough washing with 100% ethanol, the samples were bleached in an alcoholic 6% H_2_O_2_ solution to reduce autofluorescence and incubated at room temperature until they were decolorized. Samples were thoroughly washed with 100% and then 70% ethanol and rehydrated with PBSTx containing 0.3% Triton X-100. They were subsequently incubated with hybridization buffer (20 mM Tris-HCl [pH 8.0], 0.9 M NaCl, 0.01% SDS, 30% formamide) containing 100 nM of each of the probes overnight. The female internal organs were hybridized with the oligonucleotide probes Atto550-SAL227 targeting the 16S rRNA sequence of *S. capleta* from *C. alternans* and Atto488-EUKCA164 targeting the 18S rRNA sequence of *C. alternans* (Table S5A). Eggs were hybridized with the oligonucleotide probes Atto550-SAL227 targeting the 16S rRNA sequence of *S. capleta* from *C. alternans* and Atto488-EUK1195 targeting the 18S rRNA sequence of all eukaryotes (Table S5A). All probes were dually labeled with fluorescent dyes at 5’ and 3’ ends. Samples were washed thoroughly with PBSTx, mounted using ProLong® Gold antifade mounting media (Thermo Fisher Scientific, USA), and observed using an LSM 980 NLO confocal microscope (Zeiss, Germany).

### Fluorescence *in situ* hybridization on paraffin sections

To localize *S. capleta* in foregut symbiotic organs of the six tortoise beetle species and in *C. alternans* eggs, fluorescence *in situ* hybridization (FISH) was applied on paraffin sections. Adult female foregut symbiotic organs and eggs were dissected and fixed in Carnoy’s solution at room temperature overnight. Dehydration was performed by incubating the samples in 70 and 80% ethanol for 10 min each, followed by three 10-min incubations in 96 and 100% ethanol. Samples were gradually transferred into paraffin by passing through three incubations of Roti®-Histol (CarlRoth, Germany) at room temperature (two steps of 40 min and one step overnight), followed by incubations at 60°C in Roti®-Histol: paraffin (1:1 v/v) for 60 min and paraffin (Paraplast plus, Leica, Germany) (three steps of 60 min and one step overnight). The paraffin-embedded samples were cross-sectioned at 10 μm using a microtome and mounted on poly-L-lysine-coated glass slides (Epredia, Germany). Paraffin sections were dried at room temperature overnight and incubated at 60°C for one hour to improve tissue adherence. They were dewaxed with Roti®-Histol in three consecutive steps for 10 min each, followed by ethanol 100% for 10 min. Slides were then dried at 37°C for 30 min. Foregut symbiotic organs were hybridized with the oligonucleotide probe Cya5-SCA600 targeting the 16S rRNA sequence of *S. capleta* from 52 cassidines (Table S5A). Eggs were hybridized with the oligonucleotide probes Atto550-SAL227 targeting the 16S rRNA sequence of *S. capleta* from *C. alternans* and Atto488-EUKCA164 targeting the 18S rRNA sequence of *C. alternans* (Table S5A). Probes, dually labeled with the fluorescent dye at their 5’ and 3’ ends, were dissolved at 900 nM in the hybridization buffer containing 35% formamide (v/v), 900 mM NaCl, 20 mM Tris-HCl pH 7.8, 1% blocking reagent for nucleic acids (v/v) (Roche, Switzerland), 0.02 SDS (v/v), and 10% dextran sulfate (w/v). Hybridization was performed at 46°C for four hours. Sections were then transferred to fresh 48°C washing buffer for 15 min [70 mM NaCl, 20 mM Tris-HCl pH 7.8, 5 mM EDTA pH 8.0, and 0.01% SDS (v/v)] followed by room-temperature washes in PBS (20 min) and Milli-Q® water (1 min). They were counterstained with DAPI (5 μg/ml) for 10 min at room temperature, dipped in Milli-Q® water, ethanol 100%, and dried at 30°C for 10 min. Slides were mounted using the ProLong® Gold antifade mounting media (Thermo Fisher Scientific, USA), cured overnight at room temperature, and visualized using an LSM 980 NLO confocal microscope (Zeiss, Germany).

### Experimental symbiont exchange across beetle species

Egg masses containing 30 or more eggs with well-defined caplets were collected from *C. alternans* mating pairs to avoid pseudo-replication. Egg masses were then separated into different experimental treatments depending on the experiment: untreated control, aposymbiotic, re-infected with the native symbiont, and cross-infected with symbionts from a different beetle species. Aposymbiotic individuals were generated by removing caplets from eggs using sterile dissection scissors and without piercing the developing embryo ^37,40,41^. Re-infection with the native bacterium was performed by eliminating caplets from eggs and reapplying all enclosed spheres to the anterior pole of the egg, as previously described ^40,41^. Cross-infections with non-native symbionts were achieved by removing caplets from eggs and reapplying all enclosed spheres collected from eggs of different beetle species to the anterior pole of *C. alternans* eggs. As with the re-infection procedure, spheres were obtained from eggs of the same mass for each replicate. To test whether *S. capleta* can be reciprocally exchanged across tortoise beetles, whole-mount fluorescence *in situ* hybridization (FISH) was performed on entire embryos and larval foregut symbiotic organs of two beetle species *C. alternans* and *C. gressoria*, as described above. For each beetle species, two to four egg masses were collected from separate females and divided into four experimental treatments: untreated control, eggs with caplets removed (aposymbiotic), eggs with caplets removed but re-supplied with their original symbiont-bearing spheres, and eggs with caplets removed but supplied with spheres collected from different host species. A near-emergence embryo (∼ 10.5 days following oviposition) and 5-day-old larvae were collected after experimental manipulation for each replicate. Samples were placed in 70% ethanol to remove the embryo legs and scoli and dissect the larval foregut symbiotic organs using sterilized scissors. The following oligonucleotide probes were used for *in situ* hybridization: Atto550-SAL227 targeting the 16S rRNA sequence of *S. capleta* from *C. alternans*, Cy5-SCG224 targeting the 16S rRNA sequence of *S. capleta* from *C. gressoria,* Atto488-EUKCA164 targeting the 18S rRNA sequence of *C. alternans*, and Atto488-EUKCG167 targeting the 18S rRNA sequence of *C. gressoria* (Table S5A). Symbiont-targeting probes are specific and used simultaneously during hybridization. The formamide curve generator tool in mathFISH was used to analyze the formamide curves and check the specificity of each probe. All probes were dually labeled with fluorescent dyes at 5’ and 3’ ends. Controls consisted of the two beetle species infected with their native symbionts (Figure S1).

### Phylogenetic inference

Host mitochondrial genomes were extracted from metagenomic assemblies and aligned using MUSCLE (v3.8.1551) ^87^, and as described by Garcia-Lozano and colleagues ^39^ (Table S5B). A concatenated alignment of 15 mitochondrial genes (13 protein-coding genes + two ribosomal rRNA genes) was partitioned to assign the most appropriate substitution model to each gene using PartitionFinder2 (branchlengths=unlinked, models = GTR+G and GTR+I+G, model_selection = bic) ^88^. Phylogenetic analyses were performed using Maximum Likelihood (ML) in RAxML-NG ^89^ (ngen=1000). Members of Spilopyrinae (*Spilopyra sumptuosa*) and Eumolpinae (*Bromius obscurus*, *Colaspidea* sp., and *Paria* sp.) subfamilies from the Chrysomelidae were used as outgroups for this analysis.

To infer the relationships between the 6 corresponding *S. capleta* symbionts, 61 single-copy core genes identified by anvi’o (v8.1-dev) ^90^ and present in all symbionts were extracted from each *S. capleta* genome and aligned using MUSCLE (v3.8.1551) ^39,87^. Concatenated alignments of these genes and four outgroups (*Escherichia coli*, *Buchnera aphidicola*, *Candidatus* Ishikawaella capsulata, and *Vibrio fischeri*) were included to construct a ML phylogeny using RAxML-NG ^89^ (ngen=1000) (Table S5B). The best-fit substitution model was selected using PartitionFinder2 (branchlengths=unlinked, models = GTR+G, GTR+I+G, model_selection = bic) ^88^. The tree reconciliation software eMPRess GUI^91^ was applied to investigate the evolutionary relationship between the 6 beetle species and their symbionts. This software reconciles symbiont and host trees using the DuplicationTransfer-Loss model. ML trees were used as input and the analysis was conducted using the following parameters: duplication cost = 1, transfer cost = 1, and loss cost = 1. The significance of reconciliation between host and symbiont tree was calculated by randomizing the tips of the branches and then, re-calculating the cost to reconcile the phylogenies. Congruent phylogenies are obtained when the original cost of reconciliation is less than expected by chance (*p* < 0.01).

### *S. capleta* genetic distance

A dual approach was used to evaluate sequence similarity and structural gene organization across symbiont genomes^39^. The Average Nucleotide Identity (ANI) between *S. capleta* genomes was calculated using PyANI ^92^, and as implemented in anvi’o ^90^ to assess symbiont relatedness. Gene collinearity across different *S. capleta* lineages was analyzed with MCScanX ^93^ and SynTracker ^94^. Collinearity scores were then calculated using the collinearity scripts from Nowell et al ^95^. To adapt SynTracker for the comparison of different species we used modified parameters (minimal blast identity = 70%, minimal coverage=70%). SynTracker comparison was carried out with the collection of six *S. capleta* genomes as potential references. The Average Pairwise Synteny score (APSS) was calculated using all comparable regions, for each genome pair. Next, the score for each genome-pair was multiplied by the number of comparable regions, normalized by the length of the smaller genome in each genome-pair, and transformed to fit a scale of 1-100.

### Experimental manipulation to quantify the impact of non-native *S. capleta* on beetle development

To estimate whether non-native symbionts can successfully colonize the foregut symbiotic organs of *C. alternans* and restore its survival, egg masses were collected from different female beetles and divided into eight experimental treatments: untreated control, aposymbiotic eggs, eggs whose caplets were removed but re-infected with the native bacterium, or eggs whose caplets were removed but cross-infected with *S. capleta*-bearing spheres collected from 5 different beetle species (*C. bullata*, *C. gressoria*, *A. sparsa*, *A. quinquefasciata*, and *C. rubiginosa*).

Symbiont infection frequencies were validated across all treatments using diagnostic PCR. Five days after hatching, foregut symbiotic organs of larvae were dissected, and DNA was extracted using the E.Z.N.A.® Insect DNA Kit from Omega Bio-tek. PCR primers targeting either specific *S. capleta* symbionts (*groEL* gene) or all symbionts collectively (*16S rRNA* gene) were used to verify the symbiotic status of the host beetles (Table S5C). For cross-infected individuals, a groEL primer pair specific to the native symbiont was used to confirm its successful removal. PCR products were sequenced to confirm primer specificity *in vitro*. Additionally, a specific primer pair for the CO1 gene of *C. alternans* was used as a control for the DNA extraction (Table S5C). Diagnostic PCR was conducted on an Analytik Jena Biometra TAdvanced Thermal Cycler using a final volume of 20 µl containing 1 µl of DNA template, 0.5 µM of each primer, and 2× DreamTaq PCR Master Mix. The following cycle parameters were used: 5 min at 95°C, followed by 34 cycles of 95°C for 30 s, 54.7 to 62.5°C (depending on the primer pair, Table S5C) for 30 s, 72°C for 1 min and a final extension time of 2 min at 72°C. Treatments were carried out in different experimental blocks. Four to five replicates were performed depending on treatment (i.e., n= 13-to-25 larvae).

The impact of the cross-infection procedure on symbiont abundances was assessed by quantitative polymerase chain reaction (qPCR) using an Analytik Jena® qTOWER³ G cycler. DNA samples used previously served as templates. Symbiont abundance was estimated by quantifying the copy number of the *S. capleta 16S rRNA* gene, targeting all Cassidinae species studied, using the following primer pair: TTGACCGCCTGGGGAGTA and TTCGCGTTGCATCGAATTAAAC (Table S5D). Verification of primer specificity was conducted *in silico* by comparison with reference sequences of all symbionts. qPCR reactions were performed in a final volume of 25 µl, containing 1 µl of DNA template, 2.5 µl of each primer (10 µM), 6.5 µl of autoclaved distilled H_2_O, and 12.5 μl of the Platinum SYBR Green qPCR SuperMix-UDG (Thermo Fisher Scientific). Cycling conditions consisted of 3 min activation at 95°C, followed by 45 cycles of 95°C for 15 s, 62°C for 45 s, 72°C for 20 s. A melting curve analysis was conducted from 60 to 95°C, increasing the temperature by one degree every 30 s. Gene copy numbers were estimated from standard curves (10-fold dilution series from 10^−1^ to 10^−8^ ng/µl) generated from purified PCR products, whose DNA concentrations were measured with a NanoDrop^TM^ 1000 spectrophotometer. Since a slight amplification signal was detected in the negative controls (NTCs), the gene copy numbers were corrected by subtracting the average copy number obtained for the NTCs. Treatments were carried out in separate experimental blocks, with four to five replicates per treatment (n= 13-to-25 larvae).

Emerged larvae were observed daily, and survival until adulthood was recorded to assess the impact of non-native symbionts on larval survivorship. Treatments were carried out in different experimental blocks. Five-to-ten replicates were performed depending on treatment (i.e., n=27 to 109 larvae). The relationship between the percentage of survivors after adult emergence and symbiont genetic distance (Average Nucleotide Identity or collinearity score) was also tested.

### RNA sequencing

The transcriptome of foregut symbiotic organs of 5-day-old *C. alternans* larvae was sequenced across three experimental treatments: (a) re-infection with the native symbiont, (b) cross-infection with the symbiont of *C. gressoria* and (c) cross-infection with the symbiont of *A. quinquefasciata*. Three egg masses were collected from separate females and divided into these experimental treatments. Two larvae from each replicate were sampled five days after hatching, and foregut symbiotic organs were dissected using sterile scissors and snap-frozen in liquid nitrogen. RNA extraction was performed for each sample immediately after collection using the QIAGEN RNeasy Mini Kit according to the protocol 4: Enzymatic Lysis and Proteinase K Digestion of Bacteria starting from step 7. This protocol is included in the RNAprotect® Bacteria Reagent Handbook from Qiagen and as referenced in ^39^. Total RNA was quantified using the Qubit^TM^ RNA High Sensitivity (HS) kit. The two samples were pooled to obtain 40 ng of total RNA, which was used as input to prepare nine RNA sequencing libraries (3 biological replicates). Ribosomal RNA was depleted from total RNA using the NEBNext® rRNA Depletion Kit (Human/Mouse/Rat). Libraries were constructed using the NEBNext® Ultra^TM^ Directional II RNA Library Prep kit, and their size was confirmed in a 2100 Bioanalyzer system using the Agilent Technologies High Sensitivity DNA Kit. Sequencing of the final libraries was performed on an Illumina NextSeq 2000 system at the Max Planck Institute for Biology (Tübingen, Germany) using paired-end chemistry (2x100bp) with a depth of ∼ 30 million reads.

Adapter removal and quality filtering of raw reads were performed in Trimmomatic (v0.36) ^96^ and FastQC (v0.12.0) ^97^. Filtered reads were mapped to the *C. alternans* genome ^98^ using HISAT2 (v2.2.0) ^99^. Gene counts were obtained using htseq-count (v0.11.5) ^100^ (stranded=reverse -r pos). After applying a filter, read counts were normalized by the DESeq2’s median of ratios established in the *DESeq2* package ^101^. The likelihood ratio test (LRT), implemented in *DESeq2*, was used to test for differences in host gene expression across treatments. Host genes were considered significantly differentially expressed at adjusted *p*-value (*p* adj) < 0.05 and a fold-change > 2. Volcano plots were generated using the ggplot2 ^102^ and dplyr ^103^ packages for visualizing transcriptome data. We compared the differentially expressed gene (DEG) profile between treatments by testing for significant clusters using a permuted multivariate analysis of variance (PERMANOVA) and applying the function vegan::adonis(). The batch effect was removed using the *LIMMA* package^104^. DEGs were further annotated for immune functions using the *C. alternans* immune gene reference set ^98^, which was derived from the 4IN database (Innate Immunity Genes in Insects: http://bf2i300.insa-lyon.fr:443/home). A heatmap of the log2(x+1) normalized counts was then constructed to illustrate host DEG expression patterns in response to colonization by *S. capleta* symbionts. Analyses were performed in R v. 3.5.3 ^105^.

Symbiont abundances across treatments were quantified by qPCR on an Analytik Jena® qTOWER³ G cycler. cDNA generated from the corresponding RNA samples were used as templates. For cDNA synthesis, 10 ng of total RNA per sample was reverse-transcribed using the RevertAid First Strand cDNA Synthesis Kit (Thermo Fisher Scientific) with oligo(dT) primers. qPCR was performed with the same primer pair (*16S rRNA*) and conditions as described earlier, except that the standard curve was constructed from a 10-fold serial dilution of cDNA (10^−1^ to 10^−8^ ng/μl). Three replicates were performed per treatment (n= 6 larvae). In addition, the proportion of *S. capleta* symbionts within each treatment was estimated as the ratio of RNA-seq reads mapped to the genome of the target symbiont relative the total number of mapped reads (i.e., reads aligned to both the symbiont and host genomes).

### Persistence of non-native *S. capleta* across host development and during transmission

To determine whether non-native *S. capleta* can be maintained during host development, and transmitted to the next generation, egg masses were collected from different *C. alternans* females and divided into two experimental treatments: re-infected with the native symbiont and cross-infected with the symbiont of *C. gressoria*.

Emerged larvae were maintained through pupation and until adulthood. *S. capleta* infection frequency was investigated in the female foregut symbiotic organs and ovary-associated glands five- and fifteen days after adult eclosion, using diagnostic PCR specific to each *S. capleta*-species and as described above. Three to six replicates were performed depending on treatment and sampling time.

Additionally, whole-mount fluorescence *in situ* hybridization (FISH) was performed to visualize each *S. capleta* species in the ovary-associated glands, as described above. For each treatment, one female was collected five and fifteen days after emergence and dissected using sterile scissors. The following oligonucleotide probes were used for *in situ* hybridization: Atto550-SAL227 targeting the 16S rRNA sequence of *S. capleta* from *C. alternans*, Cy5-SCG224 targeting the 16S rRNA sequence of *S. capleta* from *C. gressoria,* and Atto488-EUKCA164 targeting the 18S rRNA sequence of *C. alternans* (Table S5A). Symbiont-targeting probes are specific to each species and were used simultaneously during hybridization.

Three re-infected and two cross-infected females were maintained until sexual maturation. Egg masses laid were recorded daily and subsampled for one month to assess their symbiotic status (7 days post-oviposition). Fluorescence *in situ* hybridization (FISH) was applied to paraffin sections to localize symbionts in the egg caplets, as described above. Eggs were dissected and fixed in Carnoy’s solution at room temperature overnight. The following oligonucleotide probes were used for *in situ* hybridization: Atto550-SAL227 targeting the 16S rRNA sequence of *S. capleta* from *C. alternans*, Cy5-SCG224 targeting the 16S rRNA sequence of *S. capleta* from *C. gressoria,* and Atto488-EUKCA164 targeting the 18S rRNA sequence of *C. alternans* (Table S5A). Controls consisted of untreated eggs from *C. alternans* and *C. gressoria* beetles (Figure S7). Two replicates with one-to-four eggs per replicate were carried out for each FISH treatment.

To confirm the FISH results, four egg masses from cross-infected females were tested by diagnostic PCR targeting the non-native *S. capleta*, pooling two egg caplets per egg mass. Egg masses laid by re-infected and cross-infected females were then maintained until larval emergence. The larvae were observed daily and subsampled (3-day-old larvae) to assess their symbiotic status using *S. capleta*-specific diagnostic PCR, as described above. Two and three replicates were performed for cross-infected (n=30 larvae) and re-infected (n=48 larvae) treatments, respectively.

### Dual-infection assays

Dual-infection assays were performed to investigate whether non-native symbionts can colonize *C. alternans* foregut symbiotic organs and propagate in the presence of the native symbiont. Two symbiont-bearing spheres, one bearing the native symbiont and the other collected from a *C. gressoria* egg caplet, were reintroduced into caplet-free eggs. Both spheres were deposited on the anterior pole of each egg and were observed hourly to ensuring that they were both consumed by the embryo before hatching.

To visualize the symbionts in foregut symbiotic organs of dual-infected *C. alternans* larvae, FISH was applied on paraffin sections, as described above. Foregut symbiotic organs from 5-day-old larvae were dissected and fixed in Carnoy’s solution at room temperature overnight. The following oligonucleotide probes were used for *in situ* hybridization: Atto550-SAL227 targeting the 16S rRNA sequence of *S. capleta* from *C. alternans*, Cy5-SCG224 targeting the 16S rRNA sequence of *S. capleta* from *C. gressoria,* and Atto488-EUKCA164 targeting the 18S rRNA sequence of *C. alternans* (Table S5A). Symbiont-targeting probes are specific and were used simultaneously during hybridization.

Population dynamics of native and non-native symbionts in foregut symbiotic organs of dual-infected *C. alternans* beetles was estimated through qPCR on an Analytik Jena® qTOWER³ G cycler. Dual-infected larvae were sampled one-, five- and twelve days after hatching, and adults (females and males) were sampled 50 days after pupation. Foregut symbiotic organs were dissected, and DNA was extracted using the E.Z.N.A.® Insect DNA Kit from Omega Biotek. To estimate symbiont abundance, the copy number of the *S. capleta groEL* gene from *C. alternans* was quantified with the following primer pair (AGAAGGGATGCAATTTGATAGAGGT and GCAACAGATTCTAATATAGGTAAT, Table S5D), and the copy number of the *S. capleta groEL* gene from *C. gressoria* was quantified with the following primer pair (TGAAGGAATGCAATTTGATAGAGGA and AGCAACAGCTTCTAATATAGGTAAC, Table S5D). Primers were designed to be specific to each symbiont but sharing the same amplicon and efficiency. qPCRs were performed using a final reaction volume of 25 µl containing 1 µl of DNA template, 2.5 µl of each primer (10 µM), 6.5 µl of autoclaved distilled H_2_O, and 12.5 μl of the Platinum SYBR Green qPCR SuperMix-UDG (Thermo Fisher Scientific). The cycling conditions were 3 min of activation at 95°C, followed by 45 cycles at 95°C for 15 s, 60°C for 45 s, 72°C for 20 s, and a melting curve analysis was conducted by increasing the temperature by one degree every 30 s, from 60 to 95°C. The gene copy number was estimated from the standard curves (10-fold dilution series from 10^−1^ to 10^−8^ ng/µl), generated using purified PCR products and measuring their DNA concentration using a NanoDrop^TM^ 1000 spectrophotometer. Treatments were carried out in different experimental blocks. Three to five replicates were performed per treatment.

The relative growth rate of the non-native *S. capleta* compared to the native one in foregut symbiotic organs of *C. alternans* larvae was measured assuming that the mixed infections started at 1:1. The plot of ln(non-native symbiont abundance/ native symbiont abundance) over time was generated with a slope equal to lnω ^106^ :

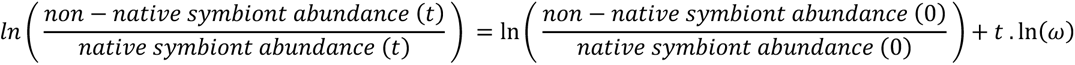

The selection coefficient (ω) for non-native bacterium versus native one was calculated using the e raised to the slope of the line fit to the plot of relative *S. capleta* abundance over time^106^. The linear regression to the non-native symbiont growth curve was fitted using R v. 3.5.3 ^105^ and ggplot2 ^102^. Another method was performed, using the selection rate (r) defined as the difference in Malthusian parameters as follows ^107^ :

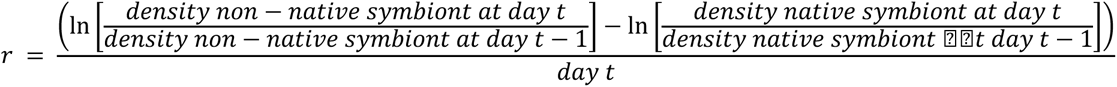

Egg masses laid by dual-infected adults were maintained until larval emergence. Larvae were observed daily and subsampled (3-day-old larvae) to assess their symbiotic status using *S. capleta*-specific diagnostic PCR, as described above. Three replicates were performed per treatment (n=48 larvae).

### Statistical analyses

To assess the colonization efficiency of different *S. capleta* species in *C. alternans* as a novel host, Fisher’s Exact test was used to assess the effect of treatment (untreated, aposymbiotic, re-infected, or cross-infected) on symbiont infection frequency in larval foregut symbiotic organs. Pairwise comparisons between treatments were performed using the pairwise.prop.test() function with Holm correction to adjust for multiple testing. Abundances of different *S. capleta* symbionts in the foregut symbiotic organs of 5-day-old *C. alternans* larvae were analyzed using a generalized linear model with a quasi-Poisson error structure and a log-link function. The treatment, replicate, and experimental block were included in the model as fixed factors. Tukey’s HSD pairwise comparisons were performed using the glht() function with Bonferroni corrections. The survival of *C. alternans* beetles into adulthood was analyzed with a Cox mixed-effect model testing for the effect of experimental treatments (untreated, aposymbiotic, re-infected or cross-infected) and replicates with experimental blocks as a random effect, and using the *coxme* package ^108^. A pairwise comparison between treatments was performed using the pairwise_survdiff() function with Bonferroni corrections in the *survminer* package. The survival data was visualized by computing the Kaplan–Meier survival functions ^109^. Spearman’s rank correlations were used to test the relationship between larval survivorship and symbiont genetic distance (Average Nucleotide Identity or collinearity score) after testing the data distribution.

To compare the expression level of differentially expressed genes after colonization of foregut symbiotic organs by *S. capleta* symbionts, normalized transcripts were analyzed using either a negative binomial generalized linear model implemented by the glm.nb() function, or a general linear model framework depending on the data distribution and after verification that model assumptions were respected. In these statistical models, treatment and replicate were considered as fixed factors. Tukey’s HSD pairwise comparisons were performed using the glht() function with Bonferroni corrections. Abundances of different *S. capleta* symbionts in the foregut symbiotic organs of 5-day-old *C. alternans* larvae (RNA-seq experiment) were analyzed using a general linear model after appropriate data transformation and verification of model assumptions. Treatment, replicate, and their interaction were included as fixed factors, and pairwise comparisons were conducted using Tukey’s HSD with Bonferroni corrections via the glht() function. Fisher’s Exact test was used to assess the effect of treatment (re-infected or cross-infected) on the proportion of *S. capleta* reads mapped to the symbiont genome relative to total reads mapped. Pairwise comparisons between treatments were performed using the pairwise.prop.test() function with Holm correction to adjust for multiple testing. Akaike information criterion was used for model selection.

The effect of experimental treatments (re-infected or cross-infected) on symbiont infection frequencies in foregut symbiotic organs or ovary-associated glands of *C. alternans* females 5-and 15-days post-emergence was analyzed using generalized linear models with a binomial error structure and a logit-link function. In these statistical models, treatment, time, and their interaction were considered as fixed factors. The effect of experimental treatments (re-infected or cross-infected adults) on symbiont infection frequency in their offspring was analyzed using a generalized linear model with a binomial error structure and a logit-link function, and the treatment was considered a fixed factor.

To compare the population dynamics of native and non-native bacteria in the foregut symbiotic organs of dual-infected beetles during development, their abundance was analyzed using a general linear model framework after verification that model assumptions were respected. The time, symbiont, replicate, experimental block, and the interaction between the time and symbiont were used as fixed factors. A pairwise comparison of the interaction was performed using least-squares means with Tukey corrections. Given that the data associated with adult foregut symbiotic organs include both females and males, the effect of sex on *S. capleta* population dynamics was determined by fitting a negative binomial generalized linear model implemented by the glm.nb() function. Beetle sex and symbiont were considered as fixed factors. The effect of beetle development time on the abundance of native and non-native symbionts was analyzed using general linear models after checking that model assumptions were respected and using time as a fixed factor. Tukey’s HSD pairwise comparisons were performed using the glht() function with Bonferroni corrections. The impact of dual-infected beetles on *S. capleta* infection frequencies in their offspring was analyzed using a generalized linear model with a binomial error structure and a logit-link function, and the treatment was considered a fixed factor. Akaike information criterion was used to select all previous models. Statistical analyses were performed in R v. 3.5.3 ^105^ using the prop.test() function to obtain the 95% confidence intervals for a binomial distribution, the *survival* package for survival analyses ^110^, *multcomp* and *lsmeans* packages for pairwise comparisons ^111,112^, *MASS* package for negative binomial generalized linear models ^113^, and *ggplot2* for producing figures ^102^.

## Acknowledgments

Financial support from the Max Planck Society, European Research Council (ERC CoG 101171076 “Symbivore” to H.S.), Alexander Von Humboldt Foundation (to I.P.), German Research Foundation (SA 3105/2-1 to H.S.), and European Molecular Biology Organization (to H.S.) is gratefully acknowledged. This study complies with Nagoya Protocol regulations in the Republic of Panama and France per the following permits: ARG-027-2023 and TREL2302365S/693. We thank Brice Menet for his assistance with beetle maintenance and experiments, and Aileen Berasategui for helpful comments on a previous version of the manuscript. We are also grateful to Sven Helfer, Parthena Prasota, and the Microscopy Facility for their technical support, as well as Donald Windsor for sharing his expertise during field trips to collect beetles.

## Notes

### Competing Interest Statement

The authors have declared no competing interest.

## References

1. Moran, N. A. Symbiosis as an adaptive process and source of phenotypic complexity. Proc. Natl. Acad. Sci. U. S. A. 104 Suppl 1, 8627–8633 (2007).

2. Bennett, G. M. & Moran, N. A. Heritable symbiosis: The advantages and perils of an evolutionary rabbit hole. Proc. Natl. Acad. Sci. U. S. A. 112, 10169–10176 (2015).

3. McCutcheon, J. P. & von Dohlen, C. D. An interdependent metabolic patchwork in the nested symbiosis of mealybugs. Curr. Biol. 21, 1366–1372 (2011).

4. Perreau, J. & Moran, N. A. Genetic innovations in animal-microbe symbioses. Nat. Rev. Genet. (2021) doi:10.1038/s41576-021-00395-z.

5. Simonet, P. et al. Bacteriocyte cell death in the pea aphid/*Buchnera* symbiotic system. Proc. Natl. Acad. Sci. U. S. A. 115, E1819–E1828 (2018).

6. Vigneron, A. et al. Insects recycle endosymbionts when the benefit is over. Curr. Biol. 24, 2267–2273 (2014).

7. Koga, R. et al. Host’s guardian protein counters degenerative symbiont evolution. Proc. Natl. Acad. Sci. U. S. A. 118, e2103957118 (2021).

8. Hosokawa, T. et al. Reductive genome evolution, host-symbiont co-speciation and uterine transmission of endosymbiotic bacteria in bat flies. ISME J. 6, 577–587 (2012).

9. Toju, H., Tanabe, A. S., Notsu, Y., Sota, T. & Fukatsu, T. Diversification of endosymbiosis: replacements, co-speciation and promiscuity of bacteriocyte symbionts in weevils. ISME J 7, 1378–1390 (2013).

10. Suzuki, T. A. et al. Codiversification of gut microbiota with humans. Science 377, 1328–1332 (2022).

11. Moeller, A. H., Sanders, J. G., Sprockett, D. D. & Landers, A. Assessing co-diversification in host-associated microbiomes. J. Evol. Biol. 36, 1659–1668 (2023).

12. McCutcheon, J. P. The genomics and cell biology of host-beneficial intracellular infections. Annu. Rev. Cell Dev. Biol. 37, 115–142 (2021).

13. Koga, R., Meng, X.-Y., Tsuchida, T. & Fukatsu, T. Cellular mechanism for selective vertical transmission of an obligate insect symbiont at the bacteriocyte-embryo interface. Proc. Natl. Acad. Sci. U. S. A. 109, E1230–7 (2012).

14. Rafiqi, A. M., Rajakumar, A. & Abouheif, E. Origin and elaboration of a major evolutionary transition in individuality. Nature 585, 239–244 (2020).

15. Luan, J.-B. et al. Cellular and molecular remodelling of a host cell for vertical transmission of bacterial symbionts. Proc. Biol. Sci. 283, (2016).

16. Luan, J., Sun, X., Fei, Z. & Douglas, A. E. Maternal snheritance of a single somatic animal cell displayed by the bacteriocyte in the whitefly *Bemisia tabaci*. Curr. Biol. 28, 459–465.e3 (2018).

17. Dale, C., Plague, G. R., Wang, B., Ochman, H. & Moran, N. A. Type III secretion systems and the evolution of mutualistic endosymbiosis. Proc. Natl. Acad. Sci. U. S. A. 99, 12397–12402 (2002).

18. Su, Y. et al. Rational engineering of a synthetic insect-bacterial mutualism. Curr. Biol. 32, 3925–3938.e6 (2022).

19. Kikuchi, Y., Hosokawa, T. & Fukatsu, T. An ancient but promiscuous host–symbiont association between *Burkholderia* gut symbionts and their heteropteran hosts. ISME J. 5, 446–460 (2010).

20. Ohbayashi, T. et al. Insect’s intestinal organ for symbiont sorting. Proc. Natl. Acad. Sci. U. S. A. 112, E5179–88 (2015).

21. Kremer, N. et al. Initial symbiont contact orchestrates host-organ-wide transcriptional changes that prime tissue colonization. Cell Host Microbe 14, 183–194 (2013).

22. Salem, H., Florez, L., Gerardo, N. & Kaltenpoth, M. An out-of-body experience: the extracellular dimension for the transmission of mutualistic bacteria in insects. Proc. Biol. Sci. 282, 20142957 (2015).

23. Chen, J. Z., Kwong, Z., Gerardo, N. M. & Vega, N. M. Ecological drift during colonization drives within-host and between-host heterogeneity in an animal-associated symbiont. PLoS Biol. 22, e3002304 (2024).

24. Jacobovitz, M. R. et al. Dinoflagellate symbionts escape vomocytosis by host cell immune suppression. Nat. Microbiol. 6, 769–782 (2021).

25. Hambleton, E. A. et al. Sterol transfer by atypical cholesterol-binding NPC2 proteins in coral-algal symbiosis. Elife 8, (2019).

26. McFall-Ngai, M. J. & Ruby, E. G. Symbiont recognition and subsequent morphogenesis as early events in an animal-bacterial mutualism. Science 254, 1491–1494 (1991).

27. Ruby, E. G. & Asato, L. M. Growth and flagellation of Vibrio fischeri during initiation of the sepiolid squid light organ symbiosis. Arch. Microbiol. 159, 160–167 (1993).

28. Nyholm, S. V., Stabb, E. V., Ruby, E. G. & McFall-Ngai, M. J. Establishment of an animal-bacterial association: recruiting symbiotic vibrios from the environment. Proc. Natl. Acad. Sci. U. S. A. 97, 10231–10235 (2000).

29. Nyholm, S. V. & McFall-Ngai, M. J. The winnowing: establishing the squid-vibrio symbiosis. Nat. Rev. Microbiol. 2, 632–642 (2004).

30. Kikuchi, Y., Hosokawa, T. & Fukatsu, T. Insect-microbe mutualism without vertical transmission: a stinkbug acquires a beneficial gut symbiont from the environment every generation. Appl. Environ. Microbiol. 73, 4308–4316 (2007).

31. Kikuchi, Y., Ohbayashi, T., Jang, S. & Mergaert, P. *Burkholderia insecticola* triggers midgut closure in the bean bug *Riptortus pedestris* to prevent secondary bacterial infections of midgut crypts. ISME J. 14, 1627–1638 (2020).

32. Kikuchi, Y., Hosokawa, T. & Fukatsu, T. Specific developmental window for establishment of an insect-microbe gut symbiosis. Appl. Environ. Microbiol. 77, 4075–4081 (2011).

33. Itoh, H. et al. Host-symbiont specificity determined by microbe-microbe competition in an insect gut. Proc. Natl. Acad. Sci. U. S. A. 116, 22673–22682 (2019).

34. Nyholm, S. V. & McFall-Ngai, M. J. A lasting symbiosis: how the Hawaiian bobtail squid finds and keeps its bioluminescent bacterial partner. Nat. Rev. Microbiol. 20, 315 (2022).

35. Lachat, J. et al. Hundreds of antimicrobial peptides create a selective barrier for insect gut symbionts. Proc. Natl. Acad. Sci. U. S. A. 121, e2401802121 (2024).

36. Lextrait, G. et al. Strict gut symbiont specificity in Coreoidea insects governed by interspecies competition within Caballeronia strains. ISME J. 19, (2025).

37. Salem, H. et al. Drastic genome reduction in an herbivore’s pectinolytic symbiont. Cell 171, 1520–1531.e13 (2017).

38. Salem, H. et al. Symbiont digestive range reflects host plant breadth in herbivorous beetles. Curr. Biol. 30, 2875–2886.e4 (2020).

39. García-Lozano, M. et al. Paleocene origin of a streamlined digestive symbiosis in leaf beetles. Curr. Biol. 34, 1621–1634.e9 (2024).

40. Pons, I. et al. For the road: calibrated maternal investment in light of extracellular symbiont transmission. Proc. Biol. Sci. 289, 20220386 (2022).

41. González Porras, M. Á., et al. Extracellular symbiont colonizes insect during embryo development. ISME Commun. 4, ycae005 (2024).

42. Fukumori, K. et al. Evolutionary dynamics of host organs for microbial symbiosis in tortoise leaf beetles (Coleoptera: Chrysomelidae: Cassidinae). MBio 13, e0369121 (2022).

43. Oguchi, K. et al. Intracellularity, extracellularity, and squeezing in the symbiotic organ underpin nurturing and functioning of bacterial symbiont in leaf beetles. iScience 27, 109731 (2024).

44. García-Lozano, M. & Salem, H. Microbial bases of herbivory in beetles. Trends Microbiol. 33, 151–163 (2025).

45. Berasategui, A. & Salem, H. Microbial determinants of folivory in insects. in Cellular Dialogues in the Holobiont (ed. Thomas C. G. Bosch and Michael G Hadfield) 217–232 (CRC Press, 2020).

46. Douglas, A. E. Multiorganismal insects: diversity and function of resident microorganisms. Annu. Rev. Entomol. 60, 17–34 (2015).

47. Kaiwa, N. et al. Symbiont-supplemented maternal investment underpinning host’s ecological adaptation. Curr. Biol. 24, 2465–2470 (2014).

48. Hosokawa, T., Kikuchi, Y., Nikoh, N., Shimada, M. & Fukatsu, T. Strict host-symbiont cospeciation and reductive genome evolution in insect gut bacteria. PLoS Biol. 4, e337 (2006).

49. Salem, H. et al. Vitamin supplementation by gut symbionts ensures metabolic homeostasis in an insect host. Proc. Biol. Sci. 281, 20141838 (2014).

50. Brune, A. & Dietrich, C. The gut microbiota of termites: digesting the diversity in the light of ecology and evolution. Annu. Rev. Microbiol. 69, 145–166 (2015).

51. Brune, A. Symbiotic digestion of lignocellulose in termite guts. Nat. Rev. Microbiol. 12, 168–180 (2014).

52. Kwong, W. K. et al. Dynamic microbiome evolution in social bees. Science Advances 3, e1600513 (2017).

53. Motta, E. V. S. et al. Host-microbiome metabolism of a plant toxin in bees. Elife 11, e82595 (2022).

54. Salem, H. & Kaltenpoth, M. Beetle–bacterial symbioses: Endless forms most functional. Annu. Rev. Entomol. 67, 201–219 (2022).

55. Douglas, A. E. The microbial dimension in insect nutritional ecology. Funct. Ecol. 23, 38–47 (2009).

56. Hansen, A. K. & Moran, N. A. The impact of microbial symbionts on host plant utilization by herbivorous insects. Mol. Ecol. 23, 1473–1496 (2014).

57. Nakabachi, A. & Ishikawa, H. Provision of riboflavin to the host aphid, *Acyrthosiphon pisum*, by endosymbiotic bacteria, *Buchnera*. J. Insect Physiol. 45, 1–6 (1999).

58. Douglas, A. E. Nutritional interactions in insect-microbial symbioses: aphids and their symbiotic bacteria Buchnera. Annu. Rev. Entomol. 43, 17–37 (1998).

59. Hosokawa, T., Koga, R., Kikuchi, Y., Meng, X.-Y. & Fukatsu, T. Wolbachia as a bacteriocyte-associated nutritional mutualist. Proc. Natl. Acad. Sci. U. S. A. 107, 769–774 (2010).

60. Chaboo, C. S. Biology and phylogeny of the Cassidinae Gyllenhal sensu lato (tortoise and leaf-mining beetles) (Coleoptera, Chrysomelidae). Bull. Am. Mus. Nat. Hist. 305, 1–250 (2007).

61. Lee, K. H. & Ruby, E. G. Effect of the squid host on the abundance and distribution of symbiotic Vibrio fischeri in nature. Appl. Environ. Microbiol. 60, 1565–1571 (1994).

62. Villa, S. M. et al. Specialized acquisition behaviors maintain reliable environmental transmission in an insect-microbial mutualism. Curr. Biol. 33, 2830–2838.e4 (2023).

63. Nyholm, S. V., Stewart, J. J., Ruby, E. G. & McFall-Ngai, M. J. Recognition between symbiotic Vibrio fischeri and the haemocytes of Euprymna scolopes. Environ. Microbiol. 11, 483–493 (2009).

64. Byeon, J. H. et al. A specific cathepsin-L-like protease purified from an insect midgut shows antibacterial activity against gut symbiotic bacteria. Dev. Comp. Immunol. 53, 79–84 (2015).

65. Futahashi, R. et al. Gene expression in gut symbiotic organ of stinkbug affected by extracellular bacterial symbiont. PLoS One 8, e64557 (2013).

66. Jang, S., Ishigami, K., Mergaert, P. & Kikuchi, Y. Ingested soil bacteria breach gut epithelia and prime systemic immunity in an insect. Proc. Natl. Acad. Sci. U. S. A. 121, e2315540121 (2024).

67. Huang, X., Warren, J. T., Buchanan, J., Gilbert, L. I. & Scott, M. P. Drosophila Niemann-Pick type C-2 genes control sterol homeostasis and steroid biosynthesis: a model of human neurodegenerative disease. Development 134, 3733–3742 (2007).

68. Shi, X.-Z., Zhong, X. & Yu, X.-Q. Drosophila melanogaster NPC2 proteins bind bacterial cell wall components and may function in immune signal pathways. Insect Biochem. Mol. Biol. 42, 545–556 (2012).

69. Neophytou, C., Soteriou, E. & Pitsouli, C. The sterol transporter Npc2c controls intestinal stem cell mitosis and host-microbiome interactions in Drosophila. Metabolites 13, (2023).

70. Lee, K. S. et al. Molecular characterization of a Niemann–Pick disease type C2 protein from the honeybee Apis cerana. J. Asia. Pac. Entomol. 17, 555–560 (2014).

71. Turk, V. et al. Cysteine cathepsins: from structure, function and regulation to new frontiers. Biochim. Biophys. Acta 1824, 68–88 (2012).

72. Sun, Y.-X. et al. Functions of Bombyx mori cathepsin L-like in innate immune response and anti-microbial autophagy. Dev. Comp. Immunol. 116, 103927 (2021).

73. Conus, S. & Simon, H. Cathepsins and their involvement in immune responses. Swiss Med. Wkly (2010) doi:10.4414/smw.2010.13042.

74. Salem, H., Kreutzer, E., Sudakaran, S. & Kaltenpoth, M. Actinobacteria as essential symbionts in firebugs and cotton stainers (Hemiptera, Pyrrhocoridae). Environ. Microbiol. 15, 1956–1968 (2013).

75. Fukatsu, T. & Hosokawa, T. Capsule-transmitted gut symbiotic bacterium of the Japanese common plataspid stinkbug, *Megacopta punctatissima*. Appl. Environ. Microbiol. 68, 389–396 (2002).

76. Salem, H., Onchuru, T. O., Bauer, E. & Kaltenpoth, M. Symbiont transmission entails the risk of parasite infection. Biol. Lett. 11, 20150840 (2015).

77. Hosokawa, T., Kikuchi, Y., Shimada, M. & Fukatsu, T. Symbiont acquisition alters behaviour of stinkbug nymphs. Biol. Lett. 4, 45–48 (2008).

78. Nikoh, N., Hosokawa, T., Oshima, K., Hattori, M. & Fukatsu, T. Reductive evolution of bacterial genome in insect gut environment. Genome Biol. Evol. 3, 702–714 (2011).

79. Kaltenpoth, M. et al. Partner choice and fidelity stabilize coevolution in a Cretaceous-age defensive symbiosis. Proc. Natl. Acad. Sci. U. S. A. 111, 6359–6364 (2014).

80. Sun, Y. et al. Intraspecific competition impacts Vibrio fischeri strain diversity during initial colonization of the squid light organ. Appl. Environ. Microbiol. 82, 3082–3091 (2016).

81. Gu, S. et al. Competition for iron drives phytopathogen control by natural rhizosphere microbiomes. Nat. Microbiol. 5, 1002–1010 (2020).

82. Sprockett, D. D. et al. Home-site advantage for host species-specific gut microbiota. Sci. Adv. 9, eadf5499 (2023).

83. Moran, N. A. & Yun, Y. Experimental replacement of an obligate insect symbiont. Proc. Natl. Acad. Sci. U. S. A. 112, 2093–2096 (2015).

84. Hosokawa, T., Kikuchi, Y., Shimada, M. & Fukatsu, T. Obligate symbiont involved in pest status of host insect. Proc. Biol. Sci. 274, 1979–1984 (2007).

85. Koga, R., Tsuchida, T. & Fukatsu, T. Quenching autofluorescence of insect tissues for *in situ* detection of endosymbionts. Appl. Entomol. Zool. (Jpn.) 44, 281–291 (2009).

86. Pons, I., Renoz, F., Noël, C. & Hance, T. New insights into the nature of symbiotic associations in aphids: Infection process, biological effects, and transmission mode of cultivable *Serratia symbiotica* bacteria. Appl. Environ. Microbiol. 85, (2019).

87. Edgar, R. C. MUSCLE: multiple sequence alignment with high accuracy and high throughput. Nucleic Acids Res. 32, 1792–1797 (2004).

88. Lanfear, R., Frandsen, P. B., Wright, A. M., Senfeld, T. & Calcott, B. PartitionFinder 2: New methods for selecting partitioned models of evolution for molecular and morphological phylogenetic analyses. Mol. Biol. Evol. msw260 (2016).

89. Kozlov, A. M., Darriba, D., Flouri, T., Morel, B. & Stamatakis, A. RAxML-NG: a fast, scalable and user-friendly tool for maximum likelihood phylogenetic inference. Bioinformatics 35, 4453–4455 (2019).

90. Eren, A. M. et al. Community-led, integrated, reproducible multi-omics with anvi’o. Nat. Microbiol. 6, 3–6 (2021).

91. Santi Santichaivekin, Qing Yang, Jingyi Liu, Ross Mawhorter, Justin Jiang, Trenton Wesley, Yi-Chieh Wu, Ran Libeskind-Hadas. eMPRess: a systematic cophylogeny reconciliation tool. Bioinformatics 37, 2481–2482 (2020).

92. Pritchard, L., Glover, R. H., Humphris, S., Elphinstone, J. G. & Toth, I. K. Genomics and taxonomy in diagnostics for food security: soft-rotting enterobacterial plant pathogens. Anal. Methods 8, 12–24 (2016).

93. Wang, Y. et al. MCScanX: a toolkit for detection and evolutionary analysis of gene synteny and collinearity. Nucleic Acids Res. 40, e49 (2012).

94. Enav, H., Paz, I. & Ley, R. E. Strain tracking in complex microbiomes using synteny analysis reveals per-species modes of evolution. Nat. Biotechnol. (2024) doi:10.1038/s41587-024-02276-2.

95. Nowell, R. W. Reubwn/Collinearity: V1.0. (Zenodo, 2017). doi:10.5281/ZENODO.891288.

96. Bolger, A. M., Lohse, M. & Usadel, B. Trimmomatic: a flexible trimmer for Illumina sequence data. Bioinformatics 30, 2114–2120 (2014).

97. Andrews, S. FastQC a quality-control tool for high throughput sequence data. http://www.bioinformatics.babraham.ac.uk/projects/fastqc (2024).

98. Genome assembly of the tortoise leaf beetle *Chelymorpha alternans* (Coleoptera: Chrysomelidae: Cassidinae). figshare https://figshare.com/s/b5485e382ccb43092fc8 (2025).

99. Kim, D., Paggi, J. M., Park, C., Bennett, C. & Salzberg, S. L. Graph-based genome alignment and genotyping with HISAT2 and HISAT-genotype. Nat. Biotechnol. 37, 907–915 (2019).

100. Anders, S., Pyl, P. T. & Huber, W. HTSeq--a Python framework to work with high-throughput sequencing data. Bioinformatics 31, 166–169 (2015).

101. Love, M. I., Huber, W. & Anders, S. Moderated estimation of fold change and dispersion for RNA-seq data with DESeq2. Genome Biol. 15, 550 (2014).

102. Wickham, H., Chang, W. & Wickham, M. H. Package ‘ggplot2’. Create elegant data visualisations using grammar graphics. Version 2, 1–189 (2016).

103. Wickham H, François R, Henry L, Müller K, Vaughan D. *dplyr: A Grammar of Data Manipulation*. R package version 1.*1*.*4* (2023).

104. Ritchie ME, Phipson B, Wu D, Hu Y, Law CW, Shi W, Smyth GK. limma powers differential expression analyses for RNA-sequencing and microarray studies. Nucleic Acids Res. 43, (2015).

105. R Core Team. R: a language and environment for statistical computing. Foundation for Statistical Computing, Vienna, Austria. (2013).

106. Mirchandani, C. et al. Mixed *Wolbachia* infections resolve rapidly during in vitro evolution. PLoS Pathog. 20, e1012149 (2024).

107. Alseth, E. O. et al. The impact of phage and phage resistance on microbial community dynamics. PLoS Biol. 22, e3002346 (2024).

108. Therneau TM, T. M. Package “coxme.” R package version https://cran.hafro.is/web/packages/coxme/coxme.pdf (2015).

109. Mackenzie, G., Kalbfleisch, J. D. & Prentice, R. L. The statistical analysis of failure time data. Statistician 31, 278 (1982).

110. Therneau, T. M. & Grambsch, P. M. The cox model. in *Statistics for Biology and Health* 39–77 (Springer New York, New York, NY, 2000).

111. Hothorn, T., Bretz, F. & Westfall, P. Simultaneous inference in general parametric models. Biom. J. 50, 346–363 (2008).

112. Lenth, R. V. Least-squares means: the R package lsmeans. J. Stat. Softw. 69, (2016).

113. Venables WN, R. B. D. Modern Applied Statistics with S. (Springer, New York, 2002).

