## Supplemental Figures and Tables for "Fidelity in co-diversified symbiosis"

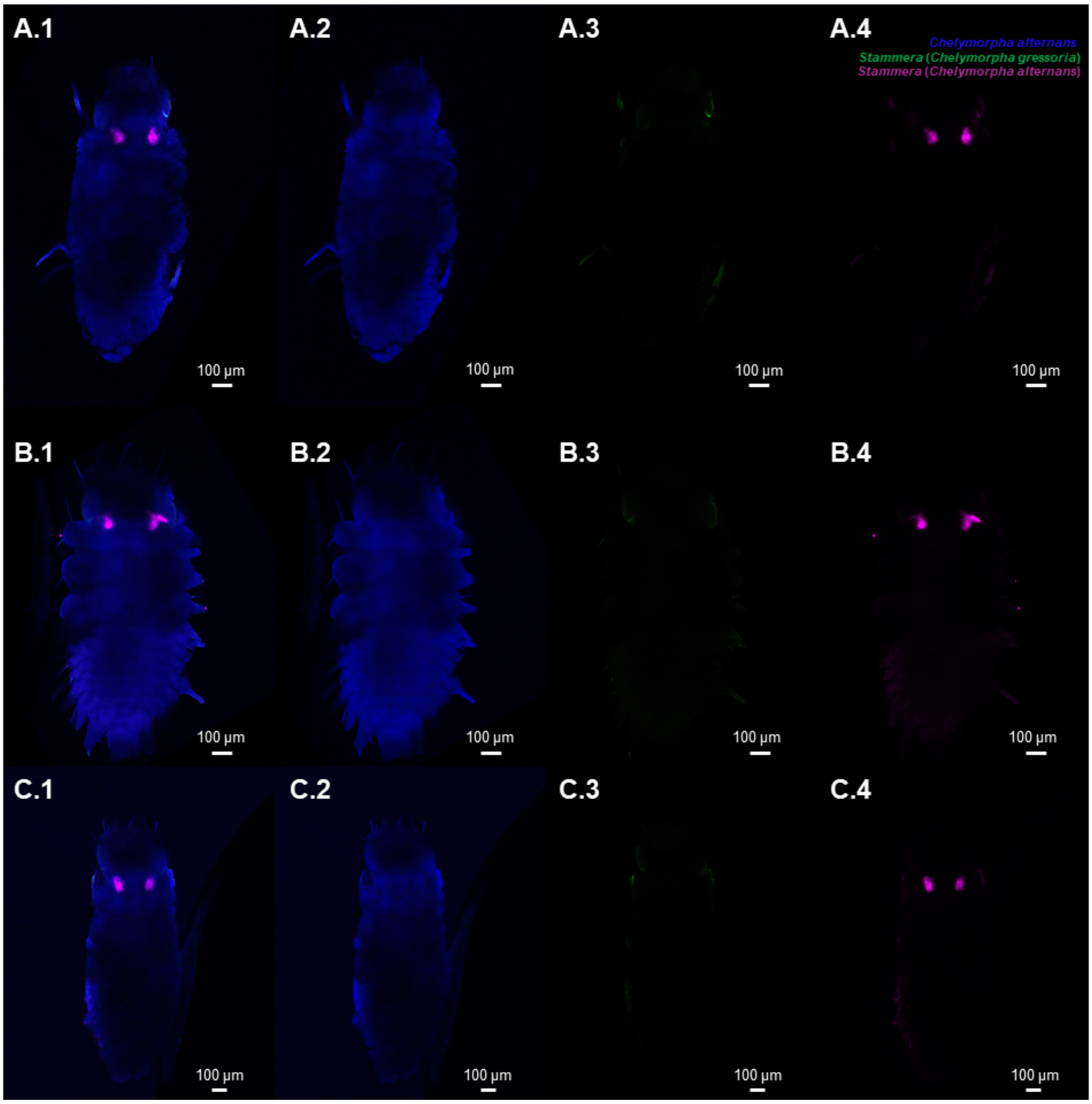


**Figure S1i** (A-C)**.** Fluorescence *in situ* hybridization (FISH) replicates on whole-mounts of *Chelymorpha alternans* embryos (untreated control). Probes used: *Chelymorpha alternans* host (blue: 18S rRNA), *Stammera* from *Chelymorpha gressoria* (green: 16SrRNA), and *Stammera* from *Chelymorpha alternans* (magenta: 16S rRNA). **(1)** correspond to the merged channels images while the others correspond to individual channel images: **(2)** host probe, **(3)** *Stammera* from *Chelymorpha gressoria* probe, and **(4)** *Stammera* from *Chelymorpha alternans* probe. Scale bars (100 µm) are included for reference.

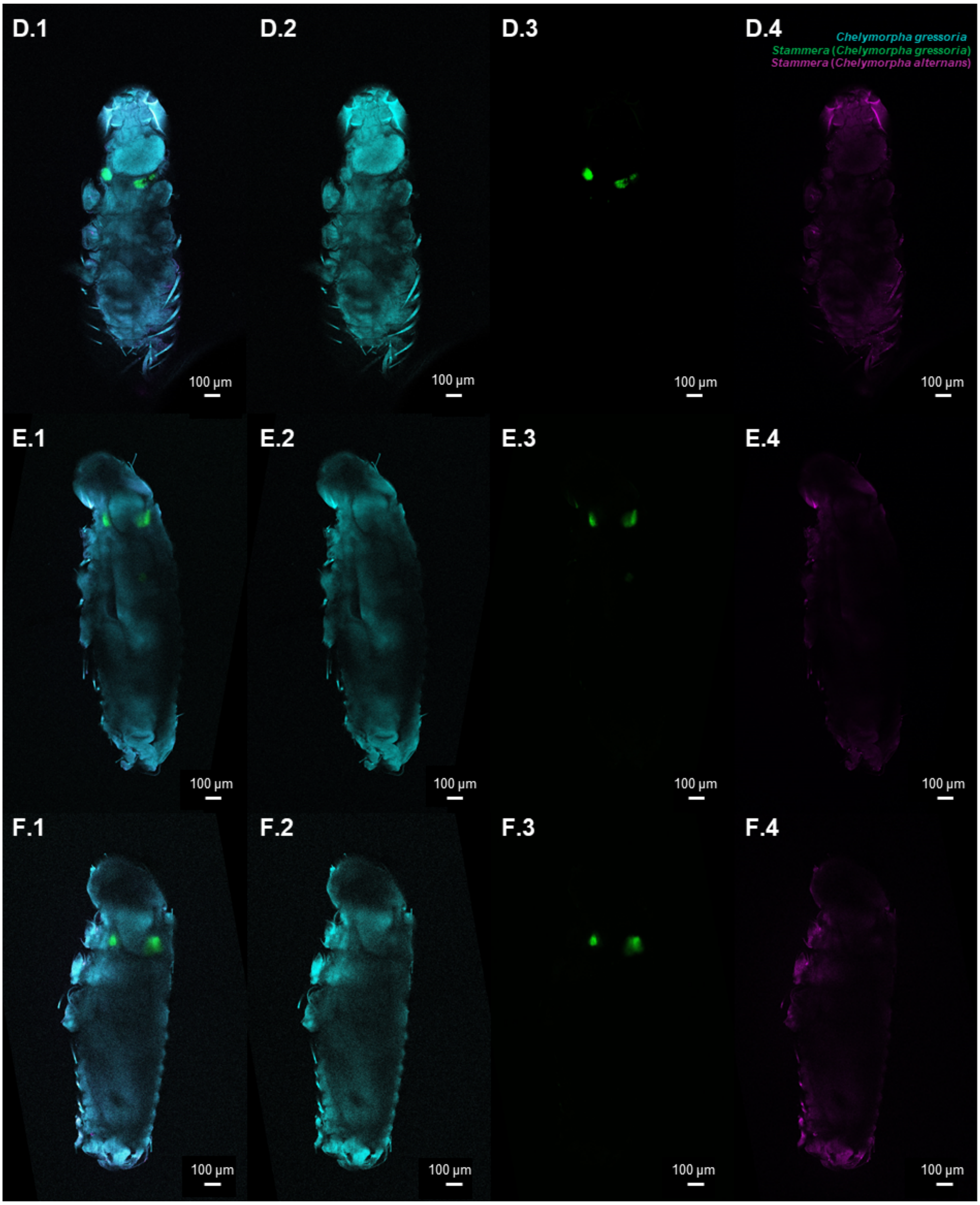


**Figure S1i** (D-F)**.** Fluorescence *in situ* hybridization (FISH) replicates on whole-mounts of *Chelymorpha gressoria* embryos (untreated control). Probes used: *Chelymorpha gressoria* host (cyan: 18S rRNA), *Stammera* from *Chelymorpha gressoria* (green: 16SrRNA), and *Stammera* from *Chelymorpha alternans* (magenta: 16S rRNA). **(1)** correspond to the merged channels images while the others correspond to individual channel images: **(2)** host probe, **(3)** *Stammera* from *Chelymorpha gressoria* probe, and **(4)** *Stammera* from *Chelymorpha alternans* probe. Scale bars (100 µm) are included for reference.

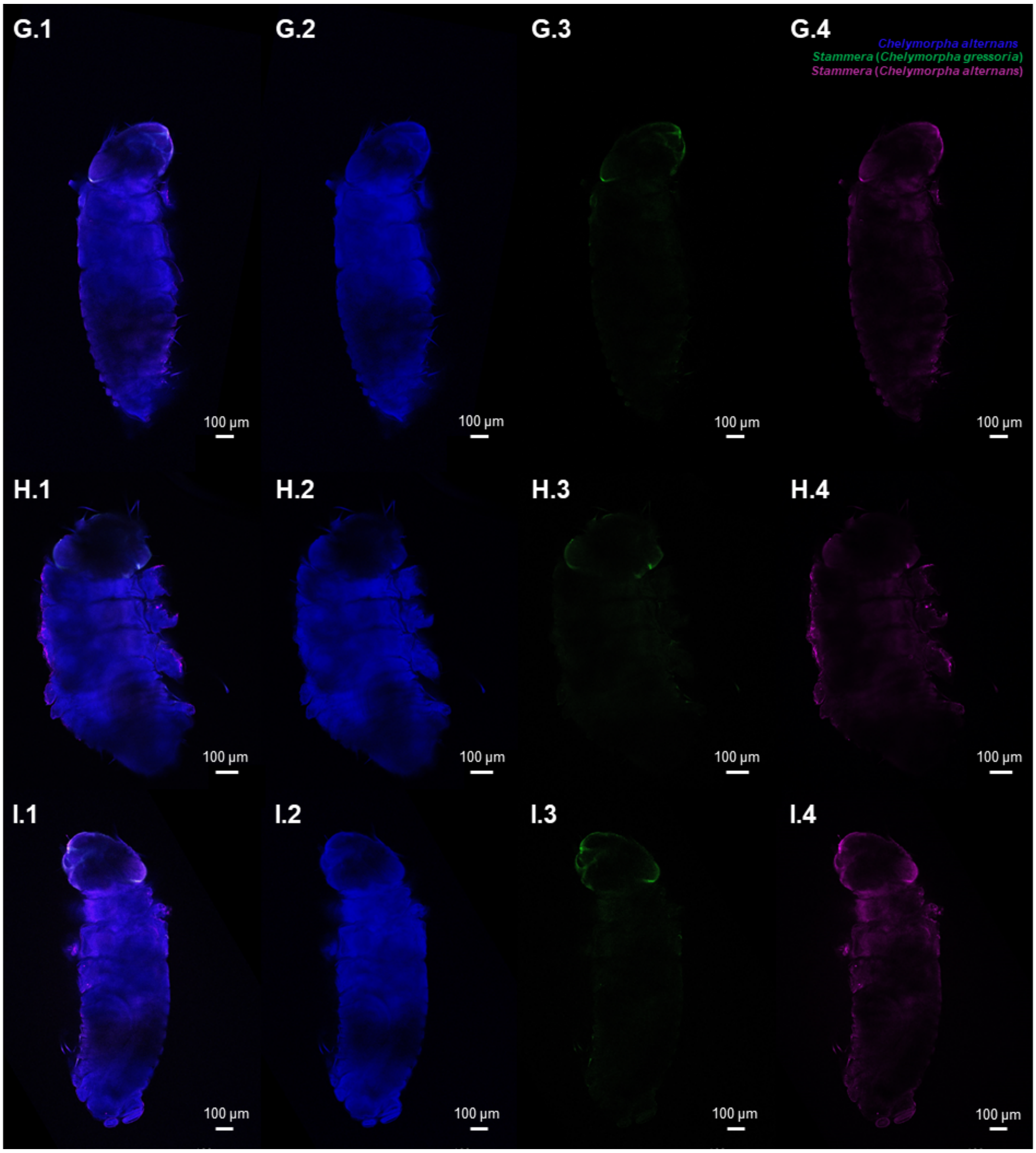


**Figure S1i** (G-I)**.** Fluorescence *in situ* hybridization (FISH) replicates on whole-mounts of *Chelymorpha alternans* embryos (aposymbiotic). Probes used: *Chelymorpha alternans* host (blue: 18S rRNA), *Stammera* from *Chelymorpha gressoria* (green: 16SrRNA), and *Stammera* from *Chelymorpha alternans* (magenta: 16S rRNA). **(1)** correspond to the merged channels images while the others correspond to individual channel images: **(2)** host probe, **(3)** *Stammera* from *Chelymorpha gressoria* probe, and **(4)** *Stammera* from *Chelymorpha alternans* probe. Scale bars (100 µm) are included for reference.

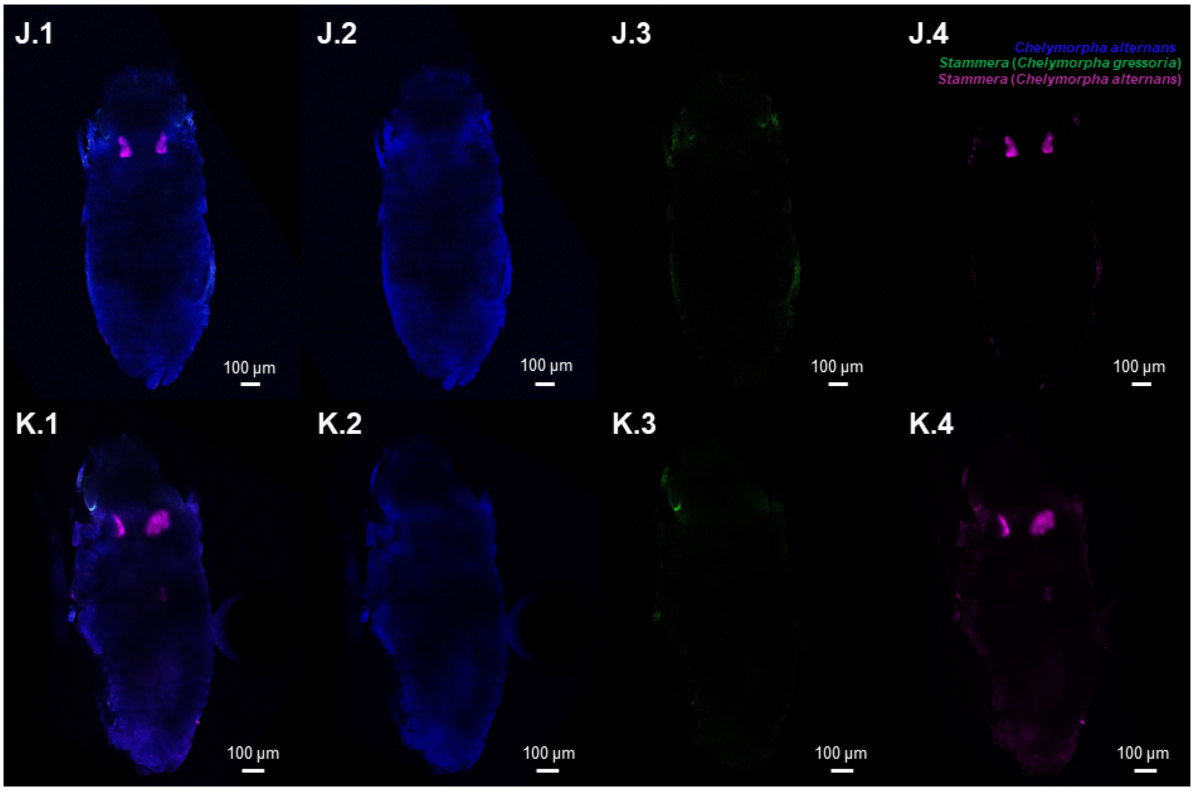


**Figure S1i** (J-K)**.** Fluorescence *in situ* hybridization (FISH) replicates on whole-mounts of *Chelymorpha alternans* embryos (re-infected). Probes used: *Chelymorpha alternans* host (blue: 18S rRNA), *Stammera* from *Chelymorpha gressoria* (green: 16SrRNA), and *Stammera* from *Chelymorpha alternans* (magenta: 16S rRNA). **(1)** correspond to the merged channels images while the others correspond to individual channel images: **(2)** host probe, **(3)** *Stammera* from *Chelymorpha gressoria* probe, and **(4)** *Stammera* from *Chelymorpha alternans* probe. Scale bars (100 µm) are included for reference.

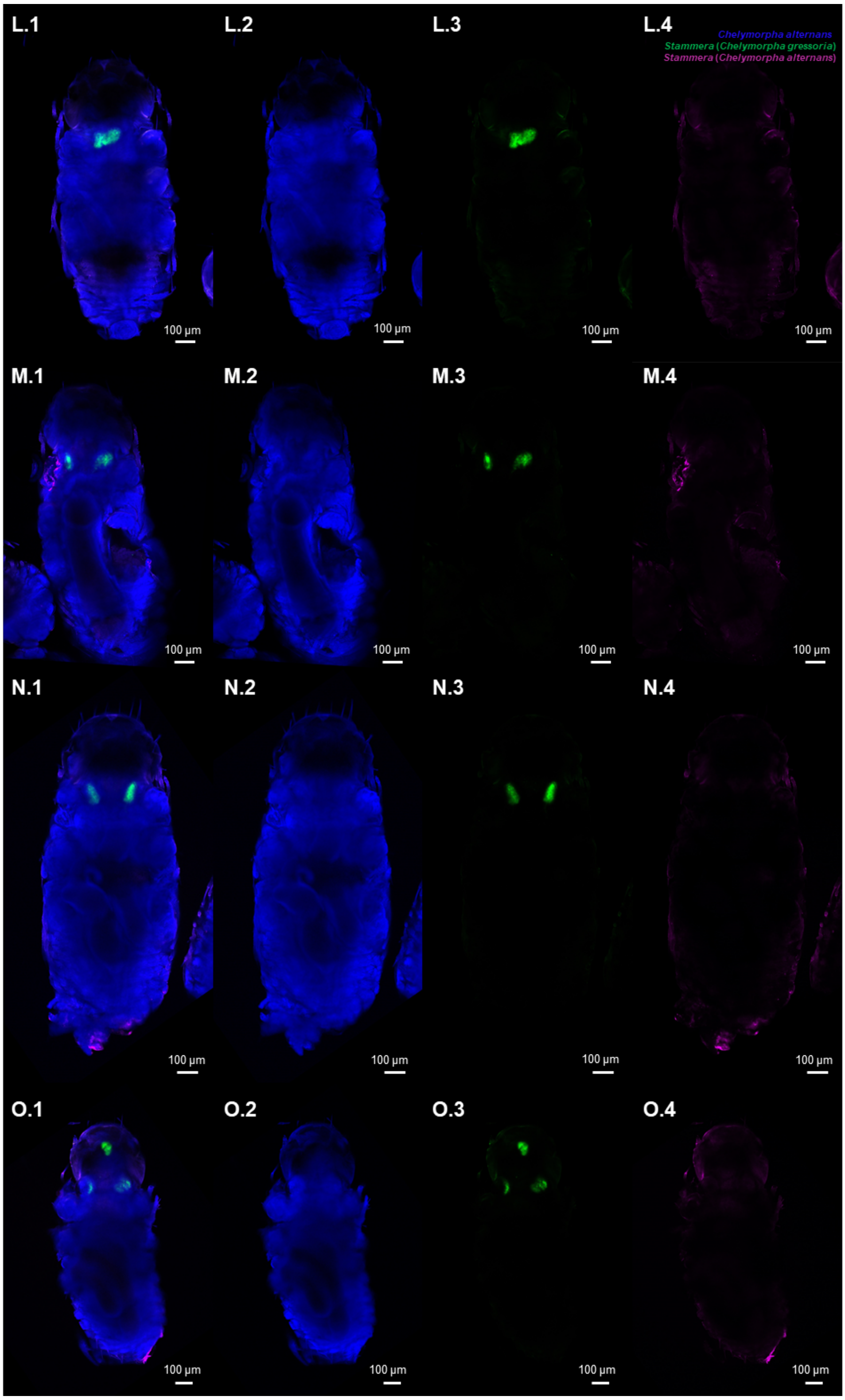


**Figure S1i** (L-O)**.** Fluorescence *in situ* hybridization (FISH) replicates on whole-mounts of *Chelymorpha alternans* embryos (cross-infected). Probes used: *Chelymorpha alternans* host (blue: 18S rRNA), *Stammera* from *Chelymorpha gressoria* (green: 16SrRNA), and *Stammera* from *Chelymorpha alternans* (magenta: 16S rRNA). **(1)** correspond to the merged channels images while the others correspond to individual channel images: **(2)** host probe, **(3)** *Stammera* from *Chelymorpha gressoria* probe, and **(4)** *Stammera* from *Chelymorpha alternans* probe. Scale bars (100 µm) are included for reference.

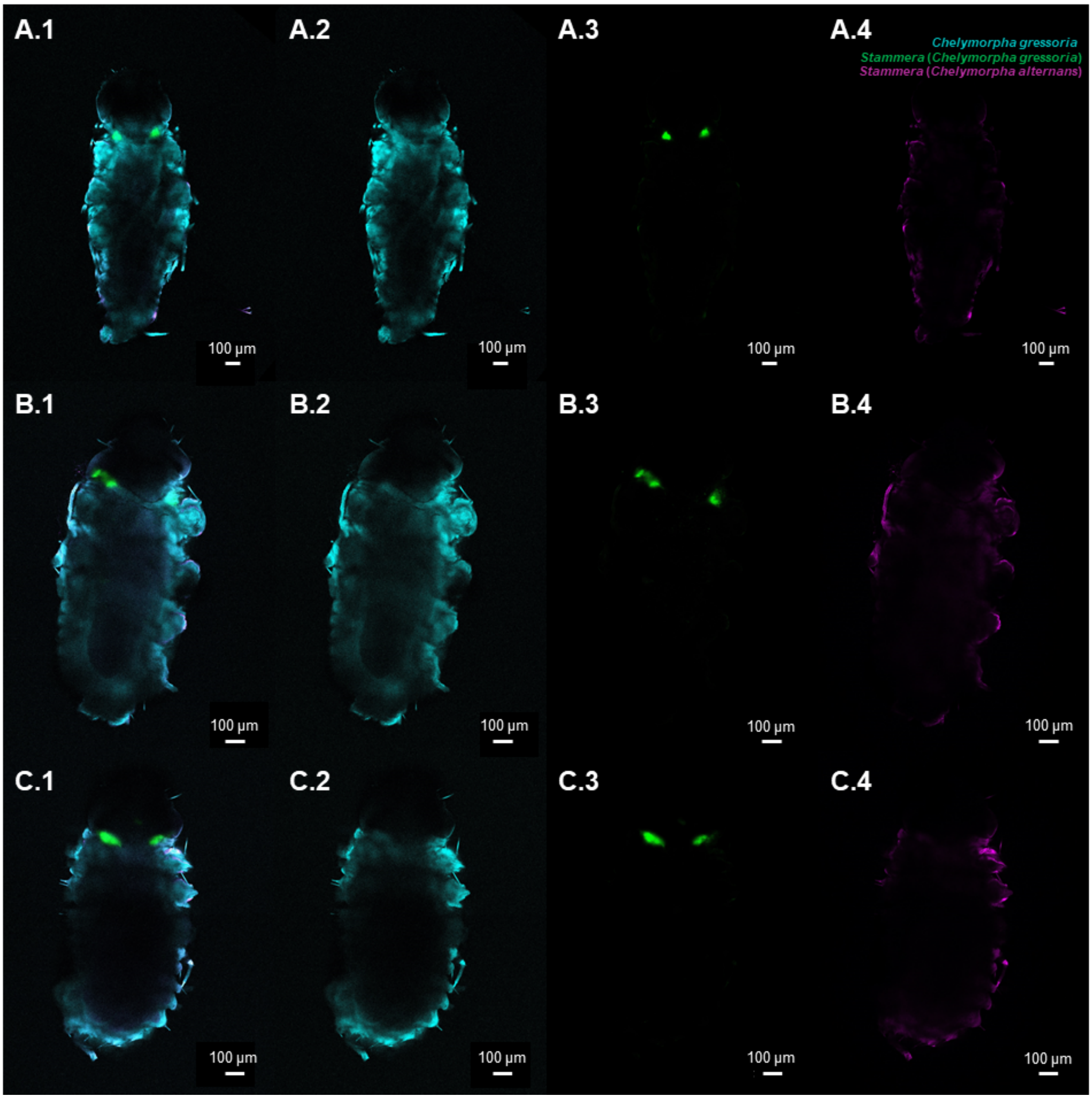


**Figure S1ii** (A-C)**.** Fluorescence *in situ* hybridization (FISH) replicates on whole-mounts of *Chelymorpha gressoria* embryos (untreated control). Probes used: *Chelymorpha gressoria* host (cyan: 18S rRNA), *Stammera* from *Chelymorpha gressoria* (green: 16SrRNA), and *Stammera* from *Chelymorpha alternans* (magenta: 16S rRNA). **(1)** correspond to the merged channels images while the others correspond to individual channel images: **(2)** host probe, **(3)** *Stammera* from *Chelymorpha gressoria* probe, and **(4)** *Stammera* from *Chelymorpha alternans* probe. Scale bars (100 µm) are included for reference.

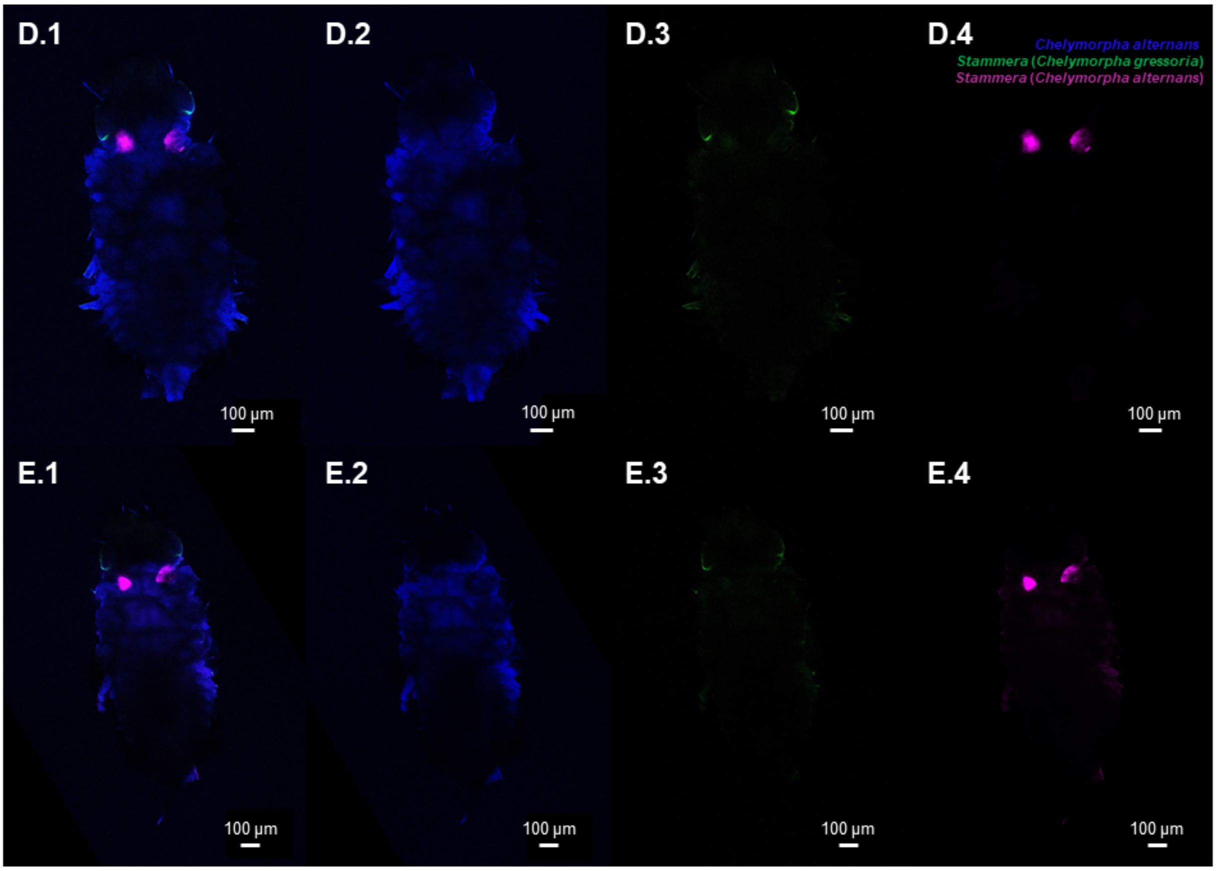


**Figure S1ii** (D-E)**.** Fluorescence *in situ* hybridization (FISH) replicates on whole-mounts of *Chelymorpha alternans* embryos (untreated control). Probes used: *Chelymorpha alternans* host (blue: 18S rRNA), *Stammera* from *Chelymorpha gressoria* (green: 16SrRNA), and *Stammera* from *Chelymorpha alternans* (magenta: 16S rRNA). **(1)** correspond to the merged channels images while the others correspond to individual channel images: **(2)** host probe, **(3)** *Stammera* from *Chelymorpha gressoria* probe, and **(4)** *Stammera* from *Chelymorpha alternans* probe. Scale bars (100 µm) are included for reference.

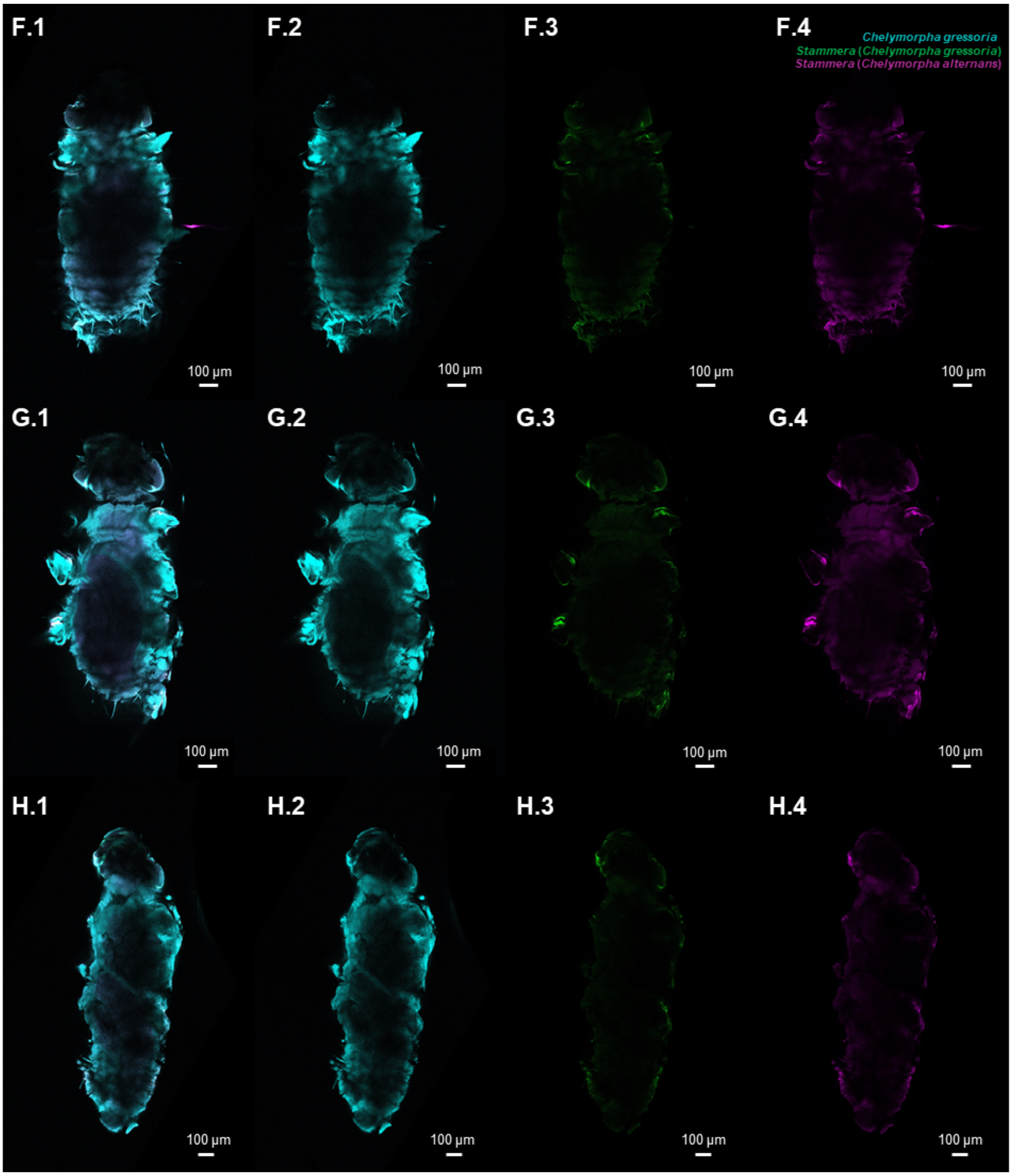


**Figure S1ii** (F-H)**.** Fluorescence *in situ* hybridization (FISH) replicates on whole-mounts of *Chelymorpha gressoria* embryos (aposymbiotic). Probes used: *Chelymorpha gressoria* host (cyan: 18S rRNA), *Stammera* from *Chelymorpha gressoria* (green: 16SrRNA), and *Stammera* from *Chelymorpha alternans* (magenta: 16S rRNA). **(1)** correspond to the merged channels images while the others correspond to individual channel images: **(2)** host probe, **(3)** *Stammera* from *Chelymorpha gressoria* probe, and **(4)** *Stammera* from *Chelymorpha alternans* probe. Scale bars (100 µm) are included for reference.

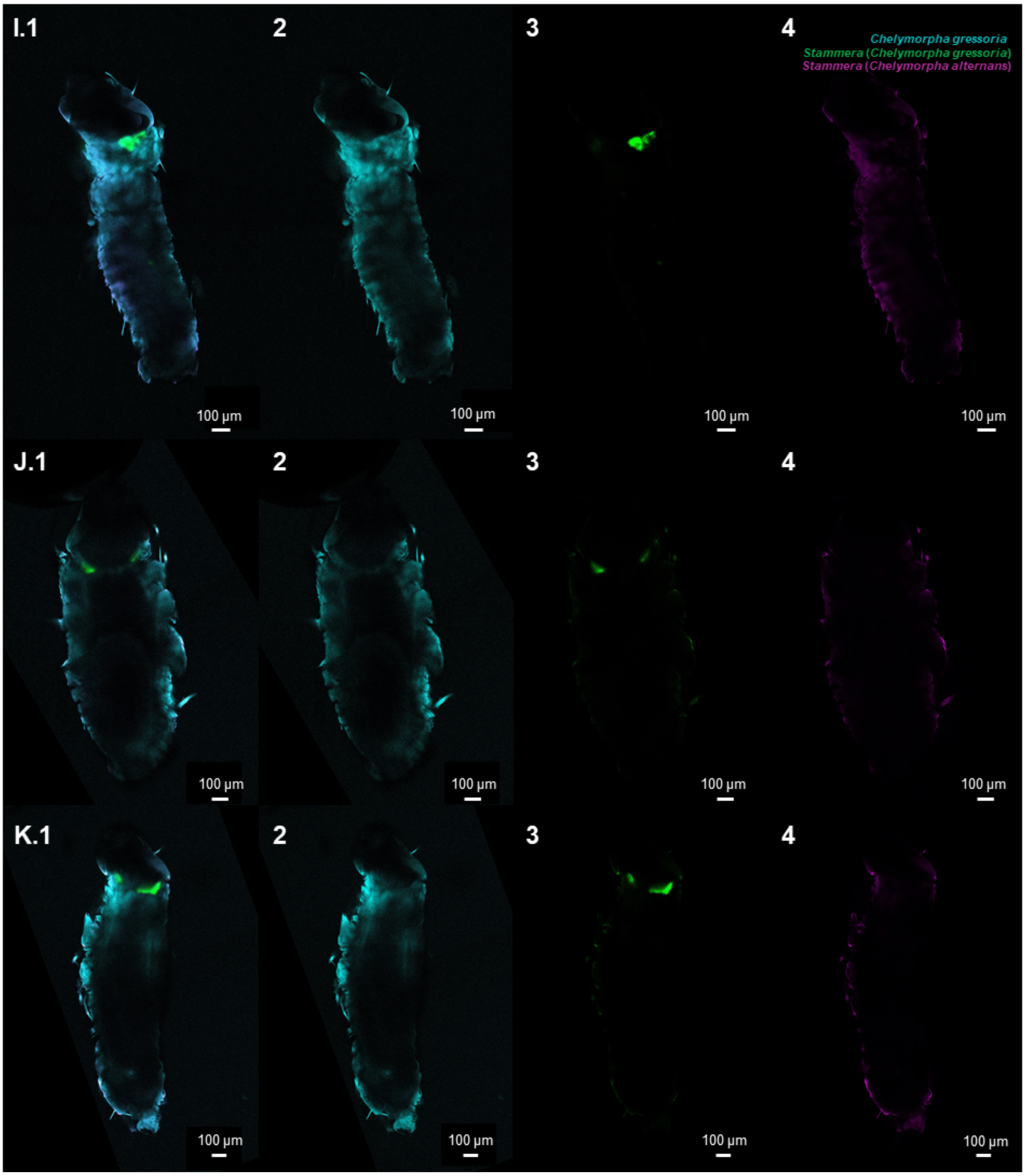


**Figure S1ii** (I-K)**.** Fluorescence *in situ* hybridization (FISH) replicates on whole-mounts of *Chelymorpha gressoria* embryos (re-infected). Probes used: *Chelymorpha gressoria* host (cyan: 18S rRNA), *Stammera* from *Chelymorpha gressoria* (green: 16SrRNA), and *Stammera* from *Chelymorpha alternans* (magenta: 16S rRNA). **(1)** correspond to the merged channels images while the others correspond to individual channel images: **(2)** host probe, **(3)** *Stammera* from *Chelymorpha gressoria* probe, and **(4)** *Stammera* from *Chelymorpha alternans* probe. Scale bars (100 µm) are included for reference.

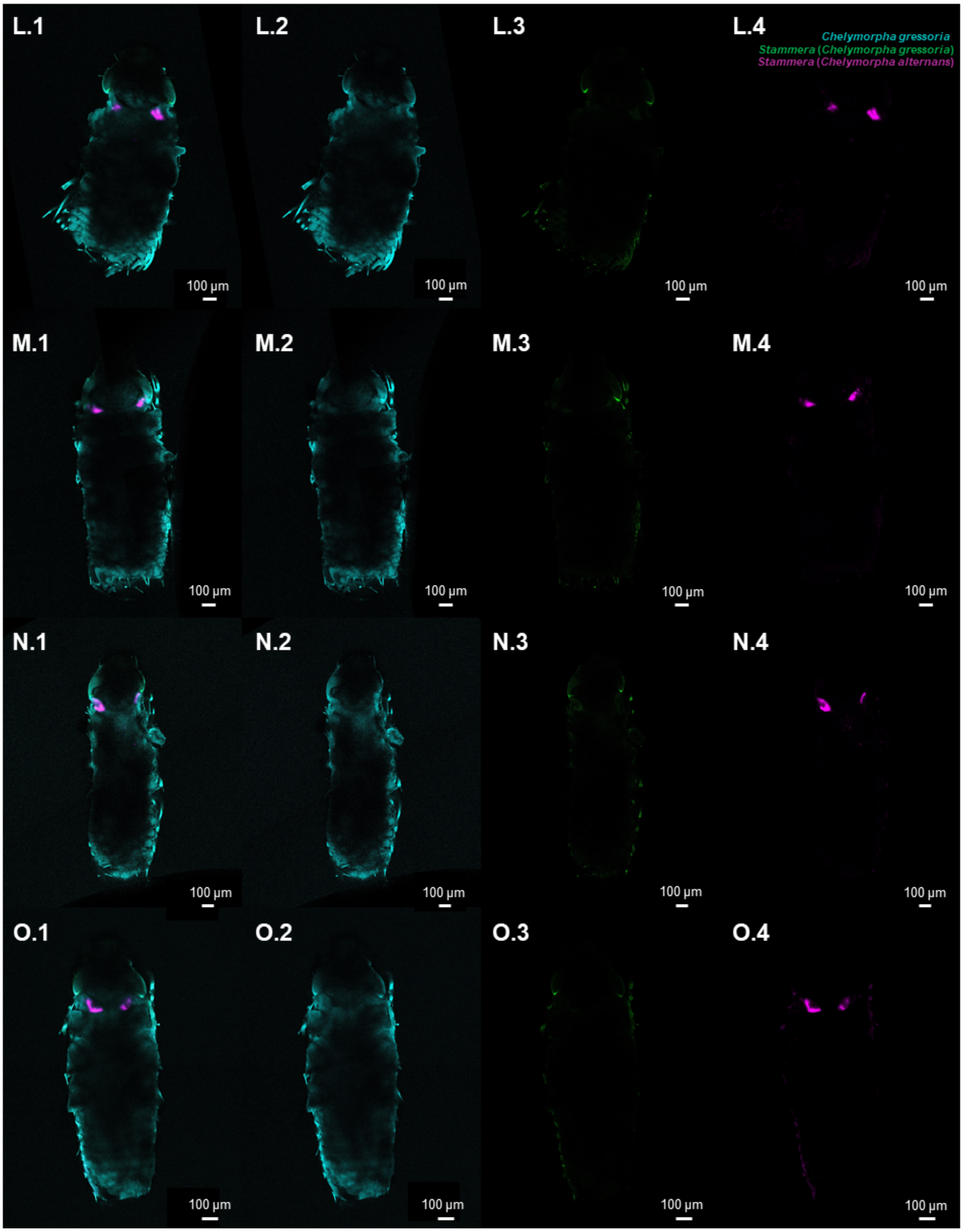


**Figure S1ii** (L-O)**.** Fluorescence *in situ* hybridization (FISH) replicates on whole-mounts of *Chelymorpha gressoria* embryos (cross-infected). Probes used: *Chelymorpha gressoria* host (cyan: 18S rRNA), *Stammera* from *Chelymorpha gressoria* (green: 16SrRNA), and *Stammera* from *Chelymorpha alternans* (magenta: 16S rRNA). **(1)** correspond to the merged channels images while the others correspond to individual channel images: **(2)** host probe, **(3)** *Stammera* from *Chelymorpha gressoria* probe, and **(4)** *Stammera* from *Chelymorpha alternans* probe. Scale bars (100 µm) are included for reference.

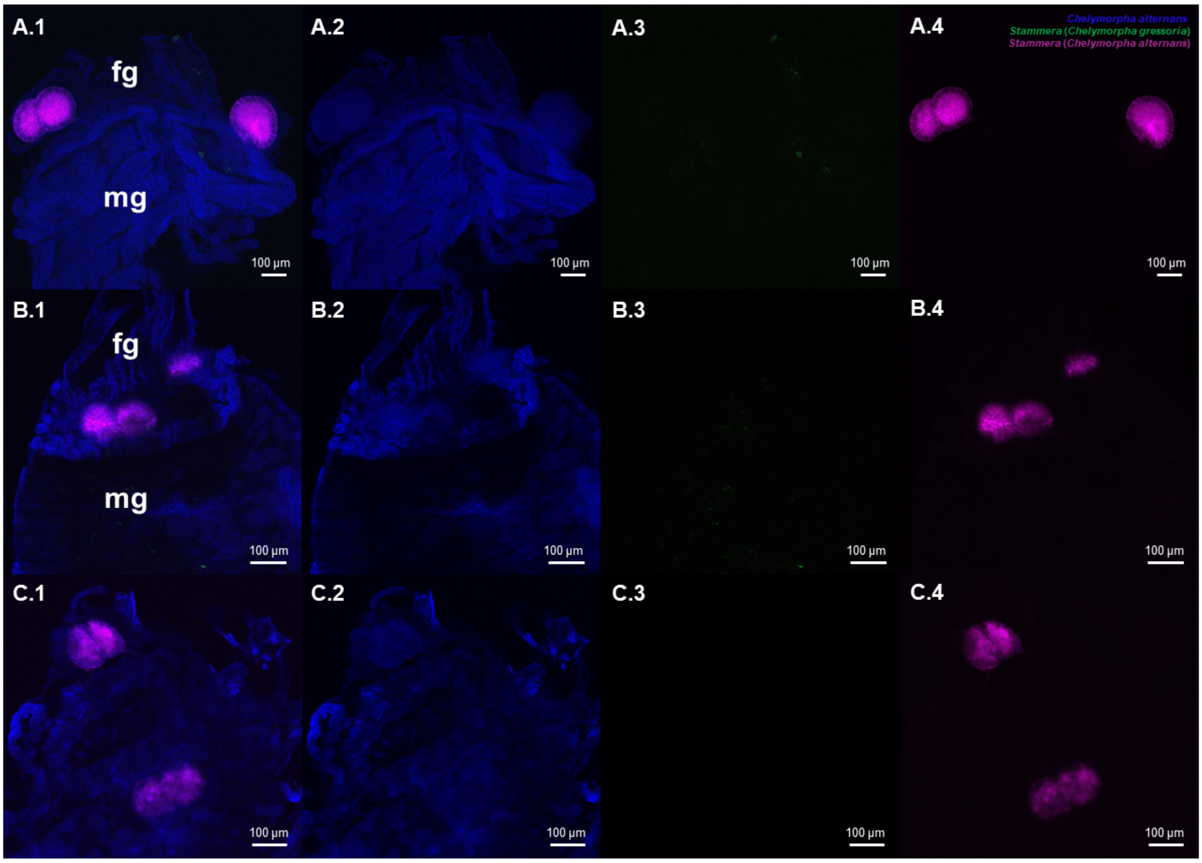


**Figure S1iii** (A-C)**.** Fluorescence *in situ* hybridization (FISH) replicates on whole-mounts of *Chelymorpha alternans* larvae foregut symbiotic organs (untreated control). Probes used: *Chelymorpha alternans* host (blue: 18S rRNA), *Stammera* from *Chelymorpha gressoria* (green: 16SrRNA), and *Stammera* from *Chelymorpha alternans* (magenta: 16S rRNA). **(1)** correspond to the merged channels images while the others correspond to individual channel images: **(2)** host probe, **(3)** *Stammera* from *Chelymorpha gressoria* probe, and **(4)** *Stammera* from *Chelymorpha alternans* probe. Abbreviations: fg, foregut; mg, midgut. Scale bars (100 µm) are included for reference.

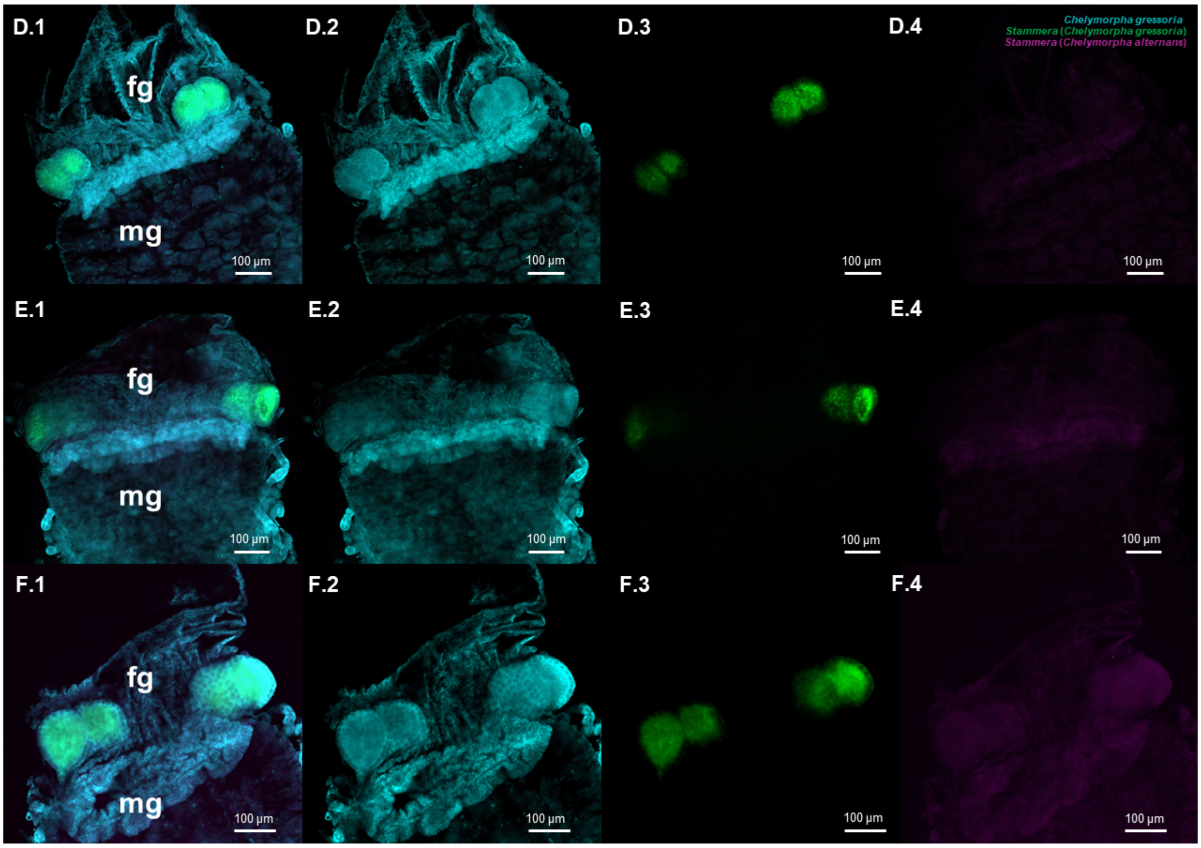


**Figure S1iii** (D-F)**.** Fluorescence *in situ* hybridization (FISH) replicates on whole-mounts of *Chelymorpha gressoria* larvae foregut symbiotic organs (untreated control). Probes used: *Chelymorpha gressoria* host (cyan: 18S rRNA), *Stammera* from *Chelymorpha gressoria* (green: 16SrRNA), and *Stammera* from *Chelymorpha alternans* (magenta: 16S rRNA). **(1)** correspond to the merged channels images while the others correspond to individual channel images: **(2)** host probe, **(3)** *Stammera* from *Chelymorpha gressoria* probe, and **(4)** *Stammera* from *Chelymorpha alternans* probe. Abbreviations: fg, foregut; mg, midgut. Scale bars (100 µm) are included for reference.

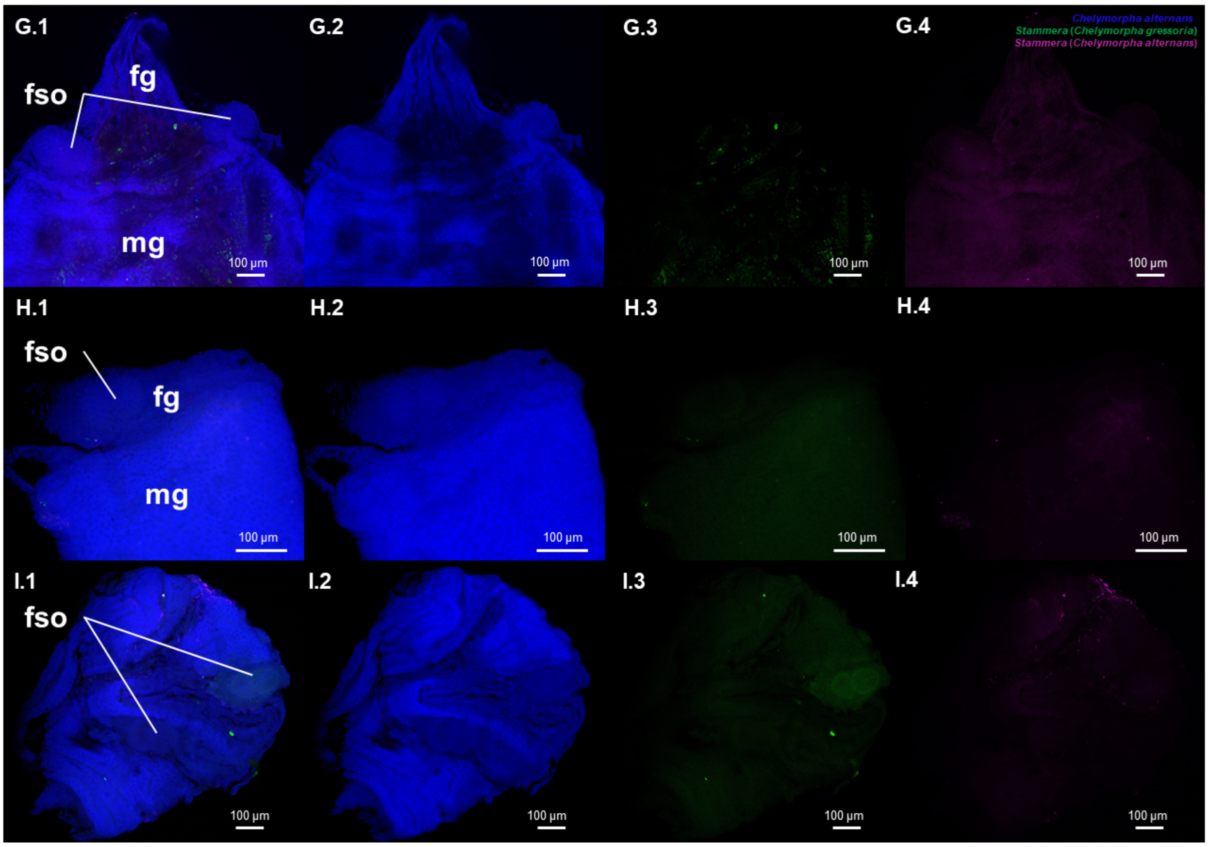


**Figure S1iii** (G-I**)**. Fluorescence *in situ* hybridization (FISH) replicates on whole-mounts of *Chelymorpha alternans* larvae foregut symbiotic organs (aposymbiotic). Probes used: *Chelymorpha alternans* host (blue: 18S rRNA), *Stammera* from *Chelymorpha gressoria* (green: 16SrRNA), and *Stammera* from *Chelymorpha alternans* (magenta: 16S rRNA). **(1)** correspond to the merged channels images while the others correspond to individual channel images: **(2)** host probe, **(3)** *Stammera* from *Chelymorpha gressoria* probe, and **(4)** *Stammera* from *Chelymorpha alternans* probe. Abbreviations: fg, foregut; mg, midgut; fso, foregut symbiotic organs. Scale bars (100 µm) are included for reference.

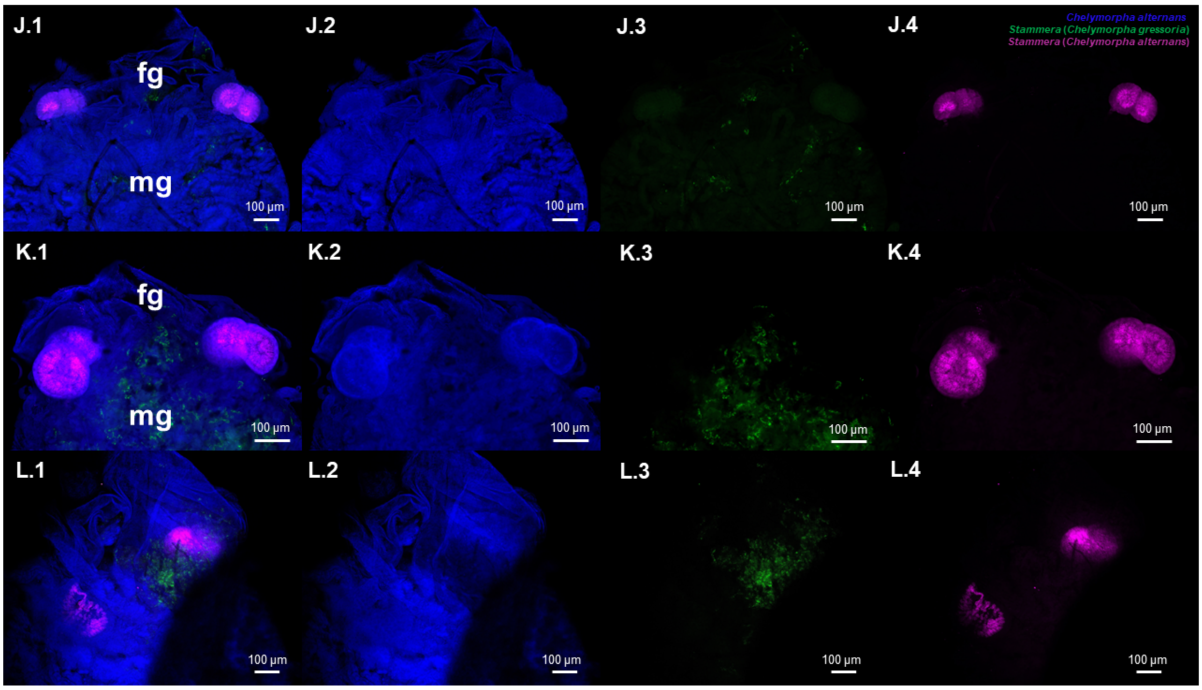


**Figure S1iii** (J-L)**.** Fluorescence *in situ* hybridization (FISH) replicates on whole-mounts of *Chelymorpha alternans* larvae foregut symbiotic organs (re-infected). Probes used: *Chelymorpha alternans* host (blue: 18S rRNA), *Stammera* from *Chelymorpha gressoria* (green: 16SrRNA), and *Stammera* from *Chelymorpha alternans* (magenta: 16S rRNA). **(1)** correspond to the merged channels images while the others correspond to individual channel images: **(2)** host probe, **(3)** *Stammera* from *Chelymorpha gressoria* probe, and **(4)** *Stammera* from *Chelymorpha alternans* probe. Abbreviations: fg, foregut; mg, midgut. Scale bars (100 µm) are included for reference.

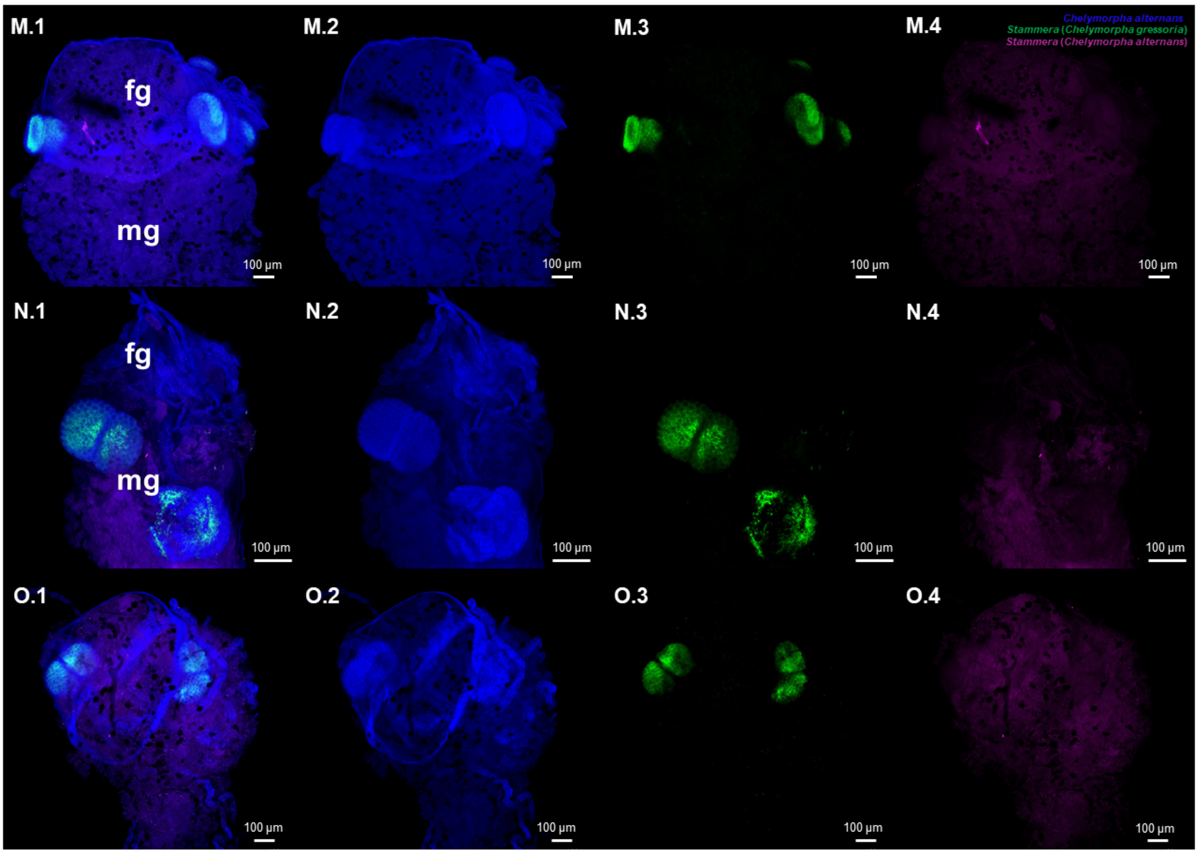


**Figure S1iii** (M-O)**.** Fluorescence *in situ* hybridization (FISH) replicates on whole-mounts of *Chelymorpha alternans* larvae foregut symbiotic organs (cross-infected). Probes used: *Chelymorpha alternans* host (blue: 18S rRNA), *Stammera* from *Chelymorpha gressoria* (green: 16SrRNA), and *Stammera* from *Chelymorpha alternans* (magenta: 16S rRNA). **(1)** correspond to the merged channels images while the others correspond to individual channel images: **(2)** host probe, **(3)** *Stammera* from *Chelymorpha gressoria* probe, and **(4)** *Stammera* from *Chelymorpha alternans* probe. Abbreviations: fg, foregut; mg, midgut. Scale bars (100 µm) are included for reference.

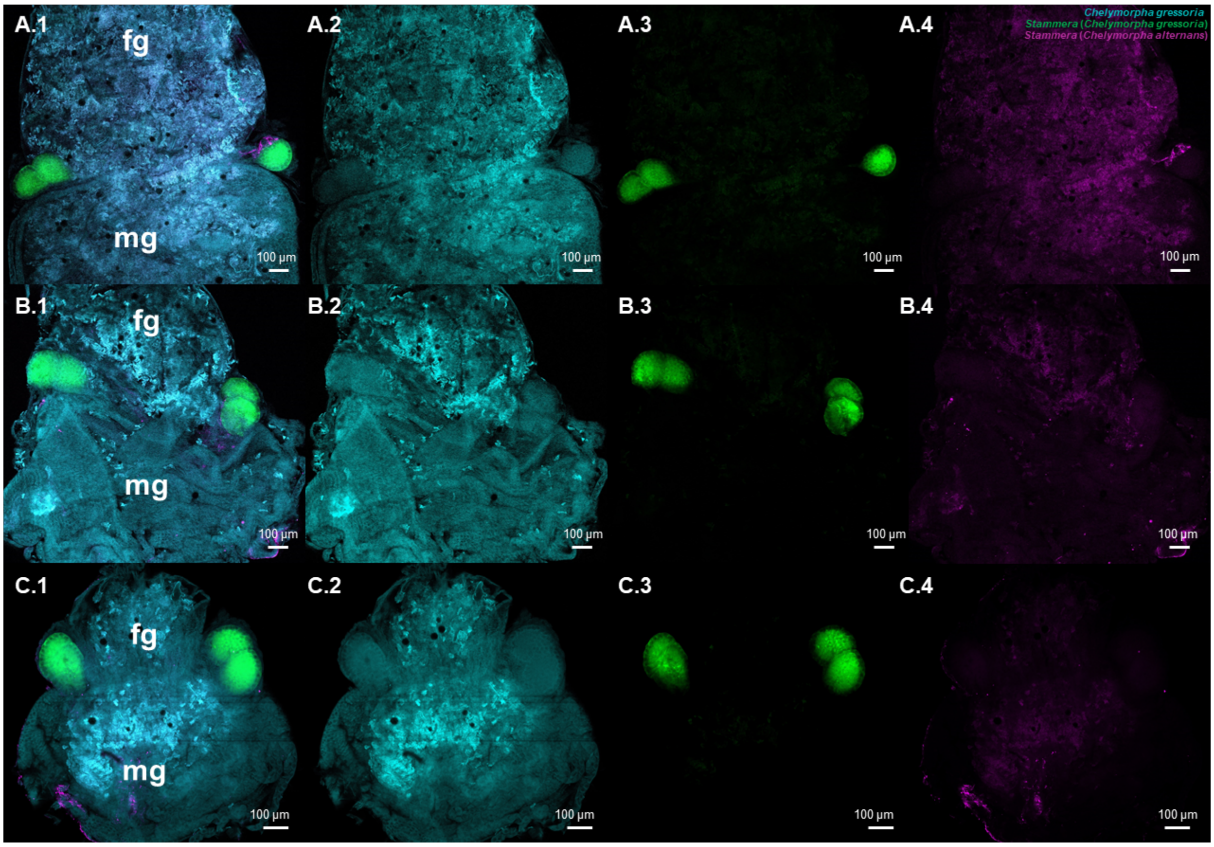


**Figure S1iv** (A-C)**.** Fluorescence *in situ* hybridization (FISH) replicates on whole-mounts of *Chelymorpha gressoria* larvae foregut symbiotic organs (untreated control). Probes used: *Chelymorpha gressoria* host (cyan: 18S rRNA), *Stammera* from *Chelymorpha gressoria* (green: 16SrRNA), and *Stammera* from *Chelymorpha alternans* (magenta: 16S rRNA). **(1)** correspond to the merged channels images while the others correspond to individual channel images: **(2)** host probe, **(3)** *Stammera* from *Chelymorpha gressoria* probe, and **(4)** *Stammera* from *Chelymorpha alternans* probe. Abbreviations: fg, foregut; mg, midgut. Scale bars (100 µm) are included for reference.

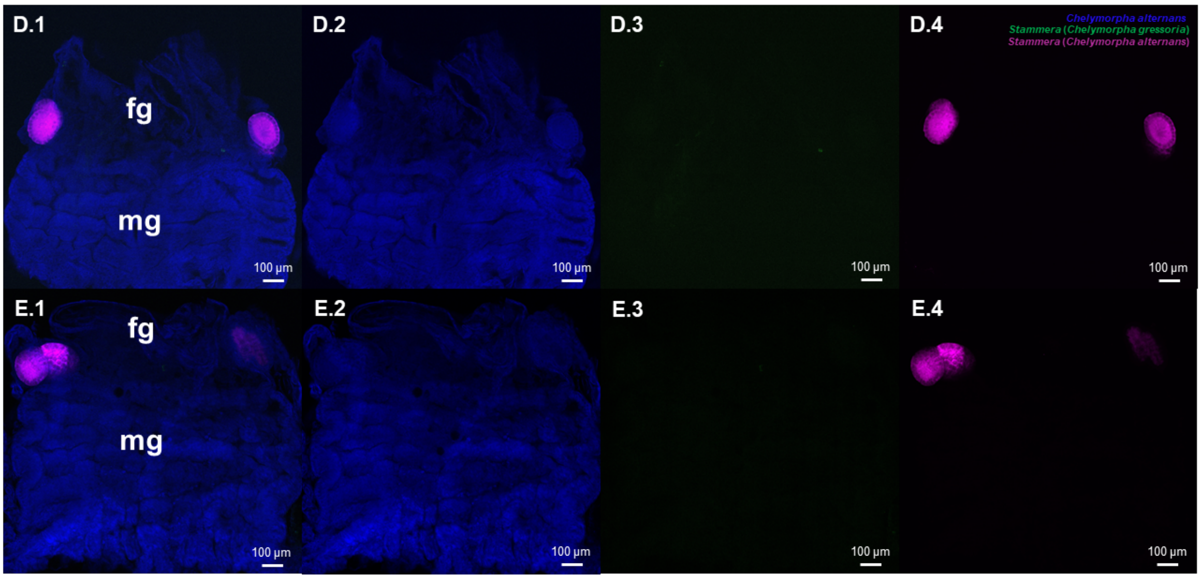


**Figure S1iv** (D-E)**.** Fluorescence *in situ* hybridization (FISH) replicates on whole-mounts of *Chelymorpha alternans* larvae foregut symbiotic organs (untreated control). Probes used: *Chelymorpha alternans* host (blue: 18S rRNA), *Stammera* from *Chelymorpha gressoria* (green: 16SrRNA), and *Stammera* from *Chelymorpha alternans* (magenta: 16S rRNA). **(1)** correspond to the merged channels images while the others correspond to individual channel images: **(2)** host probe, **(3)** *Stammera* from *Chelymorpha gressoria* probe, and **(4)** *Stammera* from *Chelymorpha alternans* probe. Abbreviations: fg, foregut; mg, midgut. Scale bars (100 µm) are included for reference.

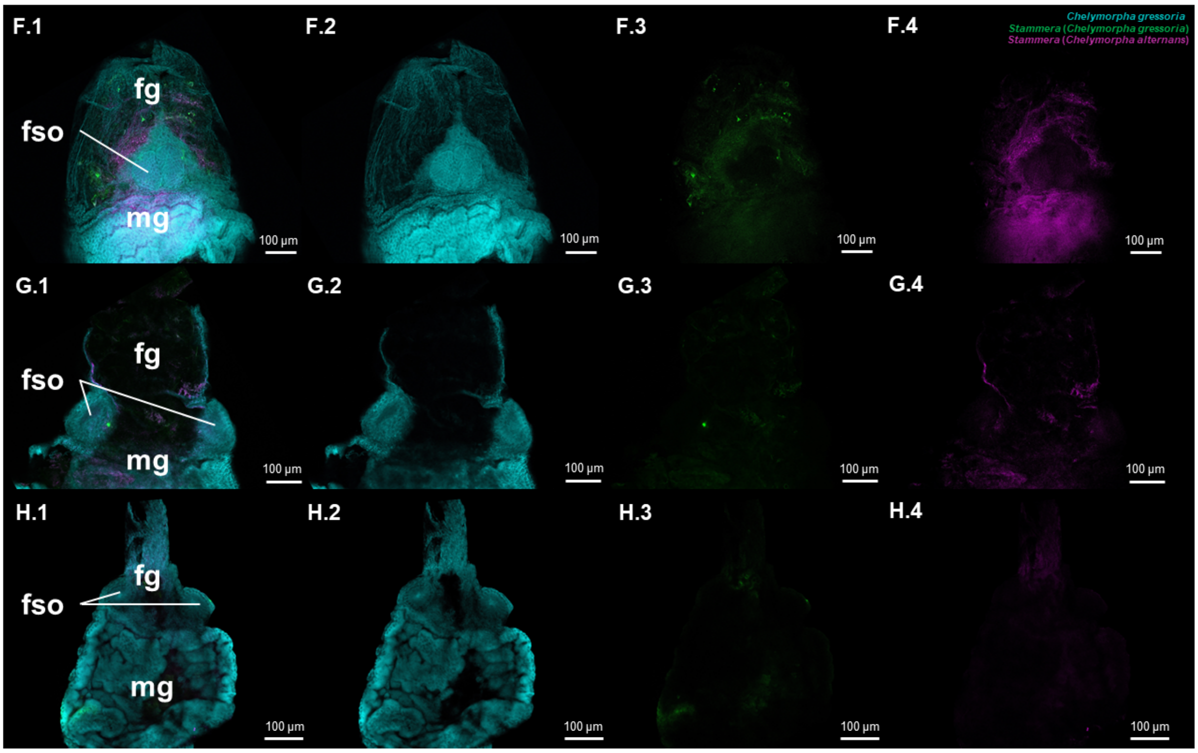


**Figure S1iv** (F-H)**.** Fluorescence *in situ* hybridization (FISH) replicates on whole-mounts of *Chelymorpha gressoria* larvae foregut symbiotic organs (aposymbiotic). Probes used: *Chelymorpha gressoria* host (cyan: 18S rRNA), *Stammera* from *Chelymorpha gressoria* (green: 16SrRNA), and *Stammera* from *Chelymorpha alternans* (magenta: 16S rRNA). **(1)** correspond to the merged channels images while the others correspond to individual channel images: **(2)** host probe, **(3)** *Stammera* from *Chelymorpha gressoria* probe, and **(4)** *Stammera* from *Chelymorpha alternans* probe. Abbreviations: fg, foregut; mg, midgut; fso, foregut symbiotic organs. Scale bars (100 µm) are included for reference.

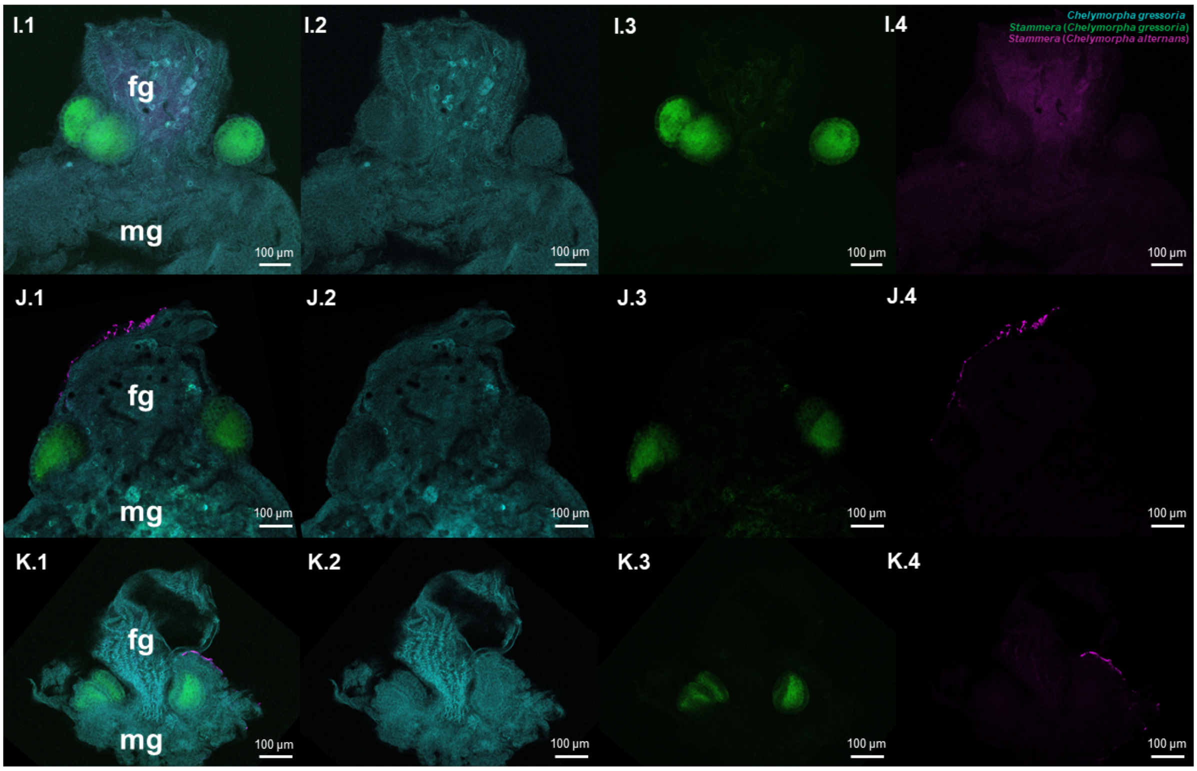


**Figure S1iv** (I-K)**.** Fluorescence *in situ* hybridization (FISH) replicates on whole-mounts of *Chelymorpha gressoria* larvae foregut symbiotic organs (re-infected). Probes used: *Chelymorpha gressoria* host (cyan: 18S rRNA), *Stammera* from *Chelymorpha gressoria* (green: 16SrRNA), and *Stammera* from *Chelymorpha alternans* (magenta: 16S rRNA). **(1)** correspond to the merged channels images while the others correspond to individual channel images: **(2)** host probe, **(3)** *Stammera* from *Chelymorpha gressoria* probe, and **(4)** *Stammera* from *Chelymorpha alternans* probe. Abbreviations: fg, foregut; mg, midgut. Scale bars (100 µm) are included for reference.

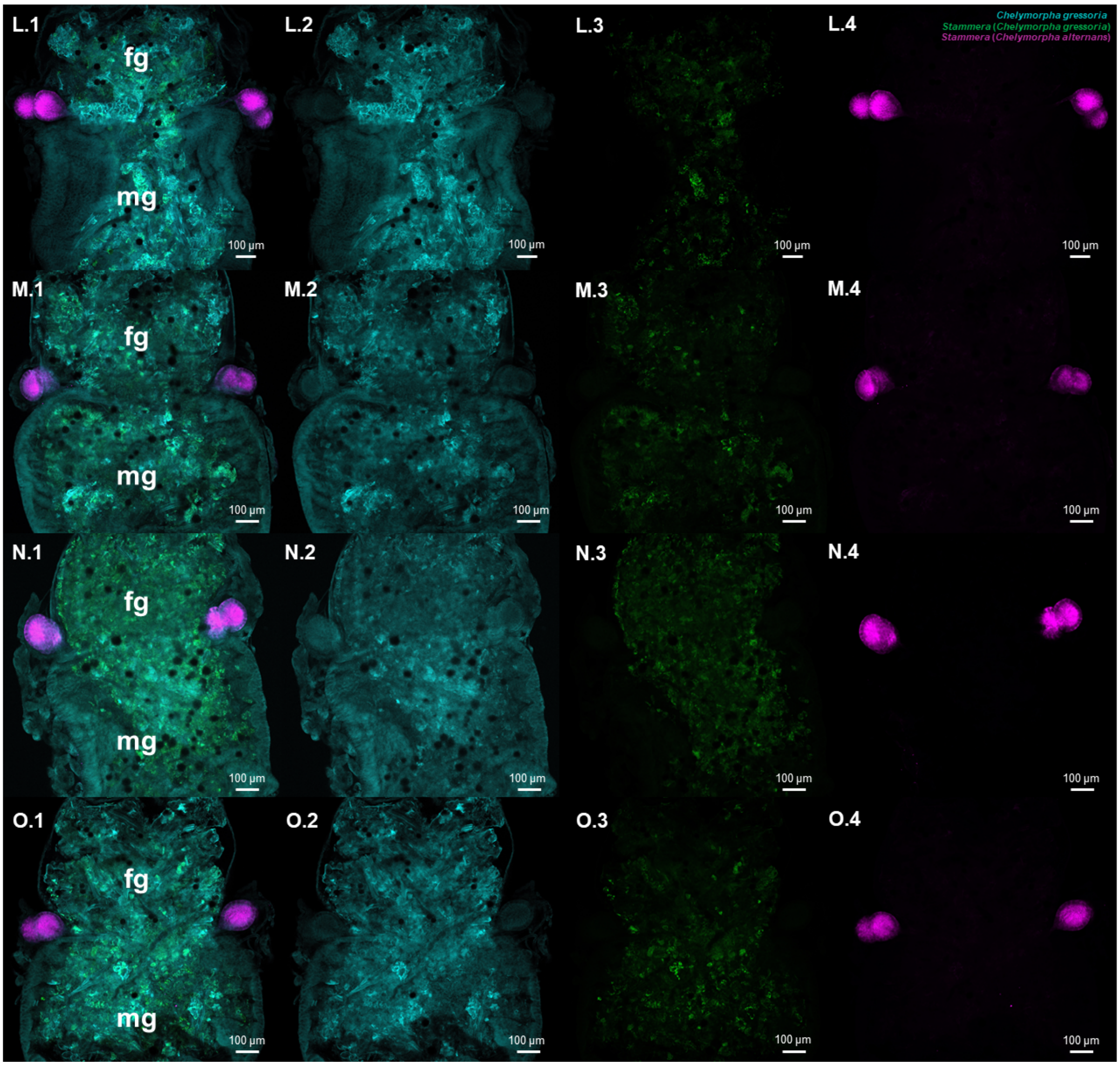


**Figure S1iv** (L-O)**.** Fluorescence *in situ* hybridization (FISH) replicates on whole-mounts of *Chelymorpha gressoria* larvae foregut symbiotic organs (cross-infected). Probes used: *Chelymorpha gressoria* host (cyan: 18S rRNA), *Stammera* from *Chelymorpha gressoria* (green: 16SrRNA), and *Stammera* from *Chelymorpha alternans* (magenta: 16S rRNA). **(1)** correspond to the merged channels images while the others correspond to individual channel images: **(2)** host probe, **(3)** *Stammera* from *Chelymorpha gressoria* probe, and **(4)** *Stammera* from *Chelymorpha alternans* probe. Abbreviations: fg, foregut; mg, midgut. Scale bars (100 µm) are included for reference.

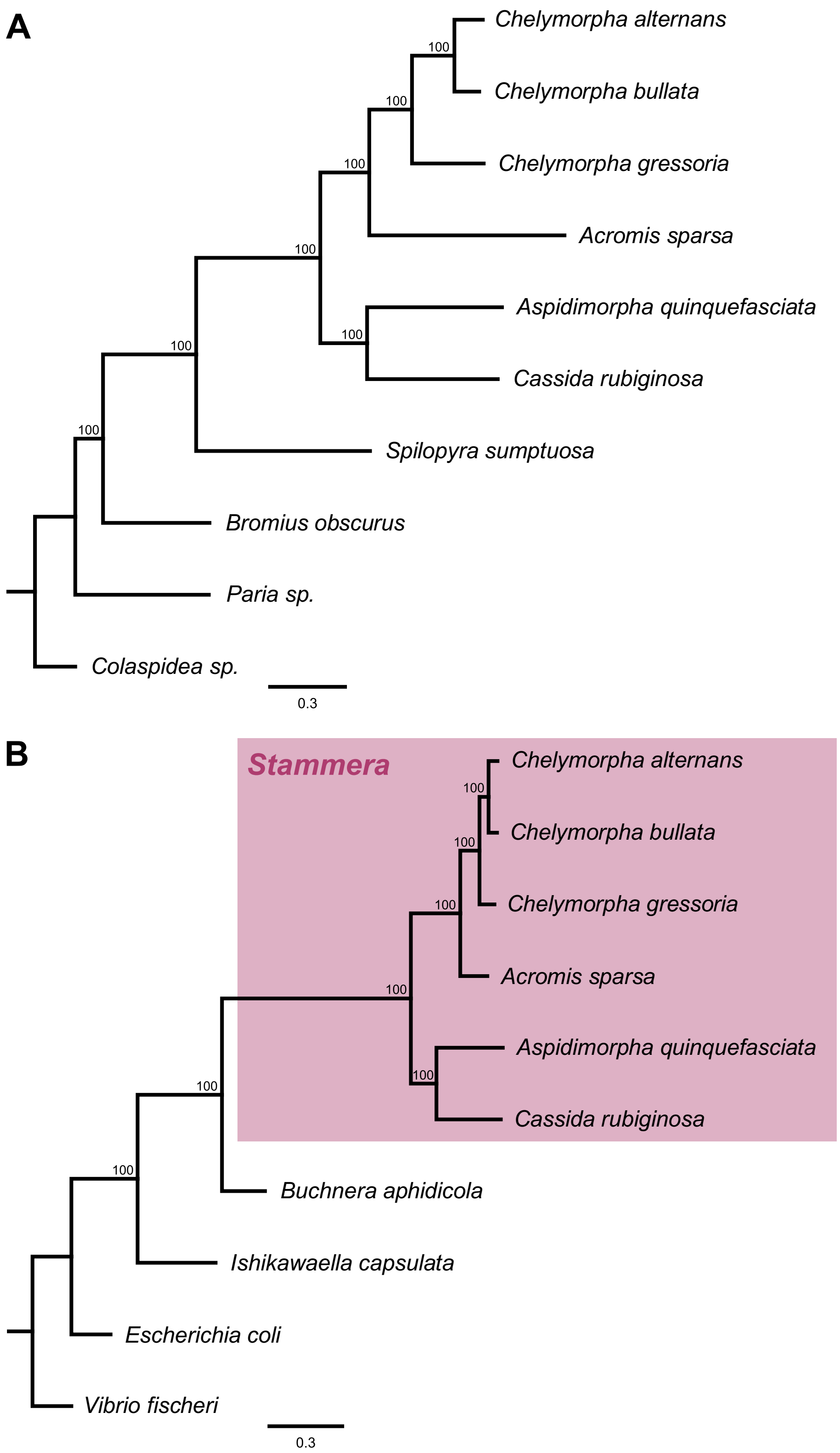


**Figure S2. (A)** Detailed host phylogeny including outgroups based on 15 mitochondrial genes. The phylogenetic tree was constructed by maximum likelihood (ML) methods. **(B)** Detailed *Stammera* phylogeny including outgroups based on maximum likelihood (ML) methods. The phylogenomic tree was constructed based on a concatenated alignment of 61 single-copy core genes in RAxML, using the most appropriate substitution model according to PartitionFinder2. Bootstrap support values are shown for each node.

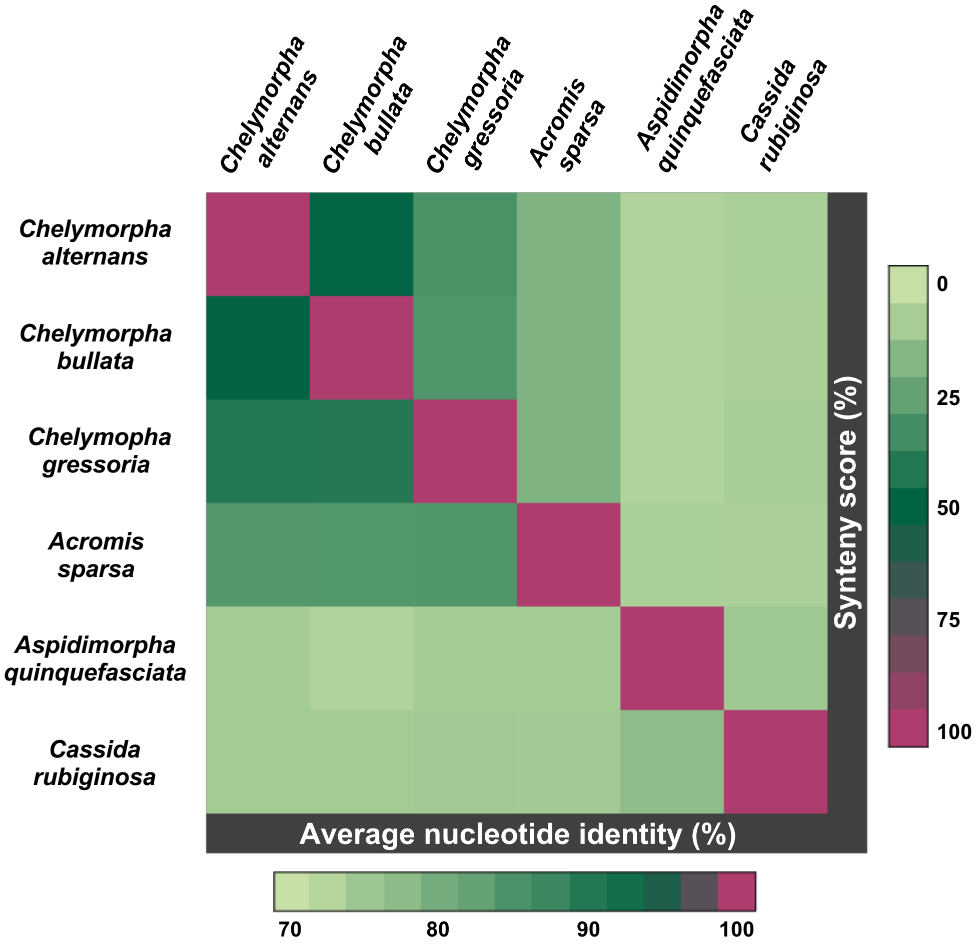


**Figure S3.** Heatmap illustrating genetic distances between *Stammera* symbionts. Pairwise comparisons of average nucleotide identity (ANI) (light green, 70%; dark green, 95%) and synteny scores (light green, 0%; dark green, 60%) are shown.

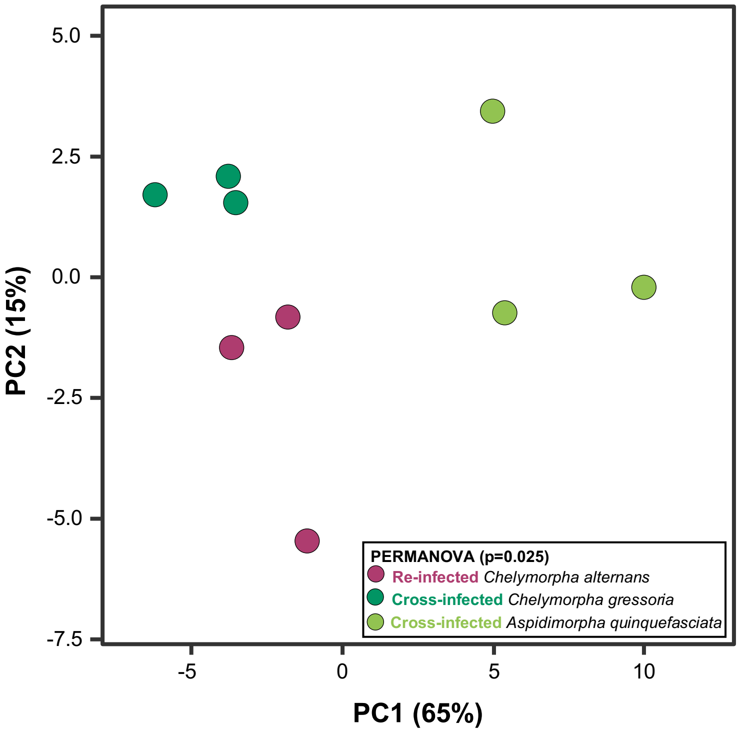


**Figure S4.** Principal coordinate analysis (PCA) of host differentially expressed genes after colonization of foregut symbiotic organs by *Stammera* symbionts. Significant clustering was assessed by PERMANOVA (after removing the batch effects) based on Euclidean distances between samples ($p$ = 0.025, Table S4A).

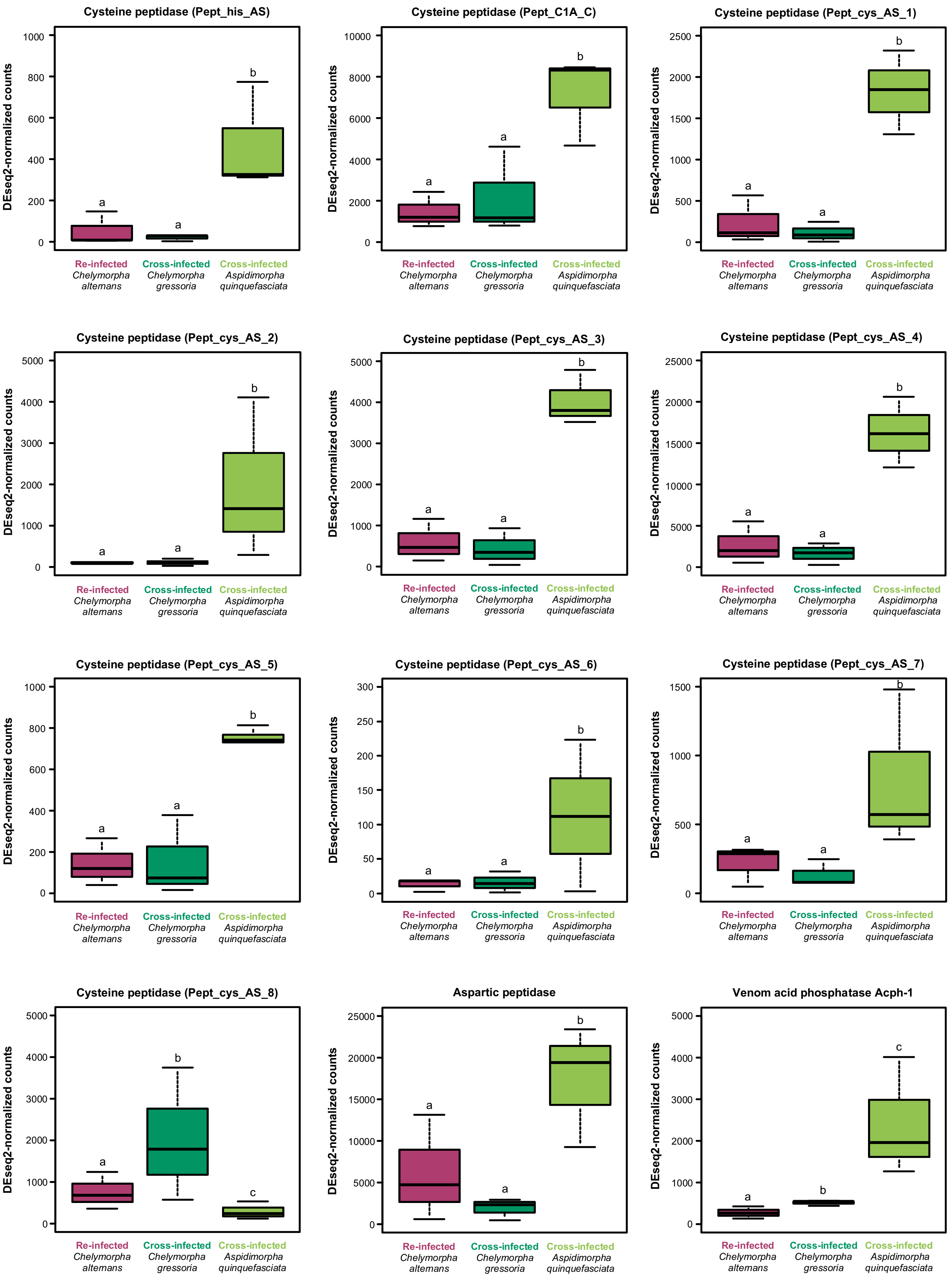

Cysteine peptidase (Pept_his_AS)
DEseq2-normalized counts
0
200
400
600
800
1000
a
a
b
Cysteine peptidase (Pept_C1A_C)
DEseq2-normalized counts
0
2000
4000
6000
8000
10000
a
a
b
Cysteine peptidase (Pept_cys_AS_1)
DEseq2-normalized counts
0
500
1000
1500
2000
2500
a
a
b
Cysteine peptidase (Pept_cys_AS_2)
DEseq2-normalized counts
0
1000
2000
3000
4000
5000
a
a
b
Cysteine peptidase (Pept_cys_AS_3)
DEseq2-normalized counts
0
1000
2000
3000
4000
5000
a
a
b
Cysteine peptidase (Pept_cys_AS_4)
DEseq2-normalized counts
0
5000
10000
15000
20000
25000
a
a
b
Cysteine peptidase (Pept_cys_AS_5)
DEseq2-normalized counts
0
200
400
600
800
1000
a
a
b
Cysteine peptidase (Pept_cys_AS_6)
DEseq2-normalized counts
0
50
100
150
200
250
300
a
a
b
Cysteine peptidase (Pept_cys_AS_7)
DEseq2-normalized counts
0
500
1000
1500
a
a
b
Cysteine peptidase (Pept_cys_AS_8)
DEseq2-normalized counts
0
1000
2000
3000
4000
5000
a
b
c
Aspartic peptidase
DEseq2-normalized counts
0
5000
10000
15000
20000
25000
a
a
b
Venom acid phosphatase Acph-1
DEseq2-normalized counts
0
1000
2000
3000
4000
5000
a
b
c
Re-infected Chelymorpha altemans
Cross-infected Chelymorpha gressoria
Cross-infected Aspidimorpha quinquefasciata

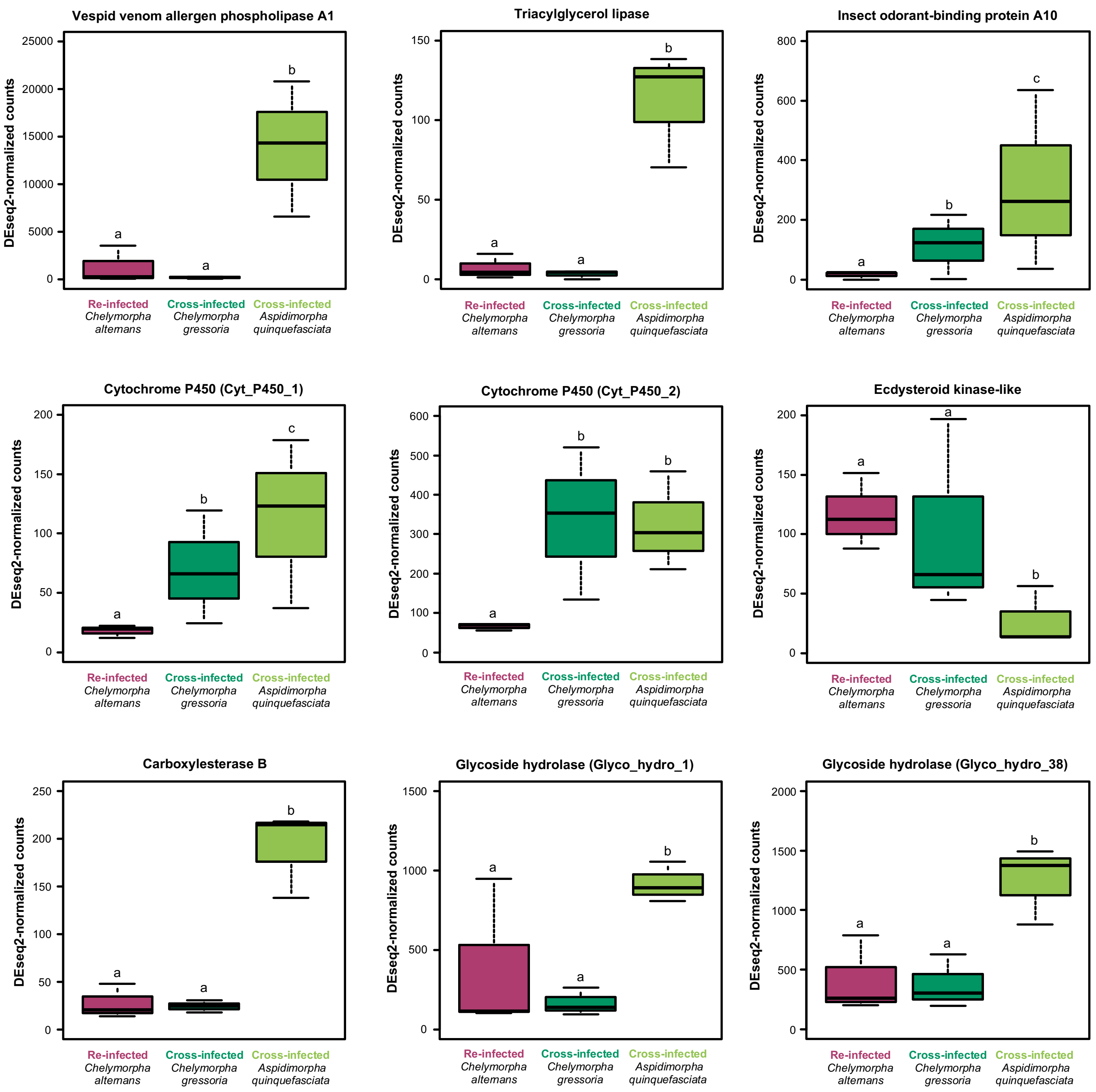


**Figure S5.** Differential expression of host genes after colonization of foregut symbiotic organs by the native symbiont (magenta), non-native symbiont of *Chelymorpha gressoria* (dark green), or non-native symbiont of *Aspidimorpha quinquefasciata* (light green). Counts were normalized by DESeq2's median of ratios. Lines represent medians, boxes indicate 25-75 percentiles, and whiskers denote range. Different letters indicated significant differences (See Table S4B).

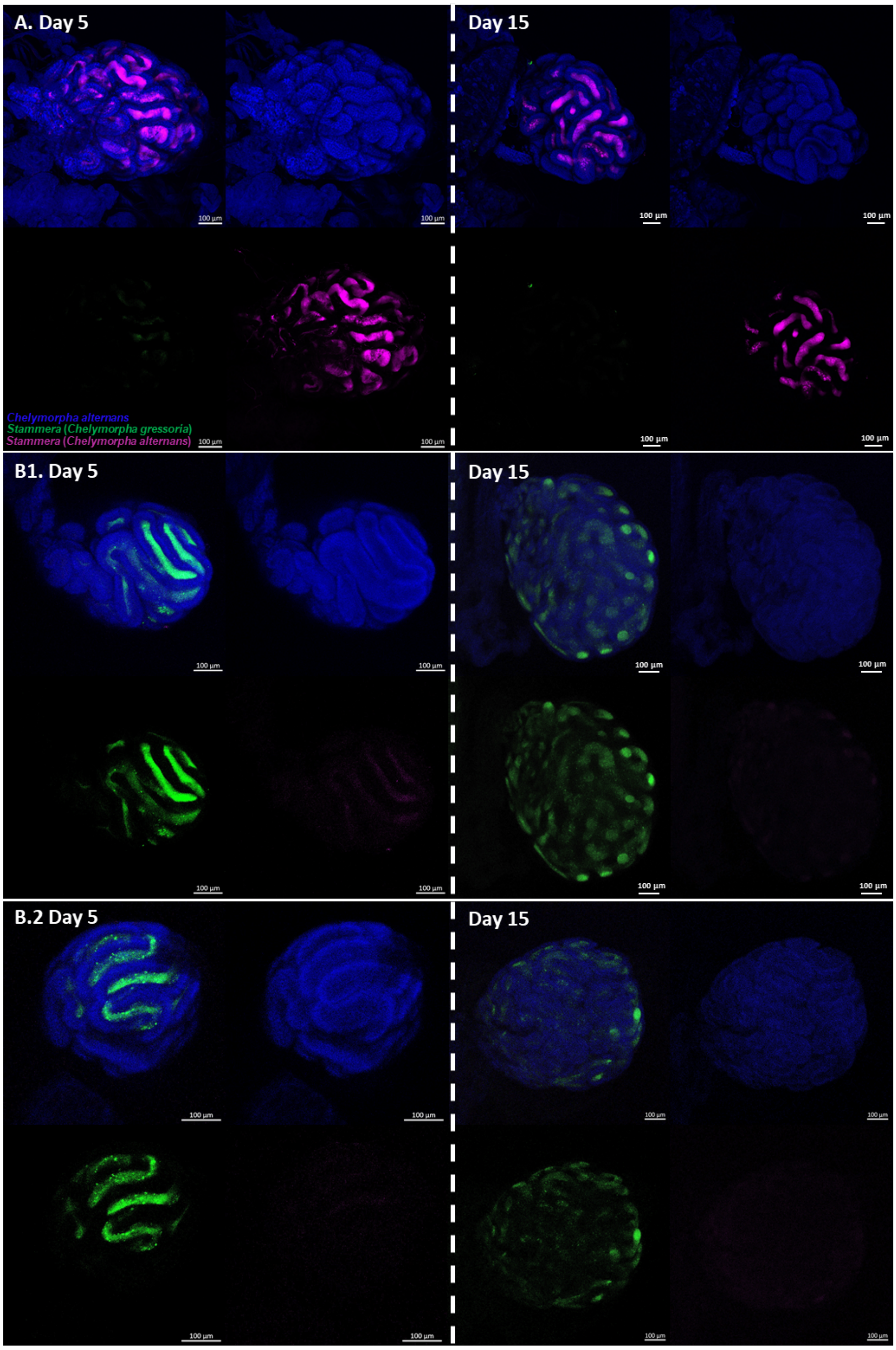


**Figure S6.** Fluorescence *in situ* hybridization (FISH) on whole-mounts of ovary-associated glands from *Chelymorpha alternans* females **(A)** re-infected with native symbiont or **(B)** cross-infected with non-native symbiont at 5 and 15 days after emergence. Both glands of the cross-infected sample were examined. Probes used: *Chelymorpha alternans* host (blue: 18S rRNA), *Stammera* from *Chelymorpha gressoria* (green: 16SrRNA), and *Stammera* from *Chelymorpha alternans* (magenta: 16S rRNA). The top left image corresponds to the merged channels images while the others correspond to individual channel images: top right image for the host probe, bottom left image for the probe targeting *Stammera* from *Chelymorpha gressoria*, and bottom right image for the probe targeting *Stammera* from *Chelymorpha alternans*. Scale bars (100 µm) are included for reference.

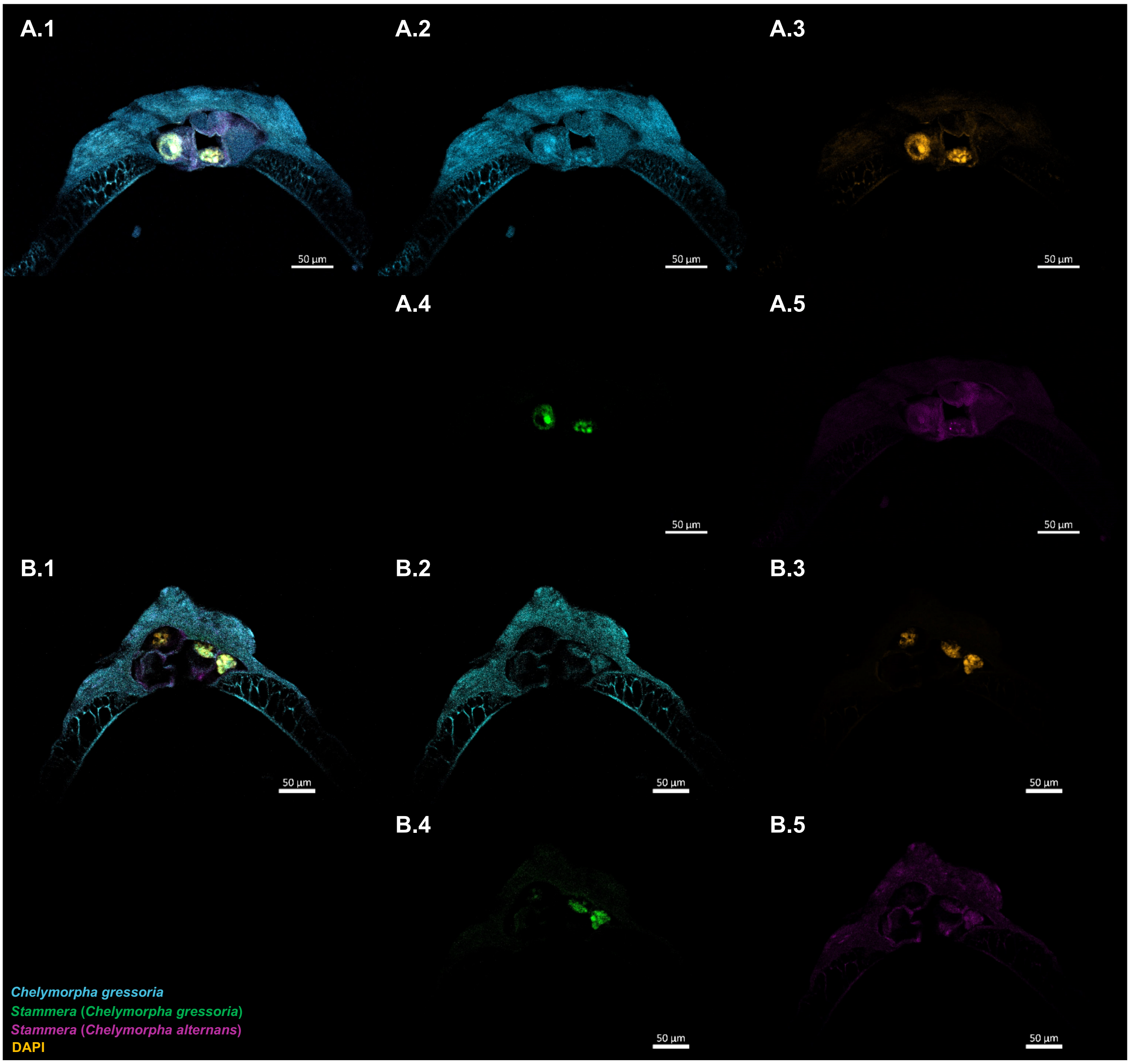


**Figure S7. (A-B)** Fluorescence *in situ* hybridization (FISH) replicates on longitudinal sections of eggs laid by *Chelymorpha gressoria* females (untreated control). Probes used: *Chelymorpha gressoria* host (cyan: 18S rRNA), the DAPI-stained DNA (orange), *Stammera* from *Chelymorpha gressoria* (green: 16SrRNA), and *Stammera* from *Chelymorpha alternans* (magenta: 16S rRNA). Autofluorescence is observed for egg shells and caplets. **(1)** correspond to the merged channels images while the others correspond to individual channel images: **(2)** host probe, **(3)** DAPI-stained DNA, **(4)** *Stammera* from *Chelymorpha gressoria* probe, and **(5)** *Stammera* from *Chelymorpha alternans* probe. Scale bars (50 µm) are included for reference.

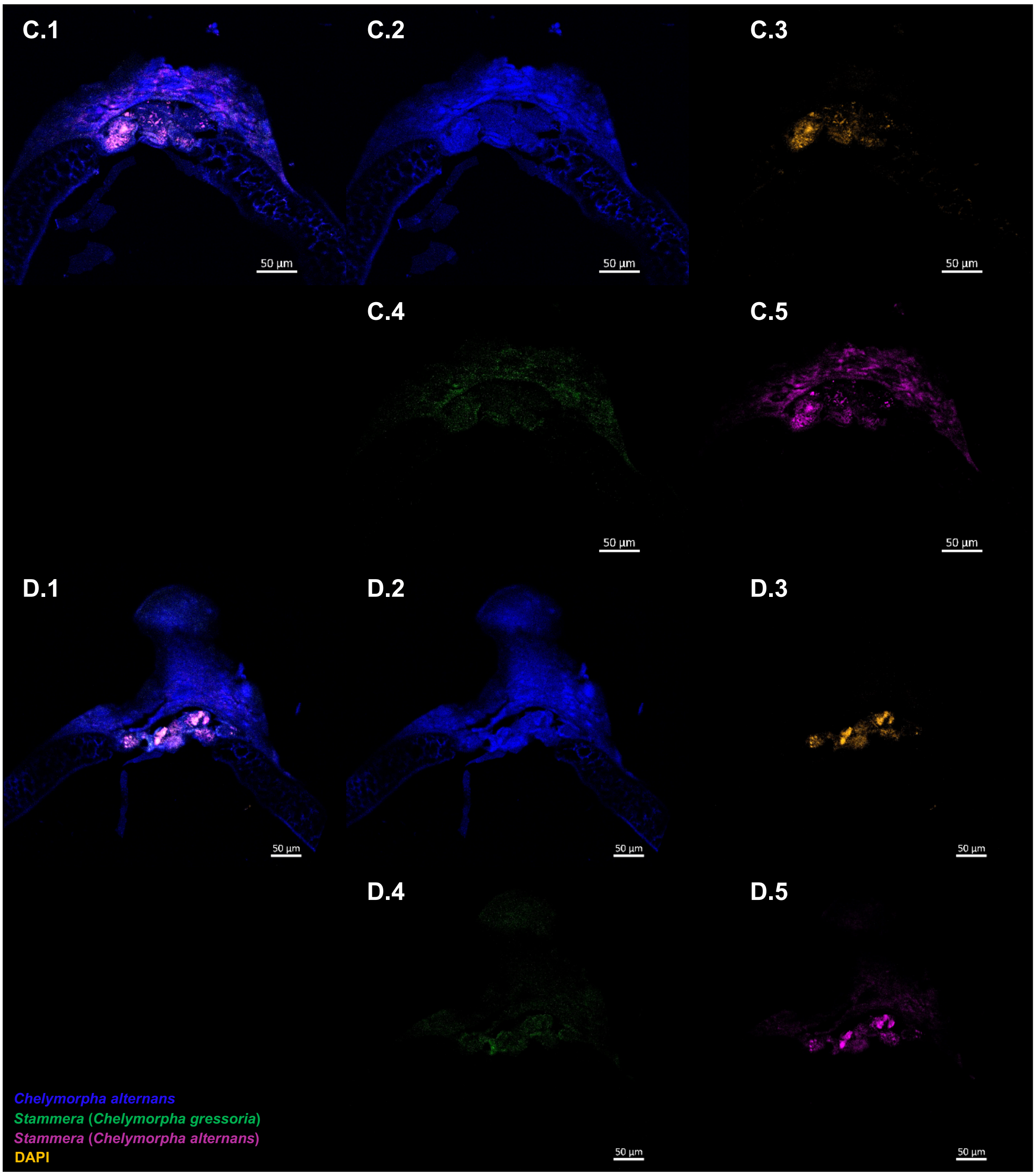


**Figure S7. (C-D)** Fluorescence *in situ* hybridization (FISH) replicates on longitudinal sections of eggs laid by *Chelymorpha alternans* females (untreated control). Probes used: *Chelymorpha alternans* host (blue: 18S rRNA), the DAPI-stained DNA (orange), *Stammera* from *Chelymorpha gressoria* (green: 16SrRNA), and *Stammera* from *Chelymorpha alternans* (magenta: 16S rRNA). Autofluorescence is observed for egg shells and caplets. **(1)** correspond to the merged channels images while the others correspond to individual channel images: **(2)** host probe, **(3)** DAPI-stained DNA, **(4)** *Stammera* from *Chelymorpha gressoria* probe, and **(5)** *Stammera* from *Chelymorpha alternans* probe. Scale bars (50 µm) are included for reference.

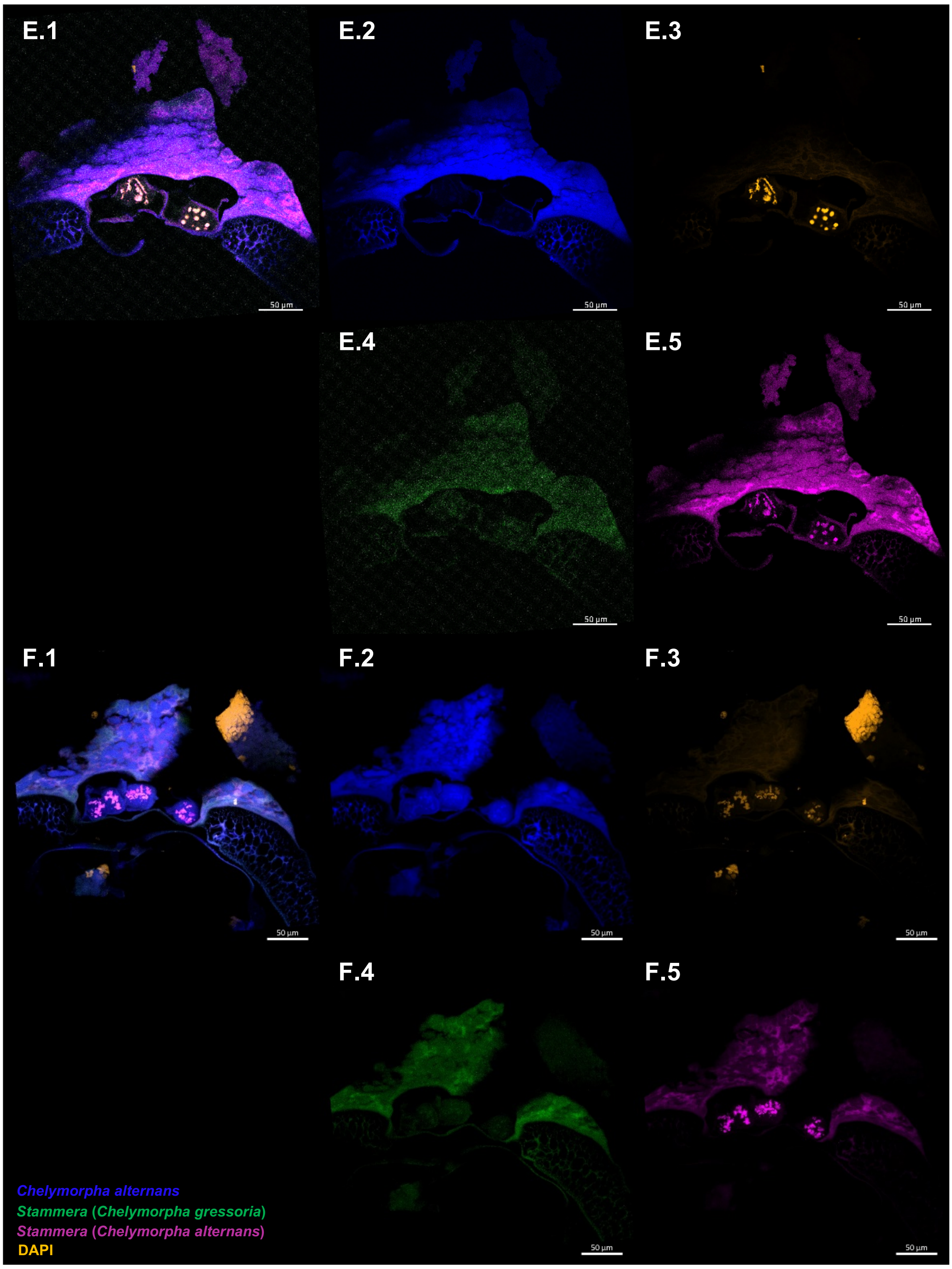


**Figure S7. (E-F)** Fluorescence *in situ* hybridization (FISH) replicates on longitudinal sections of eggs laid by *Chelymorpha alternans* females re-infected by the native symbiont. Probes used: *Chelymorpha alternans* host (blue: 18S rRNA), the DAPI-stained DNA (orange), *Stammera* from *Chelymorpha gressoria* (green: 16SrRNA), and *Stammera* from *Chelymorpha alternans* (magenta: 16S rRNA). Autofluorescence is observed for egg shells and caplets. **(1)** correspond to the merged channels images while the others correspond to individual channel images: **(2)** host probe, **(3)** DAPI-stained DNA, **(4)** *Stammera* from *Chelymorpha gressoria* probe, and **(5)** *Stammera* from *Chelymorpha alternans* probe. Scale bars (50 µm) are included for reference.

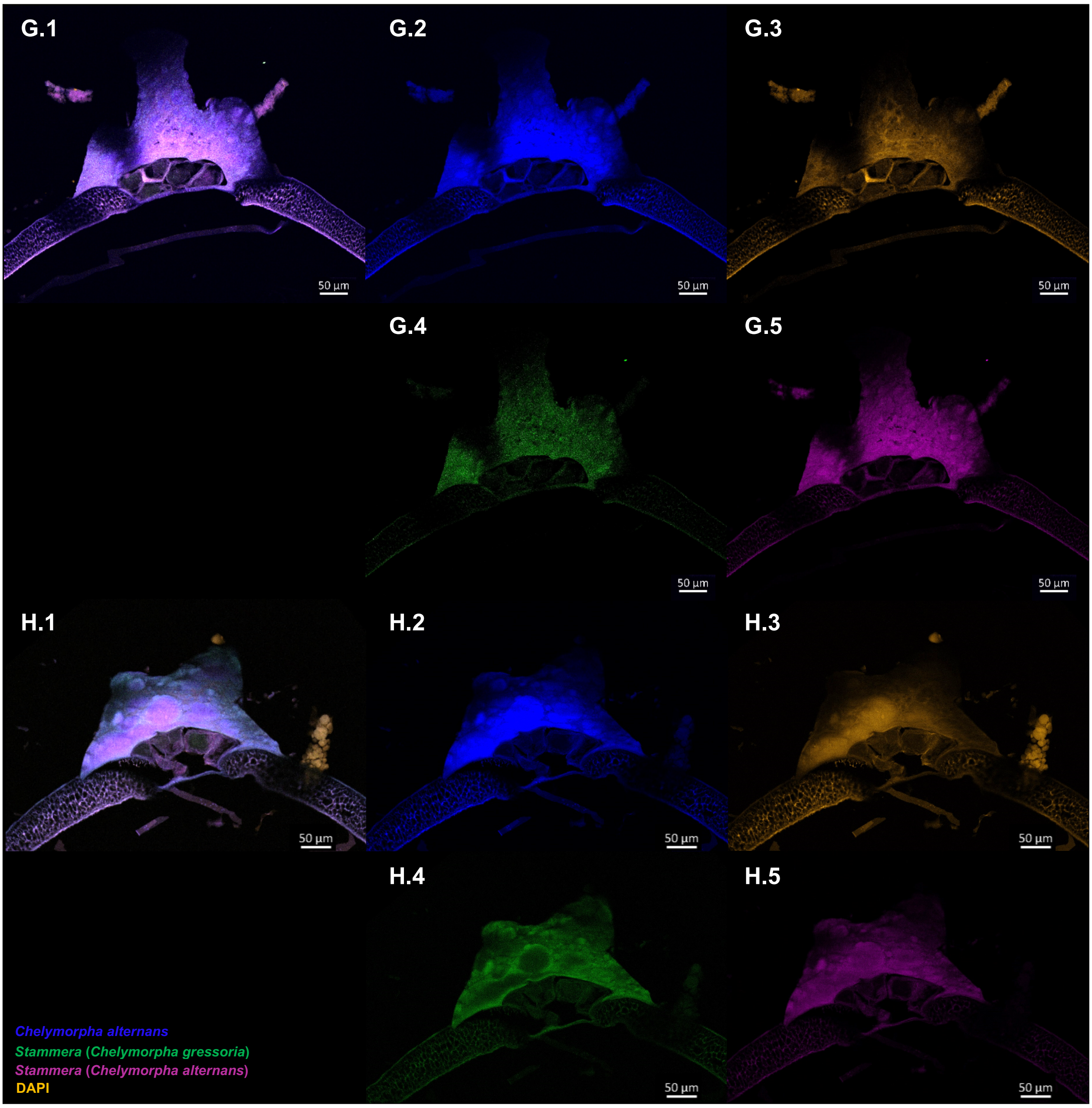
G.1
G.2
G.3
G.4
G.5
H.1
H.2
H.3
H.4
H.5
50 µm
Chelymorpha alternans
Stammera (Chelymorpha gressoria)
Stammera (Chelymorpha alternans)
DAPI

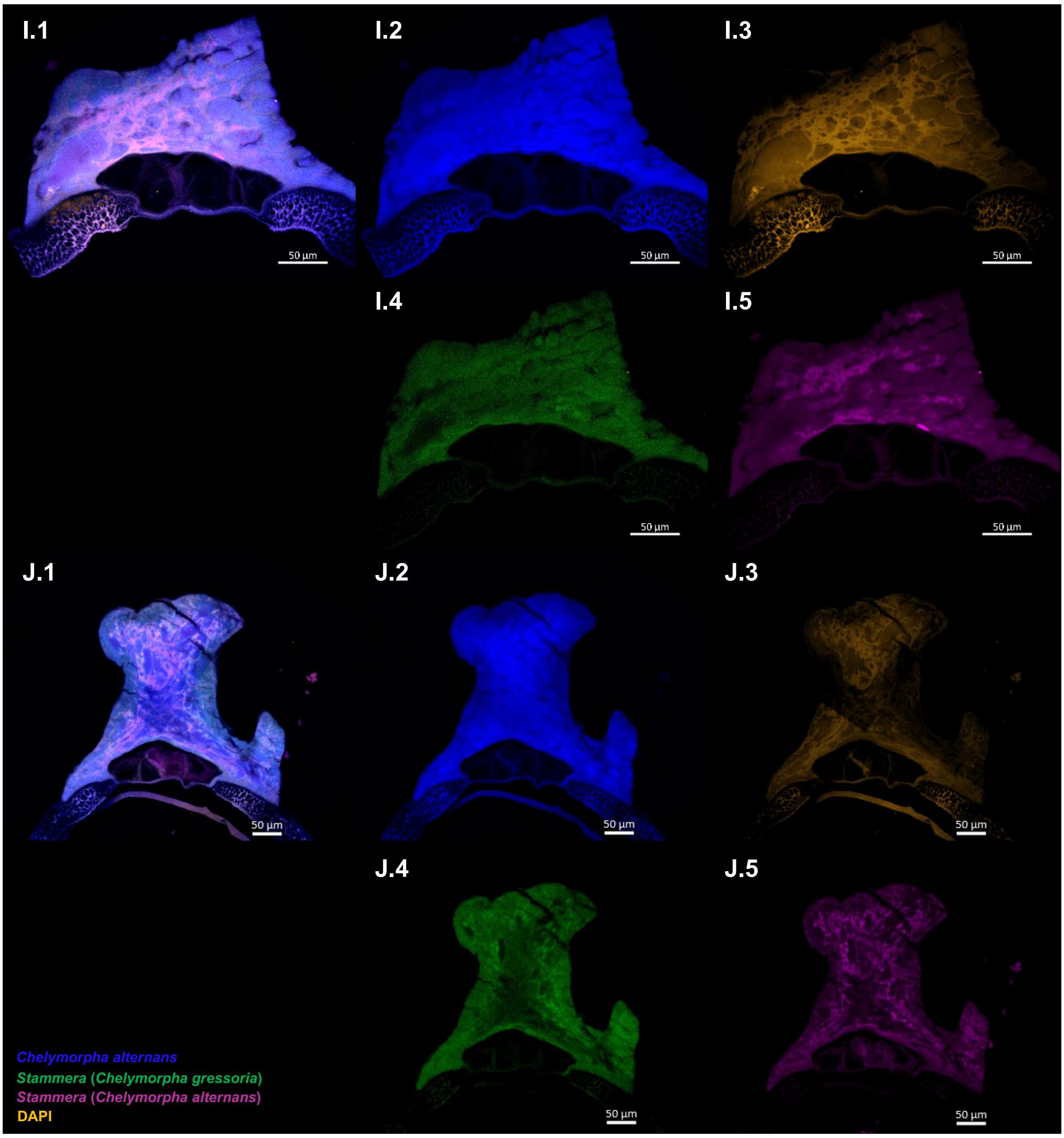
I.1
I.2
I.3
I.4
I.5
J.1
J.2
J.3
J.4
J.5
50 µm
Chelymorpha alternans
Stammera (Chelymorpha gressoria)
Stammera (Chelymorpha alternans)
DAPI

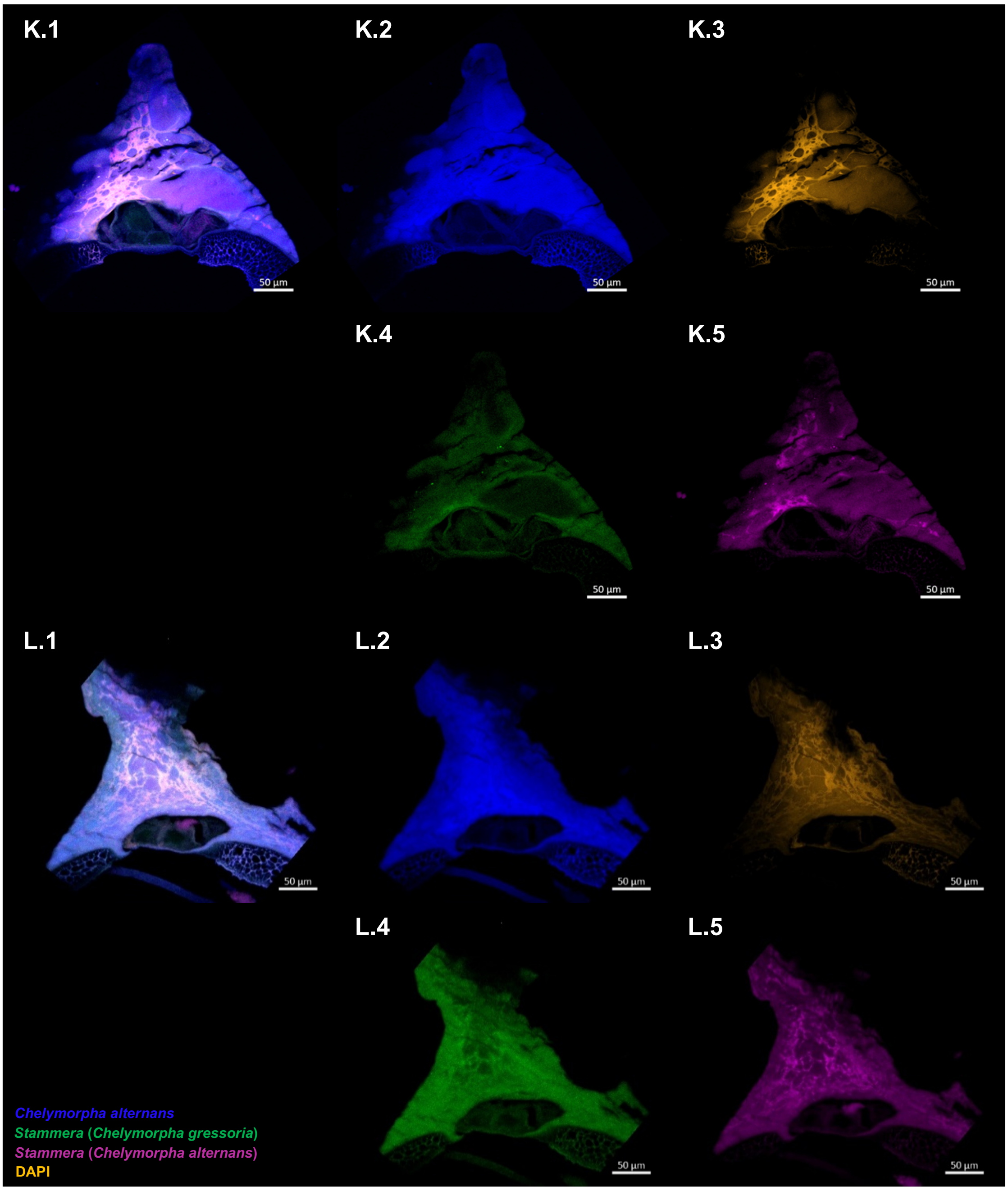


**Figure S7. (G-L)** Fluorescence *in situ* hybridization (FISH) replicates on longitudinal sections of eggs laid by *Chelymorpha alternans* females cross-infected by the non-native symbiont. Probes used: *Chelymorpha alternans* host (blue: 18S rRNA), the DAPI-stained DNA (orange), *Stammera* from *Chelymorpha gressoria* (green: 16SrRNA), and *Stammera* from *Chelymorpha alternans* (magenta: 16S rRNA). Autofluorescence is observed for egg shells and caplets. **(1)** correspond to the merged channels images while the others correspond to individual channel images: **(2)** host probe, **(3)** DAPI-stained DNA, **(4)** *Stammera* from *Chelymorpha gressoria* probe, and **(5)** *Stammera* from *Chelymorpha alternans* probe. **(G-H)** correspond to the first replicate and **(I-L)** correspond to the second replicate. Scale bars (50 µm) are included for reference.

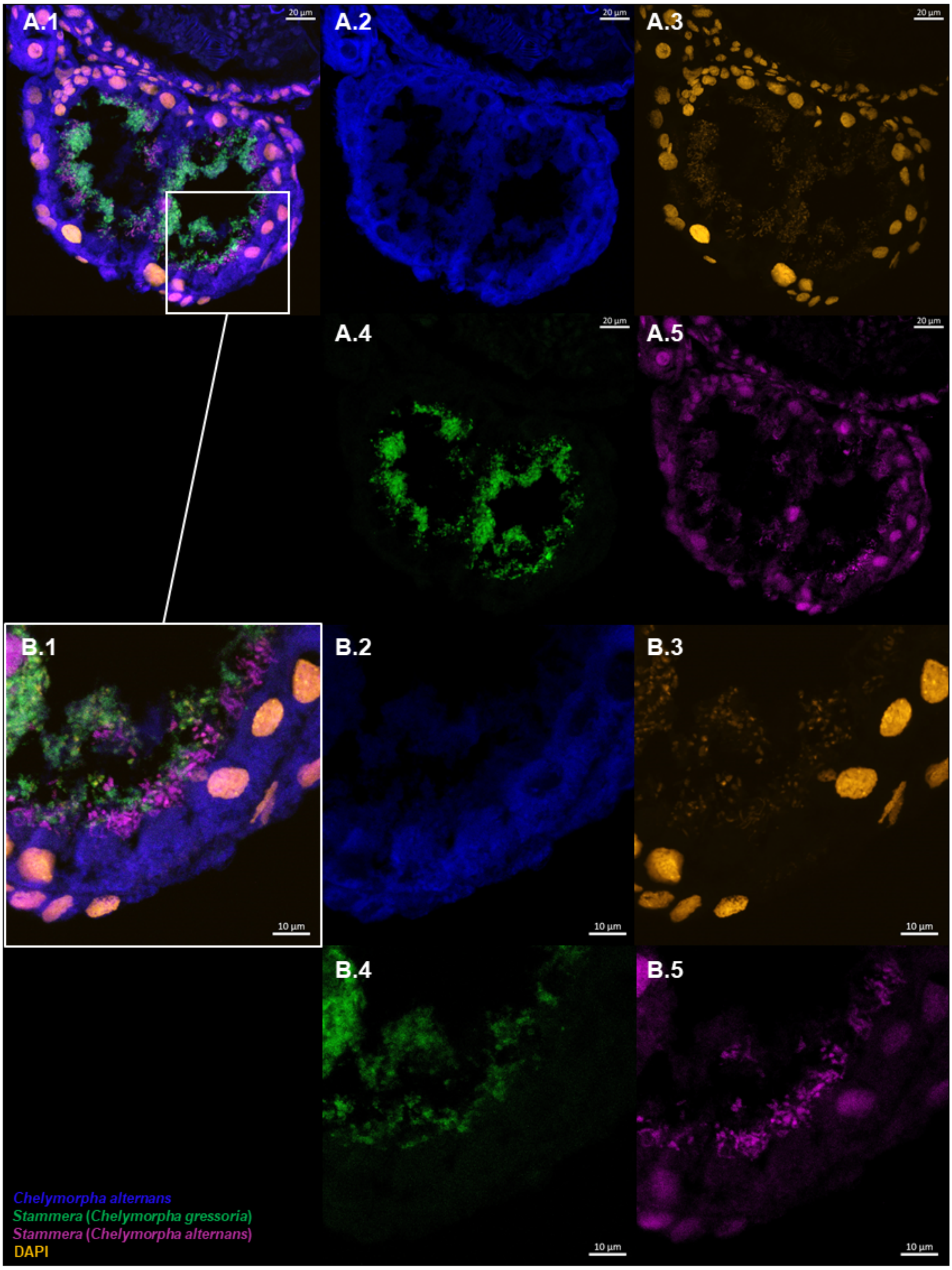


**Figure S8. (A)** Fluorescence *in situ* hybridization (FISH) cross-section of foregut symbiotic organs of 5-day-old, dually infected *Chelymorpha alternans* larvae. **(B)** Close-up view of the foregut symbiotic organ. Probes used: *Chelymorpha alternans* host (blue: 18S rRNA), the DAPI-stained DNA (orange), *Stammera* from *Chelymorpha gressoria* (green: 16SrRNA), and *Stammera* from *Chelymorpha alternans* (magenta: 16S rRNA). **(1)** correspond to the merged channels images while the others correspond to individual channel images: **(2)** *Chelymorpha alternans* host probe, **(3)** DAPI-stained DNA, **(4)** *Stammera* from *Chelymorpha gressoria* probe, and **(5)** *Stammera* from *Chelymorpha alternans* probe. Scale bars are included for reference.

| Host | Gene prediction | Pathway | Category |
|---|---|---|---|
| *Chelymorpha alternans* | Serine--tRNA ligase (serS) | Aminoacyl-tRNA synthetases | Translation, ribosomal structure and biogenesis |
| *Chelymorpha alternans* | Glyceraldehyde-3-phosphate dehydrogenase A (gapA) | Glycolysis | Carbohydrate transport and metabolism |
| *Chelymorpha alternans* | Inner membrane protein YdjM (ydjM) | NA | General function prediction only |
| *Chelymorpha alternans* | Flavodoxin (fldA) | Heme biosynthesis | Energy production and conversion |
| *Chelymorpha alternans* | hypothetical protein | Unknown | Unknown |
| *Chelymorpha gressoria* | Ribosomal RNA large subunit methyltransferase E (rlmE) | 23S rRNA modification | Translation, ribosomal structure and biogenesis |
| *Chelymorpha gressoria* | Serine hydroxymethyltransferase (glyA) | Serine biosynthesis | Amino acid transport and metabolism |
| *Chelymorpha gressoria* | UDP-N-acetylglucosamine 1-carboxyvinyltransferase (murA) | Mureine biosynthesis | Cell wall/membrane/envelope biogenesis |

**Figure S9.** Comparative list of the only eight genes that differ between the symbionts of *Chelymorpha alternans* and *Chelymorpha gressoria*.
